# CRISPR/Cas9-induced double-strand breaks in huntingtin locus lead to CAG repeat contraction through the extensive DNA end resection and homology-mediated repair

**DOI:** 10.1101/2023.11.24.568568

**Authors:** Pawel Sledzinski, Mateusz Nowaczyk, Marianna Karwacka, Marta Olejniczak

## Abstract

Expansion of the CAG/CTG repeats in functionally unrelated genes is a causative factor in many inherited neurodegenerative disorders, including Huntington’s disease (HD), spinocerebellar ataxias (SCAs) and myotonic dystrophy type 1 (DM1). Despite many years of research, the mechanism responsible for repeat instability is unknown, and recent findings indicate the key role of DNA repair in this process. The repair of DSBs induced by genome editing tools results in the shortening of long CAG repeats in yeast models. Understanding this mechanism is the first step to developing a therapeutic strategy based on controlled shortening of repeats. The aim of this study was to characterize Cas9-induced DSB repair products in the endogenous *HTT* locus in human cells and to identify factors affecting the formation of specific types of sequences. The location of the cleavage site and the surrounding sequence influence the outcome of DNA repair. DSBs within CAG repeats result in shortening of the repeats in frame in ∼90% of products. The mechanism of this contraction involves MRE11-CTIP and RAD51 activity and extensive DNA end resection reaching ∼5000 bp. We demonstrated that a DSB located upstream of CAG repeats induces polymerase theta-mediated end joining, resulting in deletion of the entire CAG tract. Furthermore, using unbiased proteomic analysis, we identified novel factors that may be involved in CAG sequence repair.

**Graphical abstract:** 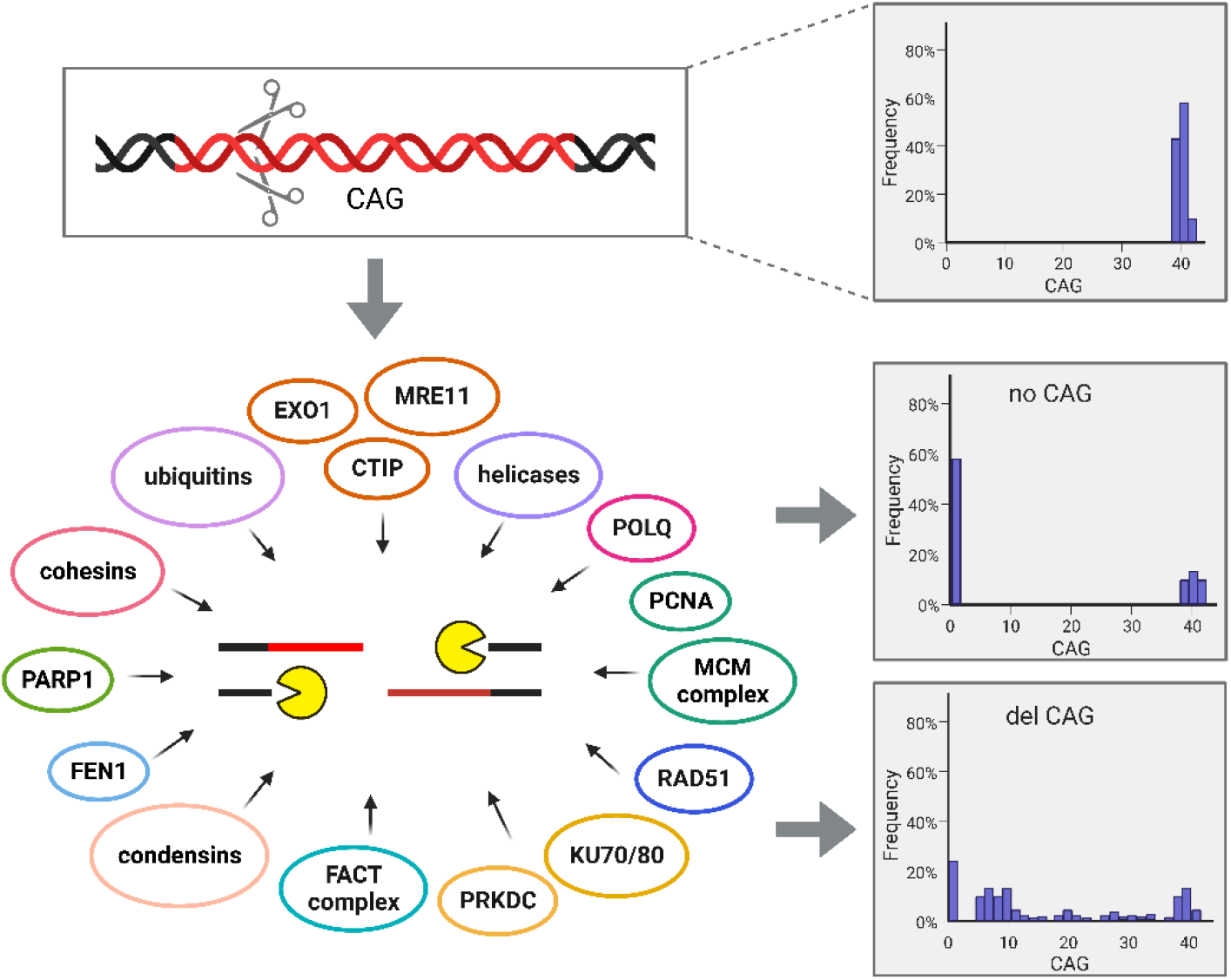

DNA double-strand break repair in the CAG repeat locus engages numerous factors from various repair pathways and depends mainly on DNA end resection, leading to tract shortening.

## INTRODUCTION

Repetitive sequences are abundant in eukaryotic genomes, constituting more than half of the total nuclear DNA content in most species [1]. They are inherently susceptible to length polymorphism that, aside from its proposed roles in genome and protein evolution [2], may also lead to disease [3]. Expanded CNG trinucleotide repeats have been found to contribute to neurodegenerative disorders, e.g., Huntington’s disease (HD), spinocerebellar ataxias (SCAs) or myotonic dystrophy type 1 (DM1) [3]. In case of HD, the disease manifests when the length of the CAG repeats in the first exon of the *HTT* gene reaches 36 or more [4]. It was suggested 20 years ago that double-strand break (DSB) repair is involved in trinucleotide repeat contractions and expansions [5]. Many subsequent studies corroborated this observation [6–11]. Long CAG repeats are often sites of chromosome breakage in yeast when DNA replication is slowed [12]. DSBs can also arise from unligated nicks or collapsed forks as structure-forming strands containing repeats promote DSBs during replication or transcription (reviewed in [3]). Additionally, it has been proposed that DSBs occur in mature neurons not only as a consequence of oxidative stress and transcription errors but also as a natural response to neuronal stimulation (reviewed in [13]).

DSBs in DNA are repaired by classical nonhomologous end joining (NHEJ) or homology-mediated repair, which includes homologous recombination (HR) (reviewed in [14]), single strand annealing (SSA) (reviewed in [15]), microhomology-mediated end joining (MMEJ) and break-induced replication (BIR) (reviewed in [16,17]). All pathways listed above require a DNA end resection process to uncover the regions of homology; however, the degree of resection and homology size are different depending on the repair pathway [18,19].

With the advent of genetic engineering techniques, especially the recent CRISPR−Cas revolution, there have been attempts to use these new tools to explore the molecular mechanism responsible for repeat instability as well as to shorten mutated repeats. Even a small reduction in tract length or in the probability of expansion may have significant implications for the prevention and treatment of repeat expansion diseases. Early studies on this topic revealed many interesting phenomena closely related to the properties of the repeated sequences, e.g., induction of DSBs within CAG or CTG tracts by SpCas9 or ZF nucleases led to both expansions and contractions of the repeats [11,20], whereas single-strand cuts generated by Cas9 nickase resulted in contractions of CAG tracts [6]. In the latter study, the authors suggested that ATR kinase plays role in preventing the tract instability and that the ATM kinase is involved in promoting contractions. They also observed that the instability induced by ATR inhibition is dependent on MSH2- and XPA-related activity [6]. In a yeast model, a break introduced in a CTG tract by the TALEN or SpCas9 endonucleases leads to contraction of repeats below the pathological length [7,8]. In search for the mechanism responsible for the observed phenomenon, the authors found that bidirectional resections take place and that Rad50 is essential for the process, whereas Sae2 (the yeast homologue of CTIP) is required to resect the DNA end containing most of the repetitive tract. Knockout of RAD51, POL32 or DNL4 did not impact the repair process. The authors hypothesized that both the NHEJ and SSA pathways are involved in the process of repeat shortening and that Sae2 plays a role in the removal of secondary structures formed by single DNA strands exposed by resection [7]. Unexpectedly, repair of DSBs generated large chromosomal deletions around the cutting site [8].

Interestingly, even a single DSB in a flanking region of the expanded CGG repeats in the *FMR1* gene and expanded CTG repeats in the DM1 locus was found to be enough to induce the uncontrollable deletion of the entire tract in patient cells [9]. On the other hand, precise excision of the CNG repeats was possible using dual cutting at both sides of the tract [11,21]. The idea of contracting the expanded repeats has also been tested in animal models [22,23].

Despite these interesting observations, the instability of repeats as a result of a DSB-induced DNA repair mechanism is poorly understood. This knowledge is necessary to develop more effective and specific genome editing tools and to predict and control expansions and contractions of repeated sequences.

Here, we try to identify key mechanisms responsible for the shortening of the CAG tract by studying the phenomena taking place after SpCas9 induction of DSBs in the CAG microsatellite region in the *HTT* gene. In contrast to the approach taken in previous work in this field, we used an endogenous model of the human *HTT* gene without altering its genomic context. In addition, we applied techniques allowing an unbiased study of the proteins taking part in the regulation of CAG tract instability.

We showed that the endonuclease cleavage site and the DSB flanking sequence play a key role in the selection of the DNA repair mechanism, resulting in different editing outcomes. DNA end resection is a common step in the repair of the CAG repeat region. Our results suggest that DSBs located upstream of CAG repeats induce polymerase theta (POLθ)-mediated end joining (TMEJ), resulting in deletion of the entire CAG tract. In contrast, DSBs within CAG repeats result in shortening of the repeats in frame through a mechanism dependent on MRE11-CTIP and RAD51 activity. These findings shed new light on the process of DSB repair and microsatellite instability in human cells.

## MATERIALS AND METHODS

### CRISPR−Cas9 system

The guide RNAs targeting exon 1 of the *HTT* gene (HTT_sg1, HTT_sg2, HTT_sg3 and HTT_sg4) have previously been described and validated [21,24]. Depending on the experiment we used a plasmid-based CRISPR−Cas9 system (PX458) or RNP complexes. The RNP complex was prepared according to the IDT Alt-R CRISPR−Cas9 System instructions. Briefly, the crRNA and tracrRNA oligos were mixed in equimolar concentrations to a final duplex concentration of 44 μM (HTT_gRNA1, HTT_gRNA2). The mix was heated at 95 °C for 5 min and incubated at room temperature for 20 min. Then, the guide complex was mixed with Cas9 enzyme (VBCF Protein Technologies facility http://www.vbcf.ac.at) at ratio of 22 pmol to 18 pmol. The mix was incubated at room temperature for 20 min. The sequences of specific oligodeoxynucleotides are listed in Supplementary Table S1.

### Cell culture and transfection

We used previously generated cell lines: HEK293T-41CAG cells (41/41 CAG in the *HTT* gene) and HEK293T-83CAG cells (83/83 CAG in the *HTT* gene) [25]. The cells were grown in Dulbecco’s modified Eagle’s medium (Lonza; Basel, Switzerland) supplemented with 10% foetal bovine serum (Sigma−Aldrich), antibiotics (Sigma−Aldrich) and L-glutamine (Sigma−Aldrich).

Transfections were performed with the Neon^TM^ Transfection System (Invitrogen, Carlsbad, CA, USA). Briefly, 2×10^5^ cells were harvested, resuspended in PBS and electroporated with the RNP complex containing one of the four used gRNAs in 10 μl tips using the following parameters: 1150 V, 20 ms, 2 pulses. As an alternative, cells were transfected with plasmids (described previously [21]) using the Neon^TM^ Transfection System (Invitrogen). A total of 5×10^6^ cells in 10-cm tissue culture plates were transfected with 25 µg of PX458 plasmid containing cloned sgRNA using the electroporation method (1100 V, 20 ms, 2 pulses). Alternatively, the polyethylenimine (PEI) method was used. A total of 4×10^5^ cells were transfected with 2.5 ug of a plasmid complexed with 7.5 ul PEI (1 mg/mL). The method was scaled up if necessary.

### DNA amplification and targeted deep sequencing

Genomic DNA was extracted using the Genomic DNA Isolation Kit (Norgen, Biotek Corp.) according to the manufacturer’s instructions and quantified using a spectrophotometer/fluorometer (DeNovix). The DNA obtained from cell cultures transfected with the RNP complex was amplified by nested PCR using Herculase II Fusion DNA Polymerase (Agilent). The first amplification was performed with primers IG36 and IG37 as follows: initial denaturation at 98 °C for 3 min; 35 cycles at 98 °C for 20 s, 61.1 °C for 20 s, 72 °C for 20 s; final elongation at 72 °C for 3 min. PCR products were purified using the GeneJET PCR Purification Kit (Thermo Fisher), diluted 10X and used as a template for the second amplification performed with primers IG30 and IG31 as follows: initial denaturation at 98 °C for 3 min; 30 cycles at 98°C for 20 s, 56 °C for 20 s, 72 °C for 12 s; final elongation at 72 °C. Sequences of specific primers are listed in Supplementary Table S1. The length of the uncleaved product was 242 bp. The PCR products were used for library generation and next-generation sequencing to analyse the pattern of CAG tract shortening upon Cas9 cleavage. The library preparation and sequencing procedures were performed by Novogene (UK). Sequencing libraries were generated using the NEBNext Ultra II DNA Library Prep Kit for Illumina (New England Biolabs, UK) following manufactures’ protocol. Libraries were sequenced using the Illumina NovaSeq6000 platform with the following settings: 250 bp paired end reads and 1 million reads per sample. The obtained data are deposited at Sequence Read Archive (SRA, accession number: PRJNA1006315).

### Bioinformatic analysis

Analysis was performed by ideas4biology Ltd. Briefly: the overall quality of the data was tested with FastQC version 0.11.5. Adapters were removed using standard adapter sequence set from bbduk2 package. Reads with quality under 5 were discarded. The reads were merged together using bbmerge script from BBMap package. After merging, now single end reads were mapped using BBMap global aligner against supplied sequence. Mapping quality was assesed using qualimap v.2.2.2-dev. Aligned reads were filtered using -F260 and -q 30 flags. Sequences were clustered using cd-hit application. Only sequences of the same lengths and a minimum of 95% sequence similarity were merged into clusters. . bam files containing unique sets of reads were converted to pairwise alignments using sam2pairwise program. Data were supplied to Tandem Repeats Finder v409. Recommended parameters were used with one change in Minscore parameter set to 10 to allow shorter hits. Results were filtered for repeats containing only combinations of C,A,G letters. Reads with a given length of “CAG” string were counted and a ratio between number of reads containing given “CAG” length and all reads was calculated.

### Chromatin immunoprecipitation

A total of 5×10^6^ of cells were transfected with 25 µg of PX458 plasmid containing cloned HTT_sgRNA1 or HTT_sgRNA2 using the electroporation method as described above. The cells were then collected at 5 time points (0 h, 12 h, 24 h, 48 h, 72 h) in the following way.

Formaldehyde (Thermo Scientific) at final concentration of 1% was used to cross-link the cell chromatin. The reaction was stopped by using glycine (BioShop) at final concentration of 1.11 M. Cells were then collected by scraping in cold PBS with 1 mM PMSF. After centrifugation, the cells were lysed in RIPA buffer (50 mM Tris-HCl pH 8, 150 mM NaCl, 2 mM EDTA pH 8, 1% Triton X-100, 0.5% sodium deoxycholate, 0.2% SDS) with a protease inhibitor cocktail (PIC, BioShop) on ice. The chromatin was sonicated using a BioRuptor Pico sonicator (Diagenode; 5 cycles, 30 s ON, 30 s OFF, the average length of fragments was 400 bp). Finally, the lysates were centrifuged (12500xg, 4 °C, 5 min) and the supernatants were transferred to new tubes. Lysates were suspended in 710 µl of RIPA buffer with 7.5 µl PIC and incubated with gentle mixing overnight at 4 °C with 20 µl of Dynabeads^TM^ Protein G (Invitrogen) coated with antibody against the analysed protein (listed in Supplementary Table S2). Next, the beads were separated on a magnetic separator and washed 6 times with RIPA buffer and 1 time with TBS (Pierce). Elution was performed by incubation of the beads in elution buffer (50 mM Tris-HCl pH 8, 10 mM EDTA pH 8, 1% SDS) at 65 °C for 5 h. The eluate was transferred to a new tube containing 100 μl of TE buffer and then incubated with proteinase K (Diagenode) at 55 °C for 1 h. The DNA was purified with a GeneJET PCR Purification Kit (Thermo Scientific). As a normalization of the chromatin concentration between samples, the input was prepared as described above, but the incubation with magnetic beads was omitted.

### ChIP−qPCR

For input and eluate samples collected at 5 time points, quantitative PCR was performed using a CFX Connect Real-Time PCR Detection System (Bio-Rad, Hercules, CA) and SsoAdvanced Universal SYBR Green Supermix (Bio-Rad). The left and right flanking regions of the CAG tract in the *HTT* gene were amplified under the following thermal cycling conditions: denaturation at 98 °C for 3 min followed by 50 cycles of denaturation at 95 °C for 15 s and annealing at 66.2°C for 30 s and elongation at 72°C for 30 s. As the reference gene, β-actin was amplified under the following thermal cycling conditions: denaturation at 95°C for 30 s followed by 40 cycles of denaturation at 95°C for 15 s and annealing at 60°C for 30 s. Primer sequences are listed in Supplementary Table S1. The value of input recovery was determined on the basis of the average quantification cycle (Cq) for reactions carried out for the input and eluate. The ratio between input recovery for the regions flanking the CAG tract in the *HTT* gene and for the reference gene (β-actin) was used to calculate the fold enrichment.

### DNA:RNA Immunoprecipitation

A total of 5×10^6^ of cells were transfected with 25 µg of PX458 plasmid containing cloned HTT_sgRNA1 or HTT_sgRNA2 using the electroporation method. The cells were then collected after 12 h and lysates were prepared as described above for ChIP. Lysates were digested with proteinase K (Norgen Biotek Corp.) and decrosslinked at 65 °C overnight. Later, DNA containing DNA:RNA hybrids was isolated by phenol:chloroform extraction and precipitated with isopropanol (1:1 v/v) with 3 M sodium acetate (1:10 v/v). Six micrograms of isolated DNA was left as an input and 6 µg of DNA was used for DRIP. The DNA was incubated in IP buffer (50 mM Hepes/KOH at pH 7.5; 0.14 M NaCl; 5 mM EDTA; 1% Triton X-100; 0.1% Na-Deoxycholate) with 20 µl of Dynabeads^TM^ Protein A (Invitrogen) coated with 10 µl of S9.6 antibody (Sigma Aldrich, listed in Supplementary Table 2) at 4 °C with gentle mixing overnight. Next, the beads were separated on a magnetic separator and washed once with IP buffer for 1 min at 4 °C, high salt buffer (50 mM HEPES/KOH pH 7.5, 0.5 M NaCl, 5 mM EDTA pH 8, 1% Triton X-100, 0.1% Na-deoxycholate) for 1 min at 4 °C, wash buffer (10 mM Tris-HCl pH 8, 0.25 M LiCl, 0.5% NP- 40, 0.5% Na-Deoxycholate, 1 mM EDTA pH 8) for 1 min at 4 °C and washed twice with TE buffer (50 mM Tris-HCl pH 8, 10 mM EDTA, 1% SDS) for 1 min at 4°C. Elution from beads was performed by incubation in elution buffer (50 mM Tris-HCl pH 8, 10 mM EDTA pH 8, 1% SDS) at 65 °C for 45 min. The remaining antibodies were digested with proteinase K for 1 h at 55°C. The DNA was then purified by phenol:chloroform extraction and precipitated with isopropanol with sodium acetate as described above.

### DRIP-qPCR

For eluate and 100-fold diluted input, quantitative PCR was performed using a CFX Connect Real-Time PCR Detection System (Bio-Rad, Hercules, CA) and SsoAdvanced Universal SYBR Green Supermix (Bio-Rad). The left and right flanking regions of the CAG tract in the *HTT* gene as well as the positive control locus (*LOC440704*) and negative control locus (*ZNF554*) were amplified under the following thermal cycling conditions: denaturation at 98 °C for 3 min, followed by 40 cycles of denaturation at 95 °C for 15 s, annealing at 66.2 °C for 30 s and elongation at 72 °C for 30 s. An additional positive locus (*ING3*) was amplified under the following thermal cycling conditions: denaturation at 95 °C for 30 s, followed by 40 cycles of denaturation at 95 °C for 15 s and annealing at 60 °C for 30 s. Primer sequences are listed in Supplementary Table S1. The value of input recovery was determined on the basis of the average Cq for reactions carried out for the input and eluate. The ratio between input recovery for the regions flanking the CAG tract in the *HTT* gene or positive control locus and the negative control locus (*ZNF554*) was used to calculate the fold enrichment.

### Immunofluorescence

HEK293T, HEK293T-41CAG cells and HEK293T-83CAG cells were seeded on 12-well plates with glass coverslips coated with Geltrex (Thermo Fisher) 3 h prior to transfection. Cells were transfected with RNP complexes containing HTT_gRNA1 or HTT_gRNA2 using Lipofectamine RNAiMAX (Invitrogen). After 3 h, the cells were fixed with 4% paraformaldehyde for 30 min at RT. The fixed samples were washed once with PBS. The cells were permeabilized with 0.5% Triton X-100 in PBS for 10 min and blocked in 1% BSA with 0.2% Triton X-100 in PBS for 1 h at RT. The samples were washed with PBS 3 times. Coverslips were incubated overnight at 4 °C with the primary antibodies anti-DNA:RNA hybrids and anti-γH2AX (listed in Supplementary Table S2) diluted in 1% BSA with 0.2% Triton X-100 in PBS. The next day, the coverslips were washed three times with PBS and incubated with anti-mouse and anti-rabbit antibodies labelled with Alexa488 and Alexa647, respectively (1:500, listed in Supplementary Table S2), for 1 h at RT. After rinsing, the coverslips were mounted onto glass with SlowFade Diamond Antifade Mountant with DAPI (Life Technologies). Images were captured with a Leica SP5 confocal microscope under 63x magnification. Leica Application Suite X (3.3.3) was used to analyse colocalization (at least 70 spots per cell, for 15 cells of each cell type) and fluorescence intensity (16 nuclei from cells of each cell type). Colocalization of DNA:RNA hybrids and γH2AX was determined based on Pearson’s coefficient values, and the fluorescence intensity was measured as a ratio between signals from DNA:RNA hybrids and γH2AX histones.

### Resection analysis

Cells were electroporated with RNP complexes containing HTT_gRNA1 or HTT_gRNA2. After electroporation, the cells were cultivated in complete DMEM with 20% FBS and harvested for DNA isolation at 1 h, 6 h, 24 h and 48 h post transfection. The DNA was isolated using a Genomic DNA Isolation Kit (NORGEN Biotek) according to the manufacturer’s protocol. Each sample was divided into two parts: one was incubated overnight with restriction enzymes, and the other was incubated without restriction enzymes (see Supplementrary Table S3). After incubation, DNA was used in qPCR with primers that flank the restriction sites in separate reactions for each pair of primers (see Supplementary Table S1). Each PCR was performed in three technical replicates, and Cq results were used to obtain the Cq mean and to calculate the level of resection for each timepoint and site from A to H using the formula presented in [26]. Each experiment was conducted in three biological replicates.

### Inhibition of DNA repair proteins

HEK293T-41CAG cells (3×10^5^ cells per well) were cultivated in 6-well plates in complete DMEM. One day after passage, the medium was changed to complete DMEM containing a particular inhibitor (see Supplementary Table S4). After 24 h of incubation, cells were transfected with PEI-plasmid complexes: 2.5 µg of PX458 plasmid containing cloned HTT_sgRNA1 or HTT_sgRNA2 and 7.5 ul of PEI for each well. After 4 h, the medium was changed again to DMEM containing specific inhibitor at the same concentration as previously described. Twenty-four hours post transfection, cells were harvested and sorted on a FACSAria Fusion cell sorter (Becton Dickinson) in The Laboratory of Single Cell Analyses IBCH PAS.

DNA from GFP-positive cells was isolated using QuickExtract DNA Extraction Solution (Lucigen) according to the manufacturer’s protocol. Isolated DNA was used in PCR with IG36 and IG37 primers, with the forward primer linked with fluorescein on its 5’ end. Q5 polymerase (NEB) was used for the amplification of the edited CAG locus in HTT as follows: initial denaturation at 98°C for 3 min; 35 cycles at 98 °C for 10 s, 68 °C for 30 s and 72 °C for 5 s; and final elongation at 72 °C for 2 min. The PCR products were purified using the Gene JET PCR Purification Kit (Thermo Fisher) according to the manufacturer’s protocol. Capillary electrophoresis was conducted in the Molecular Biology Techniques Laboratory at Adam Mickiewicz University.

### siRNA knockdown of DNA repair proteins

A total of 2×10^5^ HEK293T-41CAG cells per well were used in reverse transfection using a particular siRNA (see Supplementary Table S4). siRNA reverse transfection was performed as follows: 5 ul of Lipofectamine2000 (Thermo Fisher) was incubated at room temperature with siRNA for 20 min in 500 ul OptiMEM (Gibco) in each well of a 6-well plate. A total of 2×10^5^ cells in 2 ml of complete DMEM were added to lipofectamine-siRNA complexes. The final concentration of siRNA was 50-100 nM (Supplementary Table S4). Seventy-two hours post transfection, cells were transfected with PEI-plasmid complexes: 2.5 µg of PX458 plasmid containing cloned HTT_sgRNA1 or HTT_sgRNA2 and 7.5 ul of PEI for each well. Twenty-four hours post transfection, cells were harvested, sorted and treated according to the aforementioned methods described in the “Inhibition of DNA repair proteins” section. The experiment was conducted in three biological replicates.

### Analysis of the PCR products in capillary electrophoresis

Capillary electrophoresis was performed on an ABI Prism 3130xl (Applied Biosystems) with POP-7 polymer using a G5 filter. One microliter of sample was diluted in 9 ul Hi-Di™ Formamide (Applied Biosystems) containing 0.25 ul GeneScan™ 600 LIZ™ Size Standard. Before resolution, samples were denatured at 95 °C for 5 min and cooled to 10 °C. The results were analysed with Peak Scanner Software V1.0 (Applied Biosystems). Each peak was described by the following parameters: Size-indicating the peak size based on base pairs; Height-indicating the peak height; Area in BP- indicating the peak area based on base pairs. To cut off the background for each sample, every peak in which the Height parameter was below 5% of the value of this parameter related to the highest peak in a particular sample was excluded from the calculation. Then, the values of Area in BP for each peak were added as well as the values of Area in BP for peaks corresponding only to PCR products containing 41 CAG repeats to calculate the percentage of shortened and unshortened products. The mean percentage for treated and control samples was calculated using values from three biological repeats. Statistical significance was determined using an unpaired t test.

### enChIP

#### HEK293T-dCas9-3xFLAG cell generation

gRNA targeting a site 208 bp upstream from the CAG tract was inserted into the 3xFLAG- dCas9-HA-2xNLS (Addgene) plasmid by digesting the plasmid with BsmBI (Esp3I) and ligating it to annealed gRNA oligos (see Supplementary Table S1). For lentivirus production, the plasmid was cotransfected with the packaging plasmids pPACKH1-GAG, pPACKH1-REV, and pVSVG (System Biosciences) into HEK293TN cells. The medium was collected on Days 2 and 3, and the viral supernatants were passed through 0.45-μm filters and concentrated using PEGit Virus Precipitation Solution (System Biosciences). The lentiviral vectors were resuspended in Opti-MEM (GIBCO, Invitrogen, Carlsbad, CA). To generate the HEK293T-dCas9-3xFLAG transgenic cell line, cells were transduced with lentivirus in the presence of 8 μg/ml polybrene and selected using 1.5 μg/ml puromycin. Monoclonal transgenic lines were generated by transduction at a low cell density using a low multiplicity of infection (MOI <1) and allowing cells that survived selection to form colonies before individual clones were isolated

#### Protein extraction and immunoblotting

Total protein extracts were generated by lysing cells in RIPA buffer. A total of 30 μg of protein was diluted in sample buffer containing 2-mercaptoethanol and boiled for 5 min. The samples were run on a 12% Tris-acetate SDS-polyacrylamide gel and transferred to a nitrocellulose membrane (Sigma−Aldrich). Membranes were blocked in TBS containing 0.1% Tween-20% and 5% nonfat dry milk for 2 h at room temperature. The primary and secondary antibodies were used in TBS/0.1% Tween 20 buffer containing 5% nonfat milk. Immunoreactions were detected using Western Bright Quantum HRP Substrate (Advansta, Menlo Park, CA).

#### enChIP−qPCR

The procedure was adapted with modifications from Fujita et al. [27]. Briefly, 5×10^6^ HEK293T- dCas9-3xFLAG cells in 10-cm tissue culture plates were transfected with 25 µg of PX458 plasmid containing cloned HTT_sgRNA1 or HTT_sgRNA2 using the PEI transfection method. After 24 h, the cells were washed 4 times with cold PBS and then collected by scraping in cold PBS containing 1 mM PMSF. After centrifugation, the cells were lysed in RIPA buffer with a protease inhibitor cocktail (BioShop) on ice, and the chromatin was sonicated using a BioRuptor Pico sonicator (Diagenode; 5 cycles, 30 s ON, 30 s OFF, the average length of fragments was 400 bp). Then, the cells were centrifuged (12500xg, 4 °C) and the supernatants were transferred to new tubes and incubated overnight at 4 °C with 15 µl of Anti-FLAG® M2 Magnetic Beads (Merck) with gentle mixing. Next, the beads were separated on a magnetic separator and washed 6 times with RIPA buffer and 6 times with TBS (Pierce). Elution was performed by incubation of the beads with 3xFLAG peptide (Sigma Aldrich) at 37 °C for 30 min. Then, the eluate was incubated with RNase in 37 °C for 1 h, and DNA was purified with a GeneJET PCR Purification Kit (Thermo Scientific).

Quantitative PCR was performed in a CFX Connect Real-Time PCR Detection System (Bio-Rad, Hercules, CA) using SsoAdvanced Universal SYBR Green Supermix (Bio-Rad) with β-actin as the reference gene under the following thermal cycling conditions: denaturation at 95 °C for 30 s followed by 40 cycles of denaturation at 95 °C for 15 s and annealing at 60 °C for 30 s. Primer sequences are listed in Supplementary Table S1.

#### enChIP-MS

The procedure was performed as described above until the final wash (6x RIPA, 6x TBS). Then, mass spectrometry was performed by the Mass Spectrometry Laboratory, Institute of Biochemistry and Biophysics, Polish Academy of Sciences. The analysis was conducted in 5 replicates. First, the cysteines were reduced by 1 h incubation with 20 mM tris(2- carboxyethyl)phosphine (TCEP) at 37 °C followed by 10 min incubation at room temperature with 50 mM methyl methanethiosulfonate (MMTS). Digestion was performed at 37 °C overnight with 1 µg of trypsin (Promega). After digestion, the peptides were eluted from the beads by using a magnet. The pulled aliquots were dried in a SpeedVac and resuspended in an extraction buffer (0.1% TFA 2% acetonitrile) with sonication. The next step was processed using single-pot solid-phase-enhanced sample preparation (SP3). The magnetic bead mixes were prepared by combining equal parts of Sera-Mag Carboxyl hydrophilic and hydrophobic particles (09-981-121 and 09-981-123, GE Healthcare). The bead mix was washed three times with MS-grade water and resuspended in a working concentration of 10 µg/µl. The bead mix was then added to the samples and suspended in 100% acetonitrile, and this step was repeated 3 times. Pure peptides were eluted from the beads by using 2% acetonitrile in MS-grade water. Using a magnet, the peptide solution was separated from the beads. The peptide mixture was dried in a SpeedVac and resuspended in 80 µl of extraction buffer (0.1% TFA and 2% acetonitrile) with sonication. Twenty microlitres of each sample was measured on an Orbitrap Exploris 480 mass spectrometer (Thermo Scientific) coupled to an Evosep One chromatograph (Evosep Biognosys). The samples were loaded onto disposable Evotips C18 trap columns (Evosep Biosystems) according to the manufacturer’s protocol with minor modifications as described previously [28]. Chromatography was carried out at a flow rate of 250 nl/min using an 88 min (15 samples per day) gradient on an EV1106 analytical column (Dr Maisch C18 AQ, 1.9 µm beads, 150 µm ID, 15 cm long, Evosep Biosystems). The resulting raw data were processed with MaxQuant software (version 2.1.1) to obtain the LFQ intensities and protein identifications using the *Homo sapiens* reference proteome from UniProt (one protein per gene) supplemented with the Cas9 protein sequence (20595 sequences) with contaminants included. Further analysis was performed in the Perseus suite (version 1.6.15). Proteins with fewer than 3 valid values in both groups were not included in the statistical analysis. The data were log transformed, the missing values were imputed from a normal distribution (width 0.3, downshift 1.8), and a two-sample t test was performed to estimate the statistical significance. The mass spectrometry proteomics data have been deposited to the ProteomeXchange Consortium via the PRIDE [29] partner repository with the dataset identifier PXD044960 and 10.6019/PXD044960

## RESULTS

### DSB location affects CAG repeat region editing results

Since both the length of the CAG repeat sequence and its location in the genome can influence DNA repair mechanisms, in our study, we used an endogenous *HTT* locus with a repeat sequence length characteristic of HD patients. As a model, we used the previously generated HEK293T cell line with 41 CAG repeats in both alleles of the *HTT* gene [25]. Although the HD patients are heterozygous, a homozygous model is more useful in studying the changes in the CAG repeat length. DSBs were induced by SpCas9 and gRNAs described previously [21,24], which utilize PAM (NGG) sequences located in regions flanking CAG repeats (HTT_sgRNA1 and HTT_sgRNA4) or directly targeting CAG repeats (HTT_sgRNA2) (Figure 1A). HTT_sgRNA2 was designed to use a noncanonical PAM sequence (NAG). HEK293T-41CAG cells were transfected with RNP complexes containing both the wtCas9 protein and HTT_gRNA. Genomic DNA was isolated 24 h post transfection. The region of interest was amplified by nested PCR and subjected to deep sequencing. We observed a shortening of the CAG tract both in agarose gel (Figure 1B) and in the NGS results (Figure 1C – 1I). The maximum read length of ∼250 bp limited the possibility of assessing the occurrence of long expansions but gel electrophoresis of the edited sequence did not expose any expansions (Figure 1B). The short, exact (CAG)n indels observed in unedited samples are most likely stutter products generated during repetitive sequence amplification [30] (Figure 1E). This phenomenon was included in the outcome analysis.

**Figure 1.**
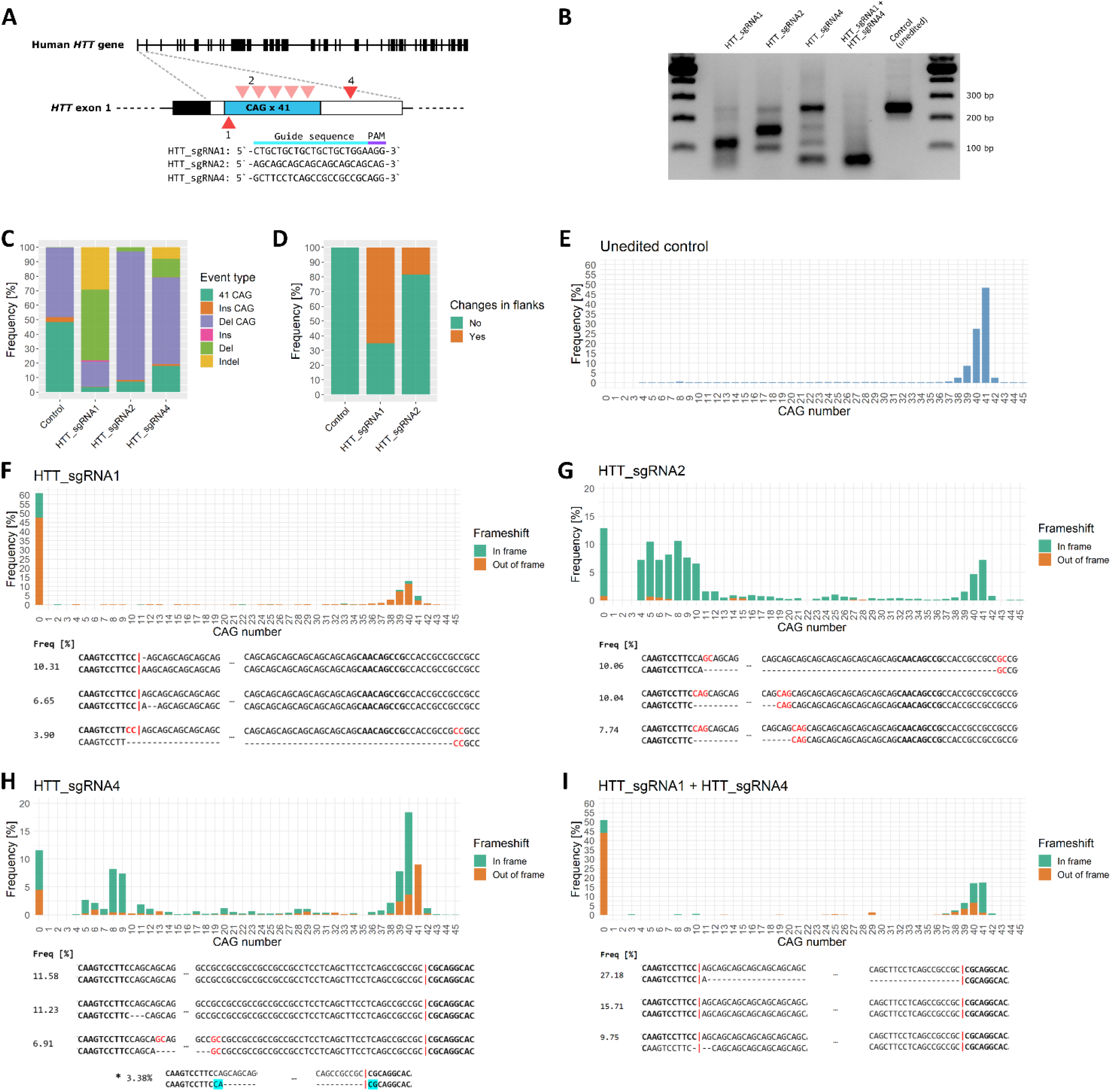
Deep sequencing analysis of CAG repetitive tract shortening induced by CRISPR-Cas9 cleavage. A) Schematic depiction of the CAG repetitive tract in the *HTT* gene with the editing sites marked in red. B) Gel electrophoresis of DNA after editing. Different patterns of shortening are generated by each gRNA used. C) CRISPR-Cas9 editing outcomes divided into six main categories and presented as a barplot comparing their proportions: 41 CAG - unedited original sequence, Ins CAG - increase in the number of CAG triplets (no frameshift), Del CAG - decrease in the number of CAG triplets (no frameshift), Ins - insertions leading to frameshift, Del - deletions leading to frameshift, Indel - composite changes leading to frameshift. D) CRISPR-Cas9 editing outcomes presented as proportions of edits involving changes in the regions flanking the CAG tract and changes in the tract only. We did not include results of HTT_sgRNA4 and HTT_sgRNA1+ HTT_sgRNA4 editing, as these variants target the flank itself. E – I) CAG tract length distribution among reads presented as the number of triplets and the presence of frameshifts. The three most frequent results are presented below the respective graphs. The microhomologous regions playing the role in the tract contraction are marked with red. ‘*’ in H denotes an excision of the entire tract, from ‘G’ of the first CAG triplet to the HTT_sgRNA4 cutting site.

We divided the NGS results into six main categories: “41 CAG” denoting unedited original sequence, “Ins CAG” and “Del CAG” meaning increase and decrease in the number of CAG triplets, respectively (these changes were “clean”, no frameshift occurred), “Ins” and “Del” denoting insertions and deletions leading to frameshift, and “Indel”, a category encompassing all composite changes leading to frameshift.

After introduction of DSBs using HTT_sgRNA1, tract shortening was efficient, as the full-length 41 CAG tract was present in only 4.87% of the reads (Figure 1C). 79% of edits resulted in a frameshift. We observed dominance of deletions (66%), including out-of-frame Del (49%) and “clean”, in-frame Del CAG (17%). The Indel group composed of combined outcomes constituted 28% (Figure 1C). The fraction of edits leading to changes in the flanks constituted 65% (Figure 1D). The single DSB at the beginning of the tract introduced by HTT_sgRNA1 was sufficient to induce robust excision of the entire CAG tract, as indicated by 61% of the reads (Figure 1F). It was, however, not a ‘clean’ excision, as in most cases, the deletion additionally spanned several nucleotides of the flanking regions at both sides of the CAG repeats. Interestingly, the single most abundant editing result was insertion of a single A nucleotide, which occurred in 10.31% of the reads (Figure 1F). This outcome can also be interpreted as a result of Cas9 cleavage in a ‘staggered’ fashion [31]. It occurs when the RuvC domain cuts the nontargeted strand after the 4^th^ nucleotide upstream of the PAM, which may subsequently lead to a duplication of the 4^th^ nucleotide. A substantial group of reads comprises sequences resulting from the deletion of the CAG repeat region between short microhomologous sequences, most frequently CC:GG (see Supplementary File 1). Additionally, a portion of the products contained duplicated sequences from the upstream flank of the CAG repeats, potentially indicating the activity of polymerase theta [32,33].

In the case of HTT_sgRNA2, tract shortening was also efficient, since the full-length tract constituted 7% of the reads (Figure 1C). The majority of the edit outcomes were “clean” in frame Del CAG (89%). The out-of-frame Del was identified in only 3% of the reads. Accordingly, the proportion of outcomes containing changes in flanks was low (18%, Figure 1D). Considerable CAG tract shortening was observed, but contrary to HTT_sgRNA1, the results were more variable in terms of the tract length distribution (Figure 1G). Deletions of the entire CAG region comprised 13% of the sequencing reads, and this group included the most frequent variant (∼10%). Contrary to the edit using HTT_sgRNA1, excisions of only parts of the repetitive tract were frequent. Within this group, sequences with 4-10 CAG repeats predominate. Of note, editing with HTT_sgRNA2 resulted in a substantially reduced incidence of frameshifts (3%) compared to HTT_sgRNA1 (Figure 1G). The ‘staggered cut’ signature was not observed.

Editing with HTT_sgRNA4 led to an interesting phenomenon, as, despite targeting the region 58 base pairs downstream of the CAG tract, it generated a significant amount of “clean” in-frame Del CAG (60%, Figure 1C). At the same time, it was the least efficient gRNA, as 13% of the reads contained full-length CAG tracts, and the most frequent variant was the unedited 41 CAG sequence (11.58%, Figure 1H). A relatively abundant fraction of edits comprised short deletions, e.g., a one CAG triplet deletion was the second most frequently detected variant (11.23%). The entire tract excision constituted 12% of all reads. Due to the nature of the edit, out-of-frame Del and complex Indels (18% and 4%, respectively) were more frequent than in the case of HTT_sgRNA2 but less frequent than in the case of HTT_sgRNA1 (Figure 1C). An interesting outcome that we observed was an excision of the entire tract, from ‘G’ of the first CAG triplet to the HTT_sgRNA4 cutting site, that appeared with a frequency of 3.38% (Figure 1H). The same editing outcome was observed in our previous work with the use of paired nickases and a combination of HTT_sgRNA1+4 sgRNAs [21]. Frameshift was considerably less frequent than in the case of HTT_sgRNA1 (26%) (Figure 1H).

The last analysed variant was a result of simultaneous editing with HTT_sgRNA1 and HTT_sgRNA4 (Figure 1I). As expected, the excision of the entire CAG tract was efficient (51%). The most frequent variant was a Del variant spanning the entire region between cleavage sites (27.18% of all reads). The frameshift frequency was 60% (Figure 1I).

We decided to use only HTT_sgRNA1 and HTT_sgRNA2 in the subsequent experiments, as they were more efficient than HTT_sgRNA4 in the induction of CAG tract shortening and led to more mechanistically interesting phenomena than the simple fragment excision observed in the case of HTT_sgRNA1 and HTT_sgRNA4 used simultaneously.

### Extended DNA end resection is engaged in the process of DSB repair within CAG repeats

The large deletions of CAG repeats we observed suggest the involvement of mechanisms dependent on the resection of DNA ends. To investigate the extent of these resections and the effect of DSB localization, we performed qPCR on genomic DNA isolated from cells edited by RNP (Cas9-HTT_gRNA1 or Cas9-HTT_gRNA2) and digested by a set of restriction enzymes (Supplementary Table S3) [26]. Sites that underwent resection were not digested by endonucleases due to their requirement of double-stranded DNA and consequently could be efficiently amplified in qPCR by primers flanking the restriction site. Therefore, we were able to assess the resection level by comparing the Cq values obtained in digested and nondigested fractions. Eight restriction sites were used: four located upstream (A, B, C, D) and four located downstream (E, F, G, H) of the CAG repeats in the *HTT* locus. The distances of these sites from the CAG tract were as follows: approximately 10 000 bp, 5 000 bp, 1000 bp and 200 bp. (Figure 2A). The relationship between the distance from Cas9-induced DSB and the resection level depending on the gRNA used was revealed. We observed that cleavage with HTT_gRNA1 led to efficient resection at sites proximal to the cutting site (D and E, approximately 35%) and less efficiently at sites 1000 bp distant from the DSB (C and F, >15%) (Figure 2A). No resection was detected in the distal regions (A, B, G, H). The maximum level of resection was observed 6 h post transfection in the proximal regions (D and E), and the edited site was completely repaired 48 h post transfection, as the level in each region was below the level of the nonedited control. In contrast, the maximum level of resection for HTT_gRNA2 was detected 24 h post transfection (approximately 35%) and in more distant regions compared to HTT_sgRNA1 (C and F) (Figure 2A). High resection levels were also observed in regions B and G (5000 bp from the CAG tract), but in the outermost regions (A and H), no resection was detected. After 48 h, the edited site was still being repaired, with the highest level detected in distal regions (B and G). The longer resection time observed for HTT_gRNA2 compared to HTT_gRNA1 may result from the possibility of its binding in several sites within the CAG sequence (max. 6) and generation of multiple DSBs or recutting of the edited template. The results from the experiment are consistent with the outcomes from capillary electrophoresis of PCR products generated on the DNA isolated from cells at several timepoints after transfection with RNP (Supplementary Figure S1). No shortenings of sequence were visible in the 4 h after the transfection with Cas9-HTT_gRNA1 as in this time resections were in progress. Interestingly, the shortening of the sequence 4 h after transfection with Cas9-HTT_gRNA2 was visible, although the resections were ongoing. This suggests the involvement of a nonresection-dependent DNA repair pathway in this case. To test whether such intense resection is characteristic of repetitive sequences, we performed a similar analysis for the low repetitive control region (only 6 CAG repeats), which was the second target *locus* (off-target) for HTT_gRNA1 (gRNA1_OT). Similarly, we chose four regions located approximately 1000 bp and 300 bp upstream and downstream of the edited site (Figure 2B). We demonstrated that there was no resection in this region, although it had been edited, which we had demonstrated using the T7E1 assay (Figure 2C). This may indicate that regions abundant in CAG repeats induce different types of DSB repair mechanisms compared to low-repetitive regions.

**Figure 2.**
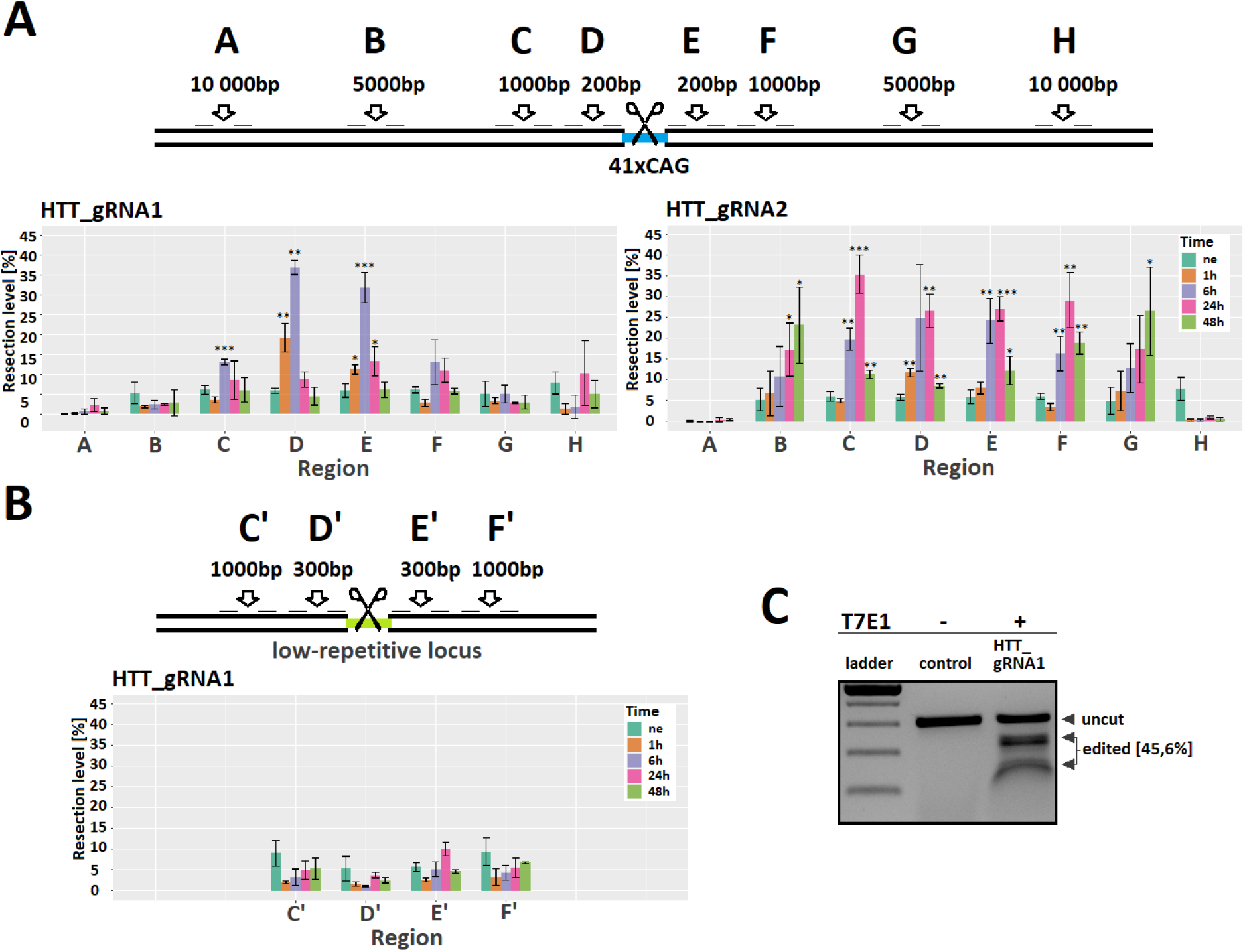
The level and range of DNA end resection after DSB induction. A) Restriction sites with their distances from the CAG repeats in the *HTT* locus. The extent and level of resections at given time points determined after induction of DSB using HTT_sgRNA1 and HTT_sgRNA2. B) Restriction sites with their distances from the second target of HTT_sgRNA1 (low-repetitive locus). No resection was detected in this region. C) The T7E1 assay confirmed the successful editing of the low repetitive locus after using HTT_gRNA1. The data shown represent the mean ± SD (n=3).

### MRE11 and FEN1 enrichment in regions flanking CAG repeats

MRE11 as the component of MRN complex, performs the initial processing of the DSB during DNA repair. Its 3’ to 5’ exo- and endonuclease activity are responsible for inducing resection at the cleaved site. It has been demonstrated in an *S. cerevisiae* model that the Mre11-Rad50 complex together with Sae2 is involved in DNA end processing at the TALEN-induced DSB site, leading to Rad52-mediated repeat contractions [7]. To identify whether MRE11 is involved in the initial resection of DNA ends leading to CAG repeat shortening, we assessed its enrichment in regions flanking the CAG repeats using ChIP−qPCR. We observed that cleavage with both guide RNAs led to a high enrichment of the MRE11 protein (Figure 3A); however, the pattern of enrichment varied depending on the location of DSBs. After cleavage with HTT_sgRNA1, MRE11 enrichment was very pronounced, especially at the right flank (15-, 17- and 8-fold increases at 12, 24 and 48 h, respectively). In contrast, after cleavage with HTT_sgRNA2, the enrichment at the right flank reached 7-fold only after 48 h, while amplification of the left flank resulted in high and consistent enrichment (15-, 12- and 8-fold increases at 12, 24 and 48 h, respectively). When homologous sequences exposed after resection anneal, overhangs called flap structures may form at the repair site. These flaps are the main substrate for FEN1 endonuclease. In addition, flaps containing trinucleotide repeats can fold into secondary structures that can be processed by FEN1 [34,35]. Therefore, we analysed FEN1 enrichment at regions flanking CAG repeats using ChIP−qPCR to identify whether FEN1 can be involved in CAG repeat shortening. Additionally we tested FEN1 enrichment in cells with different CAG repeat tract lengths (41 CAG and 83 CAG), as we supposed that the formation of structures removed by FEN1 should increase during DSB repair in longer repeat tracts. In cells with 41 CAG repeats, the FEN1 protein was slightly enriched in the right flanking region after inducing DSBs with both guide RNAs (Figure 3B). However, the distribution of enrichment varied over time depending on the location of the DNA break. For HTT_sgRNA1, we observed a 2-fold increase at 12 and 24 h. For HTT_sgRNA2, we observed more consistent enrichment, with an approximately 3-fold increase at 24 and 48 h and a 2-fold increase at 72 h. As expected, enrichment of the FEN1 protein after inducing DSBs within longer CAG repeat tracts was higher. Similar to the results for the 41 CAG repeat tract, for HTT_sgRNA1, we observed a 2-fold increase in the right flanking region at 12 and 24 h (Figure 3C). However, the enrichment values were also high for the left flanking region (4-fold increase at 12 and 24 h). After cleavage with HTT_sgRNA2, we observed high FEN1 enrichment in the left flanking region in cells with 83 CAG repeats (5-fold, 3-fold and 4-fold enrichment at 24, 48 and 72 h, respectively). Interestingly, the temporal distribution of FEN1 enrichment after DSB induction remained the same for cells with 41 CAG repeats and 83 CAG repeats (with enrichment after 12 and 24 h after cleavage with HTT_sgRNA1 and after 24, 48 and 72 h after cleavage with HTT_sgRNA2), implying that the kinetics of this process are independent of the CAG tract length.

**Figure 3.**
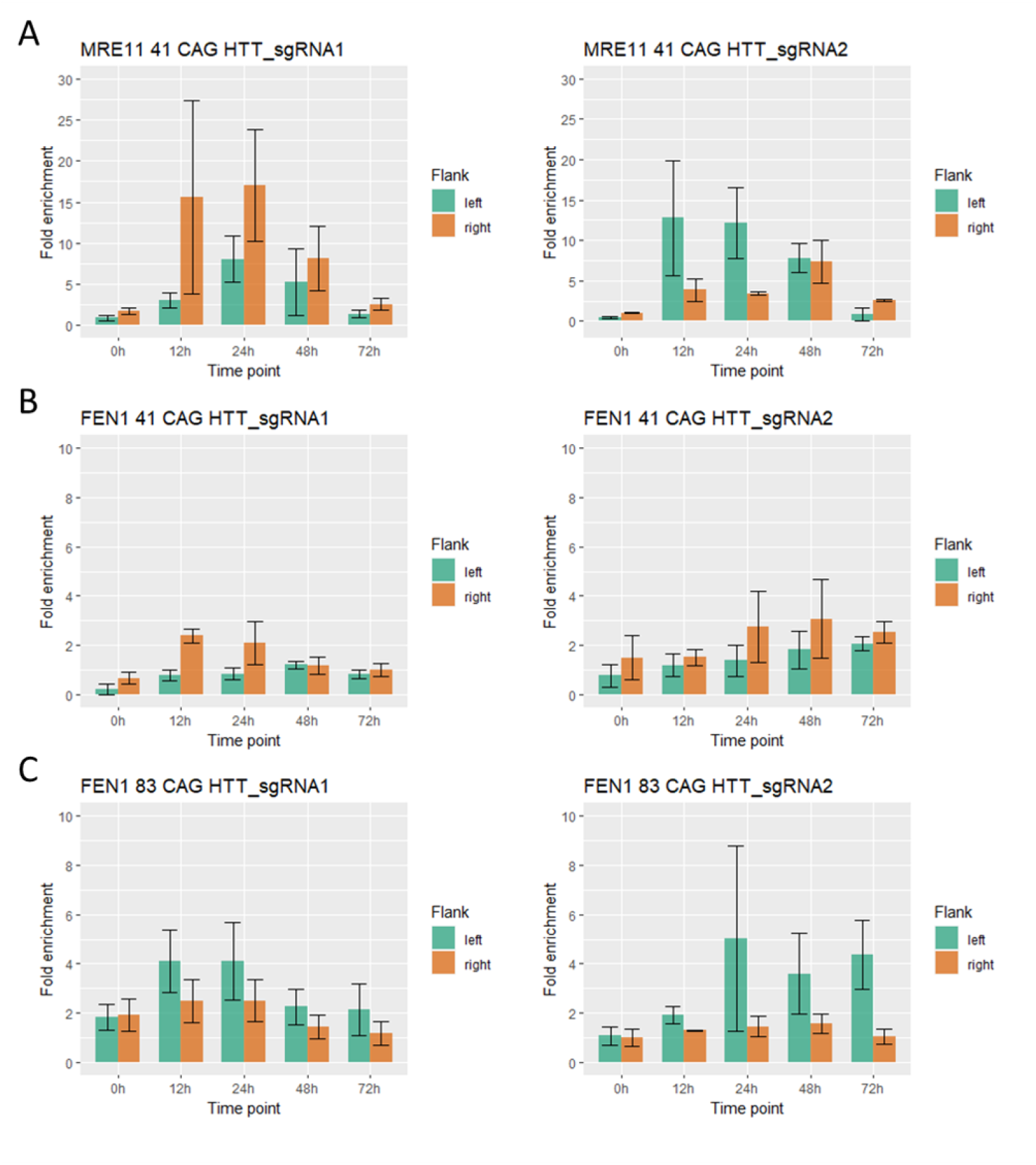
The pattern of repair protein enrichment depends on the location of DSBs within the *HTT* locus and differs between the right and left flanks of the CAG repeat region. A) The enrichment of MRE11 protein at given timepoints determined by ChIP-qPCR after induction of DSBs in the *HTT* locus with 41 CAG repeats using HTT_sgRNA1 and HTT_sgRNA2. B) The enrichment of FEN1 protein at given timepoints determined by ChIP-qPCR after induction of DSBs in the HTT locus with 41 CAG repeats or 83 CAG repeats (C) using HTT_sgRNA1 and HTT_sgRNA2. The data shown represent the mean ± SE (n=3). A 2-fold increase in value compared to the control region is considered significant. No significant differences were detected between protein enrichment on the right and left flanks and between time points.

### Searching for pathways involved in CAG repeat shortening by inhibiting key proteins involved in DSB repair

To gain deeper insight into the mechanisms behind CAG repeat shortening, we inhibited proteins representing different DNA repair pathways with chemical inhibitors or siRNAs (Figure 4A). We measured the ratio of signals from capillary electrophoresis peaks corresponding to unchanged vs. edited (shortened) products. After the induction of DSBs with HTT_sgRNA1 and HTT_sgRNA2, peaks corresponding to shortened PCR products constituted approximately 40% and 60% of all signals, respectively (Figure 4B). We used chemical inhibitors of 4 proteins (MRE11, CTIP, POLQ, KU70/80) and siRNAs to downregulate 5 genes (MRE11, POLQ, EXO1, RAD51, Artemis/DCLRE1C) (Supplementary Table S4). Then, we compared the ratio of shortened vs. unchanged products in treated and control samples (Figure 4C). Efficient inhibition of gene expression by siRNA was confirmed by Western blotting and RT−qPCR (Figure 4D). Inhibition and knockdown of the POLQ gene resulted in an ∼30% decrease in shortened products compared to controls after DSB induction with HTT_sgRNA1 (Figure 4E). Similarly, KU70/80 inhibition and EXO1 knockdown resulted in an 18% and 29% decrease in truncated products, respectively. MRE11, CTIP and RAD51 inhibition did not induce any changes in the fraction of shortened products compared to the control. Different proteins (except KU70/80) were determined to be significantly important in the CAG repeat contraction process after DSB induction with HTT_sgRNA2. Inhibition and gene knockdown of MRE11 resulted in 6% and 23%, respectively, decreases in shortened products compared to controls (Figure 4F). KU70/80, CTIP and RAD51 inhibition or gene knockdown resulted in 19%, 11% and 10%, respectively, decreases in shortened products compared to controls. Knockdown of Artemis/DCLRE1C had no effect on the proportion of the shortened products. These results indicate that DSB location may influence the selection of the DNA damage response and engage different DNA repair proteins.

**Figure 4.**
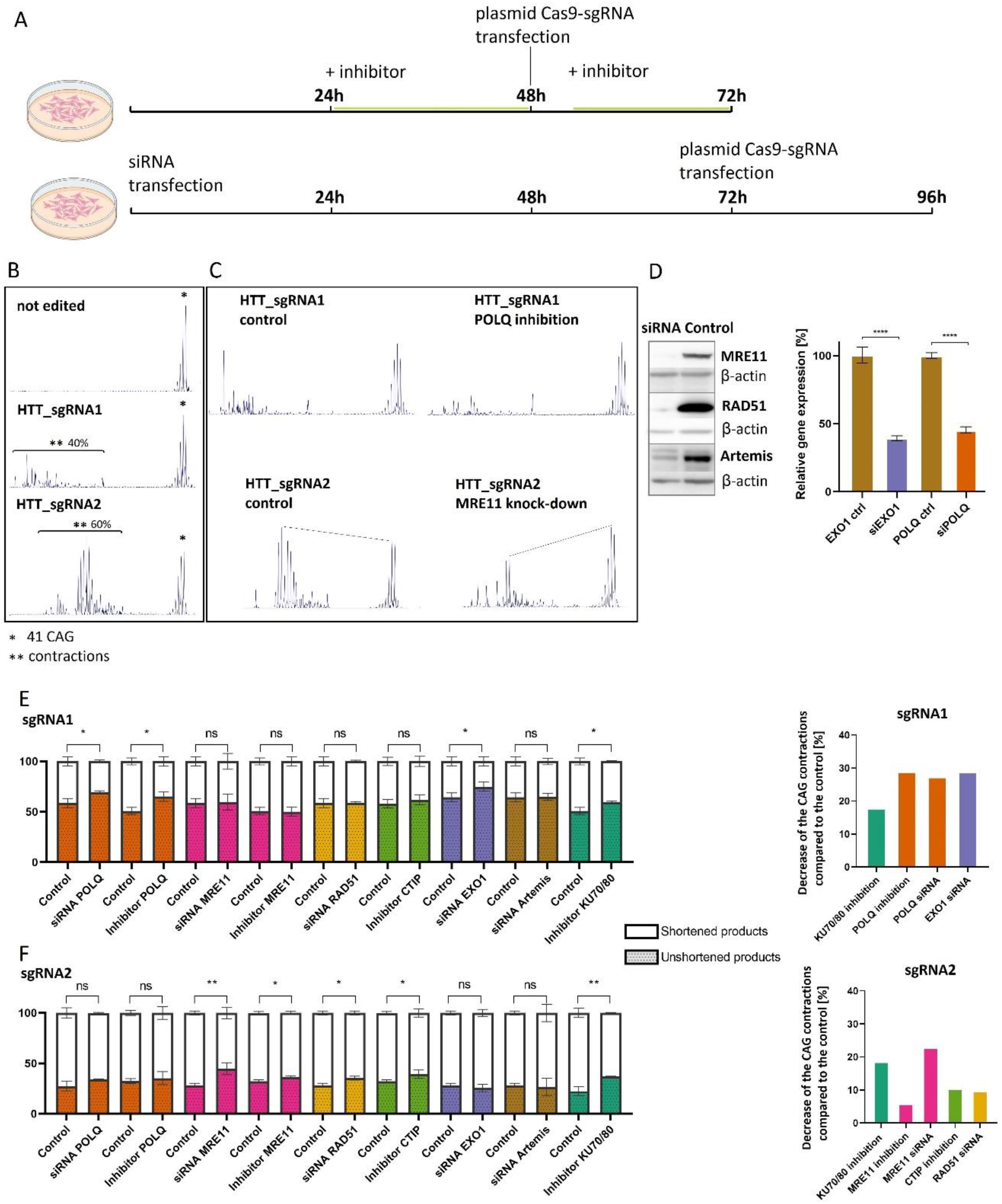
Effects of inhibiting selected DNA repair pathways with chemical inhibitors and siRNAs on CAG repeat shortening. A) Workflow of the experiment. B) Capillary electrophoresis of the PCR products after the *HTT* locus editing using HTT_sgRNA1 and HTT_sgRNA2. The percentage of CAG repeat contractions is shown. C) Comparison of peak heights corresponding to edited (contracted) and unedited products in the controls and treated samples. D) The silencing of gene expression after application of siRNA was confirmed by Western blotting or RT-qPCR. E-F) Percentage of shortened and 41 CAG sequences obtained under inhibition of individual proteins with chemical inhibitors or siRNA and DSB induction with HTT_sgRNA1 and HTT_sgRNA2, respectively. On the right, the percentage reduction in the number of shortened products due to silencing of individual DNA repair proteins. The data shown represent the mean ± SD (n=3).

### Unbiased detection of proteins interacting with the CAG tract during DNA break repair

To identify proteins involved in the repair of DNA spanning the CAG tract, we carried out engineered DNA-binding molecule-mediated chromatin immunoprecipitation (enChIP) followed by mass spectrometry (MS) (enChIP-MS) [27]. We transduced HEK293T-41CAG cells with a lentiviral vector coexpressing 3xFLAG-tagged Cas9 devoid of nuclease activity (dCas9-FLAG) and sgRNA targeting the flanking region upstream of the CAG tract in the *HTT* gene. After confirming the constitutive expression of dCas9-FLAG and efficiency of the pull-down (Supplementary Figure S2), DSBs were introduced using HTT_sgRNA1- or HTT_sgRNA2- expressing plasmids, and then complexes containing FLAG-dCas9 were immunoprecipitated and subjected to mass spectrometry for protein identification. The unedited cells were used as a control. Five replicates of every experimental variant were used.

A total of 402 and 404 proteins were identified as being differentially abundant (q-value < 0.05, |fold change| ≥ 1.5) in comparison to the noncleaved control for HTT_sgRNA1 or HTT_sgRNA2, respectively (Supplementary Figures S3 and S4). A STRING [36] network revealed that the identified proteins clustered into two relatively tight groups (translation and mRNA processing) and a looser cluster of proteins involved in chromatin organization and DNA repair. The levels of a total of 278 proteins were increased and the levels of 124 proteins were decreased in the case of HTT_sgRNA1. The ratio was 275 to 129 for HTT_sgRNA2. Among the proteins that increased in level, the category of translation and DNA repair was enriched, whereas the category of chromatin organization was enriched among the proteins that decreased in level. The proteins related to mRNA processing were present in both the increased and decreased level groups.

The proteins that were enriched in the category of DNA repair were further analysed (22 genes for HTT_sgRNA1 and 24 genes for HTT_sgRNA2, Figure 5A-B). Figure 5C depicts the STRING network of identified DNA repair-related proteins further divided into categories of DNA repair pathways. We detected proteins involved in the regulation and execution of DNA repair related to DSB repair as well as to SSB repair. We detected the presence of the KU complex: XRCC5 (fold change of 9.8 and 12.9 for HTT_sgRNA1 and HTT_sgRNA2, respectively) and XRCC6 (fold change of 5.6 and 3.9 for HTT_sgRNA1 and HTT_sgRNA2, respectively), as well as PRKDC protein (fold change of 2), which are known to be involved in NHEJ repair. Additionally, the PARP1 protein, which is considered a key player in the initiation of DNA repair, was detected (2.9-fold and 3.6-fold increase in the case of HTT_sgRNA1 and HTT_sgRNA2, respectively). We observed enrichment of the components of the MCM2-7 complex (MCM3, MCM4, MCM6, MCM7) involved in BIR and HDR. Ubiquitins related to the NER and ICLR pathways were also enriched (UBB, UBA52, UBC, RPS27A: fold changes of the entire group: 6.3 and 4.2 for HTT_sgRNA1 and HTT_sgRNA2, respectively). We also identified both components of the histone chaperone FACT (SUPT16H, SSRP1, fold changes ∼8), central component of the condensin complex (SMC2, fold changes of 5.5 and 10.5, respectively) and two cohesins (SMC1A, SMC3, fold changes of 6.7 and 16.4 for HTT_sgRNA1 and HTT_sgRNA2, respectively). Interestingly, we also detected that the nuclease FEN1 was involved in the DSB processing, but only in the case of HTT_sgRNA2 (2.6-fold increase). The HTT_sgRNA2 variant was also enriched with the SFPQ protein (1.5-fold increase), which may be involved in HDR. The set of proteins significantly downregulated in the DNA repair group comprised BLM, HMGA1, HMGA2, HMGB1, HMGB2 and HIST1H4A. Outside of the DNA repair category, we found other proteins of potential interest exhibiting level changes, such as increased levels of exportins (CSE1L, XPO1), components of the nuclear pore (NUP205, NUP93) or decreased levels of DNA topoisomerase (TOP1). Supplementary Table S5 contains the numbers of peptides detected by MS for proteins enriched in the gene ontology category of DNA repair.

**Figure 5.**
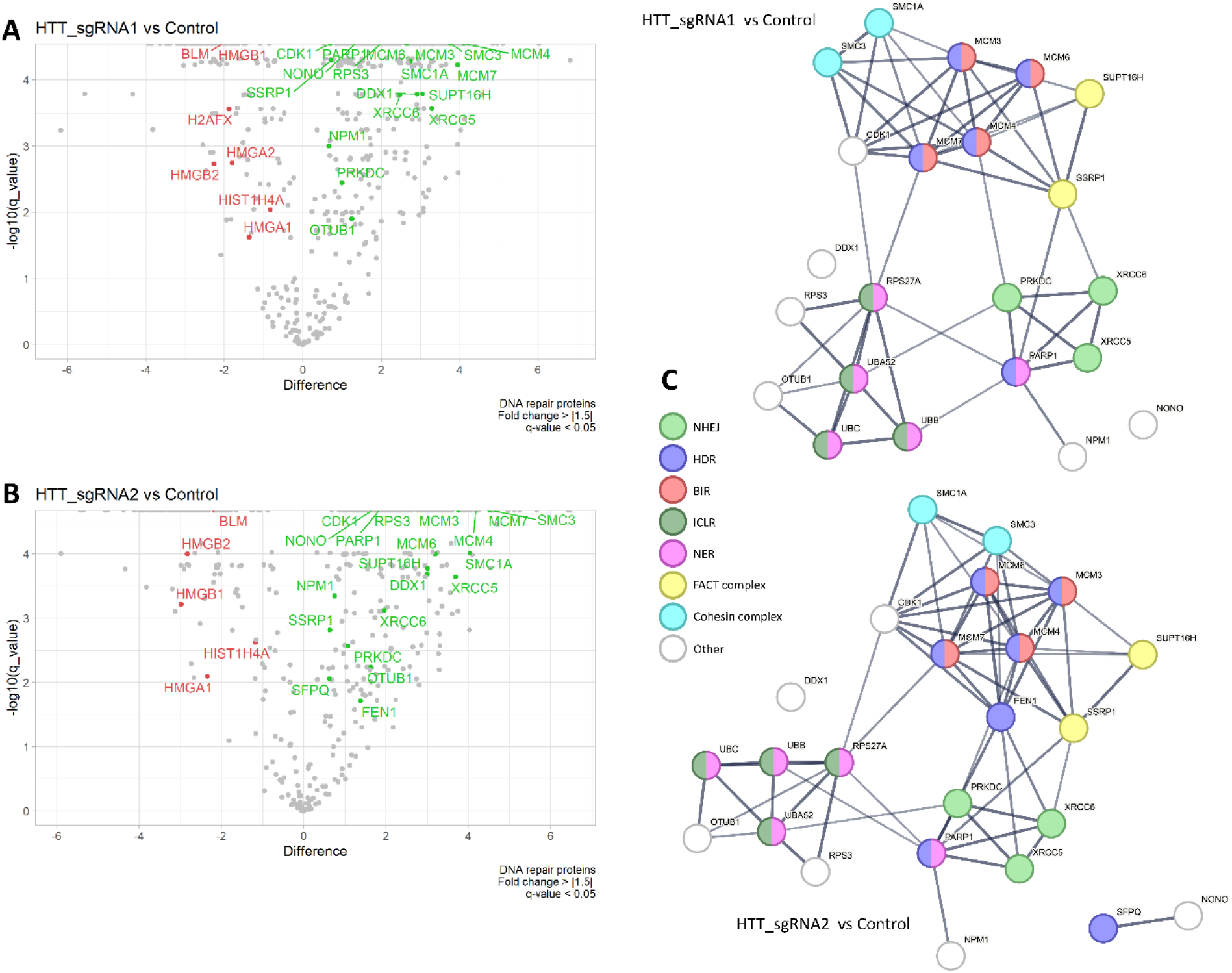
Unbiased analysis of proteins involved in DNA break repair within the CAG repeat region using enChIP- MS. A) Volcano plot presenting the distribution of the proteins identified in enChIP-MS analysis after the introduction of DSBs by using HTT_sgRNA1. The “Difference” parameter is the difference between the mean of log10 (protein spectrum intensity) of the experimental groups. The proteins enriched in the gene ontology category of DNA repair and characterized by |fold change| >= 1.5 and q-value < 0.05 are marked in red (decreased levels) or green (increased levels). B) Volcano plot presenting data obtained in enChIP-MS analysis after introduction of DSB by using HTT_sgRNA2. C) STRING networks of the proteins that were enriched in gene ontology category of DNA repair divided into subcategories. The networks were generated for data obtained after the introduction of DSBs by using HTT_sgRNA1 (upper network) and HTT_sgRNA2 (lower network).

The enChIP analysis showed enrichment of a significant number of helicases involved in the process of DNA:RNA hybrid unwinding (e.g., DDX18, DDX1, DHX9, DDX5 and DDX21). DNA:RNA hybrids, particularly R-loops, occur naturally as a result of transcription, DNA replication and DNA repair [37]. To verify whether DNA:RNA hybrids are created within the *HTT* locus during DNA repair, we performed DRIP-qPCR analysis after inducing DSBs with HTT_sgRNA1 and HTT_sgRNA2 (Supplementary Figure S5A). The results show that DNA:RNA hybrids were 2 times more abundant at the flanking regions of the CAG repeat tract after cleavage with HTT_sgRNA2 in relation to the negative control locus (ZNF554). After inducing DSBs with HTT_sgRNA1, no significant enrichment of DNA:RNA hybrids was found in the *HTT* locus. However, it should be noted that in comparison to previously published DNA:RNA hybrid-rich loci for HEK293T cells (LOC440704 and ING3, [38]), the enrichment in the *HTT* locus is noticeably lower. This may be due to the limitations caused by the efficiency of cell editing or from the reduced stability of the hybrids in the studied locus. To further verify the presence of DNA:RNA hybrids during DSB repair within the *HTT* locus after inducing breaks with HTT_gRNA2, we performed immunofluorescence analysis using anti-DNA:RNA hybrid and anti-γH2AX antibodies (Supplementary Figure S5B). The colocalization of the DNA:RNA hybrids and γH2AX histone was not significant. However, signals from the DNA:RNA hybrids in relation to γH2AX were higher in HEK293T-41CAG cells than in control cells after inducing DSBs with HTT_gRNA2 (Supplementary Figure S5C).

## DISCUSSION

Previous studies that were conducted mainly on yeast models and *in vitro* systems have identified potential proteins and pathways associated with the shortening of CAG/CTG tracts. However, it is important to note that the studies differed in their focus and methodologies, resulting in fragmented and limited knowledge regarding the underlying mechanism. Specifically, it has remained unclear whether these processes are conserved in human cells. Nevertheless, the picture emerging from the studies seemed to primarily involve the single-strand annealing (SSA) pathway.

In this study, we demonstrate the phenomenon of tract shortening in an endogenous model of the human *HTT* gene, revealing that the process is highly complex and dependent on the site of the DNA double-strand break (DSB). The analysis of the sequences of the final products of repair suggests that different mechanisms may act simultaneously on the substrate to produce various products. As expected, repair is initiated by the recruitment of NHEJ factors, but then resection of DNA ends directs the process towards pathways involving the use of microhomology.

First, our investigation revealed an extensive DNA end resection process, particularly in cases of DSBs occurring within the repetitive tract, reaching up to 5,000 base pairs. To date, the resection process has only been studied in yeast. Mosbach et al. observed resections reaching ∼1900 bp upstream and ∼2900 bp downstream from the CTG repeats [7]. In contrast, we did not observe asymmetry of the resection after HTT_gRNA1 or HTT_gRNA2 DSB induction. Importantly, our results suggest that MRE11 plays a role in initiating the resection process, as evidenced by the involvement of this protein regardless of the gRNA used.

In light of our experimental findings, we propose a model explaining the mechanism underlying CAG/CTG tract shortening following Cas9-mediated DSB. First, upon cleavage by Cas9-gRNA1, a staggered cut is generated, accompanied by the presence of NHEJ factors at the cleavage site. Intriguingly, we observed an A insertion at the site of cleavage in accordance with the -4 rule [31,39,40], which supports NHEJ involvement in the process of DSB repair (Figure 6A). Products containing insA mutations typically exhibit small deletions in the CAG repeats, which can arise from DNA slippage during replication or repair [3]. Following cleavage, Cas9 remains tightly bound with both DNA ends and releases 3’-non-target single-stranded CTG strand first [41]. Cas9 can be removed by the FACT complex [42], which we also identified in enCHIP experiments. Unrepaired DNA ends, not resolved through classical NHEJ, can enter an alternative repair pathway involving MRE11-dependent short-range resections. The single-stranded CTG strand can adopt a flap or hairpin structure [3], which can be processed by endo- or exonucleases such as MRE11 and FEN1 [43].

**Figure 6.**
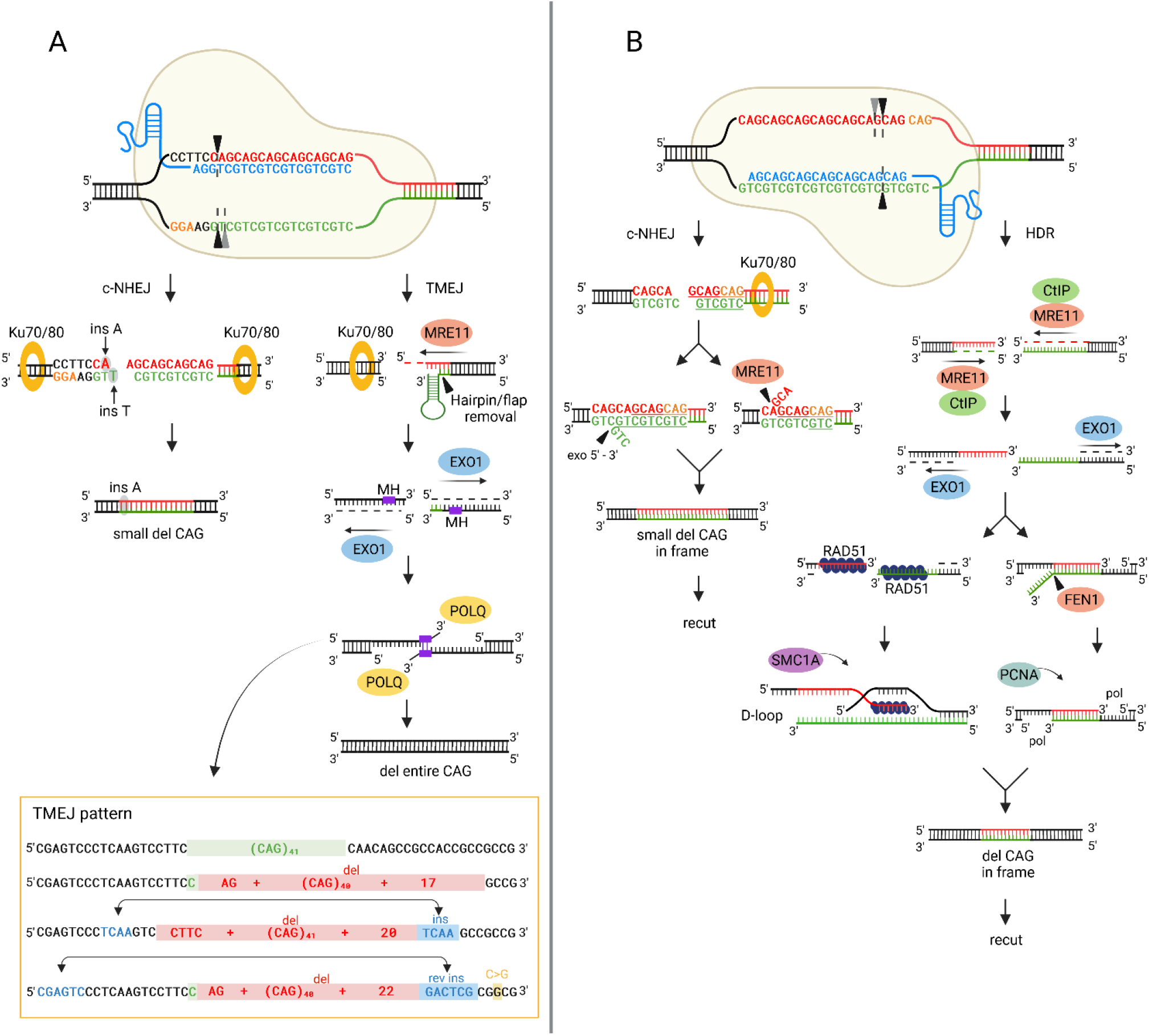
Scheme showing the complex repair process of double-strand breaks within the CAG trinucleotide repeat tract. A) After inducing DSBs with CRISPR‒Cas9 at the beginning of the CAG repeat tract (HTT_sgRNA1), repair is possible by either canonical NHEJ (c-NHEJ) or theta-mediated end joining (TMEJ). Staggered cut processing through c-NHEJ leads to repair products with small CAG repeat deletions and an insertion of a single adenine 4 nucleotides from the cleavage site. TMEJ leads to a complete deletion of the CAG repeat tract. Resections performed by MRE11 and EXO1 expose microhomologous regions (MH), which are utilized by POLQ to tether two single-stranded DNA ends, followed by gap filling. Sequence examples with patterns typical for TMEJ from NGS results are highlighted in the box. B) After inducing multiple DSBs within the CAG repeats (HTT_sgRNA2), two repair pathways are activated: c-NHEJ and HDR. Rapid repair of staggered cuts by c-NHEJ results in small in-frame CAG repeat deletions. The target site for CRISPR‒Cas9-HTT_sgRNA2 was reconstructed, and recutting was possible. HDR requires extensive resections performed by MRE11-CTIP and EXO1. The resulting long single-stranded DNA fragments with homologous sequences anneal. This process may be dependent on the RAD51 protein and D-loop formation, and additionally facilitated by cohesin SMC1A. Alternatively, after strand assembly, the resulting flap structures can be removed by FEN1 in a process independent of RAD51, and the resulting gaps are filled in by a polymerase in cooperation with the PCNA protein. HDR also causes CAG repeat deletion in-frame, allowing for recut.

This observation aligns with previous findings from the Guy-Franck Richard group in studies conducted on yeast, that demonstrated involvement of the Mre11-Rad50-Sae2 complex on the side of the break containing the repetitive tract after asymmetrical cleavage [7,8]. Then, MRE11 recruits the EXO1 protein, thereby facilitating long-range 5’-3’ resection. Aside from the observed resection events, the involvement of EXO1 is supported by the findings of siRNA experiments. Finally, our data suggest the potential involvement of polymerase theta-mediated end joining, resulting in deletion of the entire CAG repeat tract. Short microhomologous sequences located within flanks are used for anchoring DNA strands, and POLQ fills the gaps. DNA end resection, coupled with the lack of homologous recombination substrates (both alleles are cleaved) and the presence of microhomologies, implies that TMEJ may be the underlying mechanism. This notion is supported by the results of experiments involving POLQ chemical inhibitors and siRNA. Notably, even a 50% reduction in POLQ expression resulted in a significant reduction in products with delCAG. Moreover, we observed characteristic sequence mutational signatures of TMEJ, including templated insertions (Figure 6A, Supplementary File 1) [44]. This finding is particularly intriguing considering that POLQ is not expressed in *Saccharomyces cerevisiae* [45], highlighting the importance of investigating these mechanisms in human cells.

In the case of gRNA2-guided cleavage, the mechanism underlying tract shortening differs considerably, which is particularly evident in the sequence of edited products. Remarkably, the majority of these products (∼90%) exhibit in-frame deletions of CAG repeats. Furthermore, products with 4-10 repeats are predominant, which aligns with the findings reported in a previous study in a yeast model [7]. If, as with the use of gRNA1, Cas9-gRNA2 generated staggered cuts, the simplest way to repair the DSB would be to align the strands using short regions of microhomology with subsequent removal of small flaps (Figure 6B). This repair, probably c-NHEJ, results in small deletions of CAG repeats. The fact that gRNA2 binds entirely inside the repetitive tract makes subsequent recutting events possible, raising the possibility of an iterative nature of the repair process, a phenomenon suggested previously by Mosbach et al. [7]. A prolonged repair process may induce DNA resection, resulting in the release of DNA strands composed of CAG and CTG repeats of varying lengths. Strand realignment and hairpin/flap removal result in tract shortening. Our data indicate the contribution of FEN1, a structure-specific endonuclease, which may participate in the repair process by removing long flaps or hairpin structures of CTG and CAG repeats. According to previous reports, the activity of the FEN1 protein is important for the instability of the CAG repeat tract in the *HTT* gene during Long-Patch-BER [46–49] and MMEJ [50]. However, until now, the importance of FEN1 has been mainly analysed in the context of CAG repeat expansion. In the present study, we show that FEN1 also functions during the repair of DSBs within the *HTT* locus, leading to shortening of the CAG repeat tract. Long DNA resections detected after DSB induction with Cas9-gRNA2 are characteristic for HDR and SSA. In our experiments where RAD51 (key HDR protein) was effectively knocked down, the level of CAG deletions decreased, suggesting the involvement of this protein in CAG contractions. RAD51 in cooperation with other factors forms filaments on the 3’ ssDNA tail and invades donor dsDNA [51–53], creating a D-loop, followed by DNA synthesis performed by replicative DNA polymerases [51]. enChIP experiments revealed the enrichment of proliferating cell nuclear antigen (PCNA), the enhancer of the processivity of replicative DNA polymerases [52]. The enrichment of PCNA in gRNA2 samples was 60% higher than that in gRNA1 samples. Similarly, SMC1A (part of the recombination protein complex (RC-1) involved in DNA repair via recombination) [53] enrichment was almost 2.5-fold higher in gRNA2 samples than in gRNA1 samples. These evidence supports the involvement of the recombination-dependent pathway in creating in-frame CAG deletions without insertions after DSB induction with gRNA2. The same product may also be formed by the SSA pathway, as suggested in many previous studies.

The outcomes from unbiased proteomics conducted using enChIP support some previous findings. For instance, the association of the elements of the nuclear pore complex with the repetitive tract aligns with the model proposing the relocalization of CAG trinucleotide repeats to the nuclear pore complex taking place during DNA repair [54]. Furthermore, proteomics analysis has yielded novel insights revealing the participation of helicases in the repair machinery at the site of CAG repeats. The results confirm the presence of R-loops within the CAG repetitive tract, suggesting that helicases may play a role in resolving these R-loops formed by the interaction between single-stranded DNA and complementary RNA. R-loops have been implicated in diverse biological processes, such as DNA repair and transcriptional regulation [37]. The presence of R-loops at the CAG repetitive tract sheds light on their potential involvement in the repair mechanisms specific to repetitive sequences.

Admittedly, our study faced certain limitations in terms of identifying proteins that could support the proposed hypothesis. One of the limitations is the lack of cross-linking during the enChIP assay, which may result in the loss of certain proteins during the experimental process. Additionally, proteins such as POLQ are known to have low expression levels in cells, further complicating their detection. Moreover, the repair process itself is dynamic, and the timing of the experimental assays is crucial to capture specific events accurately. Technical constraints also required us to use plasmids alongside RNP in the ChIP experiments, which can influence the repair kinetics. Considering these limitations, further investigations with improved techniques and methodologies are necessary to gain a more comprehensive understanding of the involved proteins and the precise dynamics of the repair process. Importantly, our results demonstrate that the mechanism of Cas9 action itself seems to influence the repair process. We observed the insA event, in accordance with the -4 rule, only in the case of gRNA1.

In perspective, there are several promising directions for further exploration. In the next steps, investigating the effects of heterozygosity could provide valuable insights into the possible role of HDR. Currently, our study design does not allow the sister chromatids to serve as a DNA repair template. By studying heterozygous cells, we can examine how the presence of two different alleles affects the HDR process and, further, its possible role in repetitive tract shortening. Additionally, utilizing nondividing neuronal cells can offer a relevant cellular context for studying CAG tract shortening, as neurodegenerative disorders such as HD primarily affect neurons. This approach would provide valuable information on the repair mechanisms and dynamics specifically in postmitotic cells. Furthermore, efforts should be made to gain better control over the tract shortening process. Exploring strategies to modulate and regulate repair outcomes, such as controlling the cell cycle, or using inhibitors of selected DNA repair pathways, could enhance our ability to precisely control and manipulate CAG tract shortening.

## AUTHOR CONTRIBUTIONS

Pawel Sledzinski: Conceptualization, Methodology, Validation, Visualization, Writing—original draft. Mateusz Nowaczyk: Conceptualization, Methodology, Validation, Visualization. Marianna Karwacka: Conceptualization, Methodology, Validation, Visualization. Marta Olejniczak: Conceptualization, Writing—review & editing.

## ACKNOWLEDGEMENTS

We would like to thank Magdalena Dabrowska for the cell line generation, Damian Brauze for support during the ChIP experiments Agnieszka Fedoruk-Wyszomirska for support during the microscopic analyses and Magdalena Trybus from the Laboratory of Single Cell Analyses IBCH PAS for performing flow cytometry experiments. Bioinformatic analysis was performed by ideas4biology Ltd. Proteomic mass spectrometry analyses were performed in the Mass Spectrometry Laboratory, IBB PAS, Warsaw. The equipment used was sponsored in part by the Centre for Preclinical Research and Technology (CePT), a project cosponsored by European Regional Development Fund and Innovative Economy, The National Cohesion Strategy of Poland. The flow cytometry data collected in this study were acquired using the infrastructure developed under the project NEBI - National Imaging Centre for Biological and Biomedical Sciences, POIR.04.02.00-00-C004/19, co-financed through the European Regional Development Fund (ERDF) in the frame of Smart Growth Operational Programme 2014-2020 (Measure 4.2 Development of modern research infrastructure of the science sector).

## TABLE AND FIGURE LEGENDS

**Supplementary Figure S1.**

PCR products of the CAG locus in the HTT gene resolved by capillary electrophoresis. A) Cells were collected at given timepoints after electroporation with RNP containing gRNA1 or B) gRNA2. C) Insertion of A detected 24 h posttransfection in cells edited with gRNA1.

**Supplementary Figure S2.**

A) Sequence of the region containing the CAG repetitive tract in the HTT gene. The site complementary to the gRNA guiding dCas9-FLAG is underlined. The PAM sequence is highlighted in green. The sequence amplified during qPCR is highlighted in blue. B) Expression of dCas9-FLAG after lentiviral transduction. C7 – HEK293T-41CAG clone expressing dCas9-FLAG. Cell lysates were subjected to Western blot analysis with an anti-Cas9 antibody. β-Actin was used as a loading control. WT – HEK293T-41CAG cells not subjected to lentiviral transduction. C) Isolation of the HTT CAG repetitive locus by enChIP with dCas9-FLAG. qPCR analysis of enrichment was performed. The locus encoding the β-actin gene was used as a background control.

**Supplementary Figure S3.**

A) Volcano plot presenting the distribution of all the proteins identified in enChIP-MS analysis after introduction of DSBs by using HTT_sgRNA1. The “Difference” parameter is the difference between the mean of log10(protein spectrum intensity) of the experimental groups. The proteins characterized by |fold change| >= 1.5 and q-value < 0.05 were considered significant and are marked in red (downregulated) or green (upregulated). B) A STRING network of proteins identified in enChIP-MS analysis after introduction of DSB by using HTT_sgRNA1. Proteins exhibiting low fold change (|fold change| < 1.5) or high q-value (q-value > 0.05) were filtered out. The STRING confidence level was set to the highest (0.900). The proteins that were enriched in the gene ontology category of DNA repair were analysed further (see main text).

**Supplementary Figure S4.**

A) Volcano plot presenting the distribution of all the proteins identified in enChIP-MS analysis after introduction of DSBs by using HTT_sgRNA2. The “Difference” parameter is the difference between the mean of log10 (protein spectrum intensity) of the experimental groups. The proteins characterized by |fold change| >= 1.5 and q-value < 0.05 were considered significant and are marked in red (decreased levels) or green (increased levels). B) A STRING network of proteins identified in enChIP-MS analysis after introduction of DSB by using HTT_sgRNA2. Proteins exhibiting low fold change (|fold change| < 1.5) or high q-value (q-value > 0.05) were filtered out. The STRING confidence level was set to the highest (0.900). The proteins that were enriched in the gene ontology category of DNA repair were analysed further (see main text).

**Supplementary Figure S5.**

Presence of DNA:RNA hybrids in the process of DSB repair at the *HTT* locus. A) The enrichment of DNA:RNA hybrids determined by DRIP-qPCR in the positive control locus (LOC440704 and ING3) and in the regions flanking the CAG repeat tract in the *HTT* locus after induction of DSBs using HTT_sgRNA1 and HTT_sgRNA2, respectively. DNA:RNA hybrids are 2-fold enriched in the *HTT* locus after inducing DSBs with HTT_sgRNA2. No significant enrichment of DNA:RNA hybrids was observed after inducing DSBs with HTT_sgRNA1. The data shown represent the mean ± SD (n=2). B) Representative images of nuclei of HEK293T, HEK293T-41CAG and HEK293T-83CAG cells after inducing DSBs with HTT_gRNA2 with foci of γH2AX (red) and RNA:DNA hybrids (green). Scale bar = 6 μm. C) Graphs representing Pearson’s coefficient for colocalization of γH2AX and RNA:DNA hybrid foci (n=79, 94, and 105, respectively) and the ratio of the fluorescence intensity from DNA:RNA hybrids to the fluorescence from γH2AX (n=18, 16, and 19, respectively) within nuclei of HEK293T, HEK293T-41CAG and HEK293T- 83CAG cells.

