## Supplemental tables and figures for "CRISPR/Cas9-induced double-strand breaks in huntingtin locus lead to CAG repeat contraction through the extensive DNA end resection and homology-mediated repair"

### Supplementary Table S1. Oligonucleotides used in the study

| Oligonucleotide ID | Sequence (5'-3') | Description |
|---|---|---|
| LHD_F | CACACACAGCTTCGCCTCAC | RT-qPCR primer, left flank of exon 1 of *HTT* gene |
| LHD_R | GTTCTGCCTCACACAGCAAGG | RT-qPCR primer, left flank of exon 1 of *HTT* gene |
| RHD_F | CTGCACCGACCGTGAGTTTGGG | RT-qPCR primer, right flank of exon 1 of *HTT* gene |
| RHD_R | TGGGTCACTCTGTCTCTGCGGG | qRT-PCR primer, right flank of exon 1 of *HTT* gene |
| BACT_F | TGAGAGGGAAATCGTGCGTG | RT-qPCR primer for β-actin gene |
| BACT_R | TGCTTGCTGATCCACATCTGC | RT-qPCR primer for β-actin gene |
| EXO1_F | ACTAAGCTACGCTGGGCAATATG | RT-qPCR primer for *EXO1* gene |
| EXO1_R | ATGACTTCTTGAATGGGCAGG | RT-qPCR primer for *EXO1* gene |
| POLQ_F | GCCTTGGTCGCTGCCTGAA | RT-qPCR primer for *POLQ* gene |
| POLQ_R | AGGAAGTCCCCAGTTTGCCA | RT-qPCR primer for *POLQ* gene |
| GAPDH_F | GAAGGTGAAGGTCGGAGTC | RT-qPCR primer for *GAPDH* gene |
| GAPDH_R | GAAGATGGTGATGGGATTTC | RT-qPCR primer for *GAPDH* gene |
| sgRNA1_S | CACCGCTGCTGCTGCTGCTGCTGGA | oligo for HTT_sgRNA1 plasmid construction |
| sgRNA1_A | AAACTCCAGCAGCAGCAGCAGCAGC | oligo for HTT_sgRNA1 plasmid construction |
| sgRNA2_S | CACCGAGCAGCAGCAGCAGCAGCAG | oligo for HTT_sgRNA2 plasmid construction |
| sgRNA2_A | AAACCTGCTGCTGCTGCTGCTGCTC | oligo for HTT_sgRNA2 plasmid construction |
| IG36 | CCGCTCAGGTTCTGCTTTTA | PCR primer used in amplification of the CAG tract in *HTT* gene |
| IG37 | GGCTGAGGCAGCAGCGGCTG | PCR primer used in amplification of the CAG tract in *HTT* gene |
| IG30 | GCTGATGAAGGCCTTCGAGT | PCR primer used in amplification of the CAG tract in *HTT* gene |
| IG31 | GCTGAGGCAGCAGCGGCT | PCR primer used in amplification of the CAG tract in *HTT* gene |
| ZNF554_F | CGGGGAAAAGCCCTATAAAT | RT-qPCR primer for negative DNA:RNA hybrid locus |
| ZNF554_R | TCCACATTCACTGCATTCGT | RT-qPCR primer for negative DNA:RNA hybrid locus |
| LOC440704_F | TAACACACAGCCATTGGGAAA | RT-qPCR primer for positive DNA:RNA hybrid locus |
| LOC440704_R | TTTTGGTAAGAGGGTTAAGCAGTA | RT-qPCR primer for positive DNA:RNA hybrid locus |
| ING3_F | TTTTTCTTCTCTAACTACCCTCCCC | RT-qPCR primer for positive DNA:RNA hybrid locus |
| ING3_R | GTGCCCTAATCTGAATGACTACA | RT-qPCR primer for positive DNA:RNA hybrid locus |
| IG126 | AGTTGTCAAAGTGCTACCCTTC | PCR primer used in amplification of the region A (resection analysis) |
| IG127 | GGTGTGCTAACATTCCCATGTC | PCR primer used in amplification of the region A (resection analysis) |
| IG124 | AAGCTCCTTACCAGCGGTGC | PCR primer used in amplification of the region B (resection analysis) |
| IG125 | CCCGTCTCATAGGATGGCTG | PCR primer used in amplification of the region B (resection analysis) |
| IG50 | GGGGTCACACTTGGGGTCCTCA | PCR primer used in amplification of the region C (resection analysis) |
| IG51 | GCCAGAGCCATACTCACCCGGA | PCR primer used in amplification of the region C (resection analysis) |
| IG68 | CACACACAGCTTCGCCTCAC | PCR primer used in amplification of the region D (resection analysis) |
| IG69 | GTTCTGCCTCACACAGCAAGG | PCR primer used in amplification of the region D (resection analysis) |

| | | |
|---|---|---|
| IG46 | CTGCACCGACCGTGAGTTTGGG | PCR primer used in amplification of the region E (resection analysis) |
| IG47 | TGGGTCACTCTGTCTCTGCGGG | PCR primer used in amplification of the region E (resection analysis) |
| IG48 | CCAACACGTTGCTGATGGGGAGG | PCR primer used in amplification of the region F (resection analysis) |
| IG49 | GGCACATCTGAGATGCCCACGG | PCR primer used in amplification of the region F (resection analysis) |
| IG128 | CAGTAGCTTGCGTTATCAGGTT | PCR primer used in amplification of the region G (resection analysis) |
| IG129 | TGTCTTCTGATAAGCTCTTGCTTG | PCR primer used in amplification of the region G (resection analysis) |
| IG130 | CAGCCATTGGTGAACTCGTGC | PCR primer used in amplification of the region H (resection analysis) |
| IG131 | GTTATACTCCATGTTGCGGGC | PCR primer used in amplification of the region H (resection analysis) |
| IG94 | GTTCTCCGGATGAGTCCTGC | PCR primer used in amplification of the region C' (resection analysis) |
| IG95 | ACCCTGGTCCTCGTTGTTTC | PCR primer used in amplification of the region C' (resection analysis) |
| IG104 | GGGTTCTGGAGGAAAGCTCT | PCR primer used in amplification of the region D' (resection analysis) |
| IG105 | TCCCTCTCCCCACCTCTGA | PCR primer used in amplification of the region D' (resection analysis) |
| IG110 | GTTGCCACTGGACAGTTGAT | PCR primer used in amplification of the region E' (resection analysis) |
| IG111 | GCCTCCAAGTCCACTTGCC | PCR primer used in amplification of the region E' (resection analysis) |
| IG100 | CCCTGGTCTGCAGAAACTGA | PCR primer used in amplification of the region F' (resection analysis) |
| IG101 | GACAGAAGAAGCTTCAAGCCC | PCR primer used in amplification of the region F' (resection analysis) |

### Supplementary Table S2. Antibodies used in the study

| **Antibody name** | **Supplier** |
|---|---|
| Mouse monoclonal antibody anti-FEN1 (GTX70185) | GeneTex |
| Mouse monoclonal antibody anti-Cas9 (7A9) | Sigma Aldrich |
| Rabbit polyclonal anti-B-actin (#4967) | Cell Signaling Technology |

| | |
|---|---|
| Mouse monoclonal antibody anti-DNA:RNA hybrid, clone S9.6 (MABE1095) | Sigma Aldrich |
| Rabbit monoclonal antibody anti-phospho-histone H2AX-S139 (AP0687) | ABclonal |
| Rabbit polyclonal antibidy anti-MRE11 (NB100-142) | Novus Biologicals |
| Rabbit monoclonal antibody anti-Artemis (D708V) | Cell Signaling Technology |
| Rabbit monoclonal antibody anti-RAD51 (D4B10) | Cell Signaling Technology |
| Donkey Anti-Mouse IgG (715-035-150) | Jackson ImmunoResearch |
| Donkey Anti-Rabbit IgG (711-035-152) | Jackson ImmunoResearch |

#### Supplementary Table S3 Restriction enzymes used in resection analysis.

| **Localization of the amplified region containing restriction site according to GRCh38/hg38** | **Restriction enzyme** | **Supplier** |
|---|---|---|
| **A** (chr4:3064656+3064725),<br>**H** (chr4:3086756+3086827) | CvikI | New England Biolabs |
| **B** (chr4:3069639+3069741),<br>**G** (chr4:3079948+3080046) | HpyCH4V | New England Biolabs |
| **C** (chr4:3073732+3073872),<br>**E** (chr4:3075078+3075213) | AvaII | New England Biolabs |
| **D** (chr4:3074406+3074528) | HaeIII | New England Biolabs |
| **F** (chr4:3075961+3076086) | RsaI | Thermofisher Scientific |
| **C'** (chr15:47716273+47716371) | MnlI | New England Biolabs |
| **D'** (chr15:47717061+47717216),<br>**F'** (chr15:47718918+47718990) | SfcI | New England Biolabs |
| **E'** (chr15:47717878+47718023) | MluI | Thermofisher Scientific |

#### Supplementary Table S4 Inhibitors and siRNAs used in this study.

| **Inhibited protein** | **Inhibitor** | **Concentration used** | **Supplier** |
|---|---|---|---|
| CTIP | Triapine (3AP) | 0.8 uM | Sigma Aldrich |
| KU70/80 | STL127705 (Compound L) | 7 uM | MedChemExpress |
| MRE11 | Mirin | 25 uM | Sigma Aldrich |
| Polymerase Theta | Novobiocin (NVB) | 25 uM | Selleck Chemicals |
| **Target gene symbol** | **siRNA ID** | **Concentration used** | **Producer** |
| *DCLRE1C* (Artemis) | s532106 | 100 nM | Life Technologies |
| *EXO1* | s17502 | 100 nM | Life Technologies |
| *MRE11* | s8959 | 50 nM | Life Technologies |
| *POLQ* | 122556 | 100 nM | Life Technologies |
| *RAD51* | s531928 | 100 nM | Life Technologies |

**Supplementary Table S5.** The numbers of peptides detected by MS for proteins enriched in gene ontology category of DNA repair. NHEJ: Non-Homologous End Joining; HDR: Homology-Directed Repair; BIR: Break Induced Replication; ICLR: Inter-strand Crosslink Repair; NER: Nucleotide Excision Repair; FACT: Facilitates Chromatin Transcription.

| Proteins | HTT_sgRNA1 | | | | | HTT_sgRNA2 | | | | | Control (unedited) | | | | | Gene ontology category |
|---|---|---|---|---|---|---|---|---|---|---|---|---|---|---|---|---|
| | Exp 1 | Exp 2 | Exp 3 | Exp 4 | Exp 5 | Exp 1 | Exp 2 | Exp 3 | Exp 4 | Exp 5 | Exp 1 | Exp 2 | Exp 3 | Exp 4 | Exp 5 | |
| **BLM** | 3 | 3 | 4 | 3 | 3 | 2 | 2 | 3 | 3 | 2 | 4 | 4 | 4 | 2 | 4 | **HDR** |
| **CDK1** | 12 | 12 | 11 | 9 | 11 | 14 | 14 | 12 | 13 | 13 | 6 | 7 | 6 | 7 | 8 | **Other** |
| **HIST1H4A** | 7 | 7 | 7 | 7 | 7 | 7 | 7 | 7 | 7 | 7 | 5 | 7 | 7 | 5 | 7 | **NHEJ** |
| **HMGA1** | 1 | 2 | 1 | 1 | 1 | 1 | 1 | 1 | 1 | 1 | 3 | 3 | 3 | 3 | 3 | **Other** |
| **HMGA2** | 1 | 2 | 1 | 1 | 2 | 1 | 1 | 1 | 0 | 0 | 2 | 2 | 2 | 2 | 2 | **Other** |
| **HMGB1** | 2 | 2 | 2 | 2 | 3 | 2 | 4 | 2 | 2 | 1 | 2 | 3 | 4 | 4 | 3 | **BER** |
| **HMGB2** | 2 | 1 | 2 | 2 | 2 | 2 | 2 | 2 | 2 | 1 | 2 | 1 | 2 | 2 | 2 | **Other** |
| **MCM3** | 13 | 14 | 12 | 13 | 14 | 15 | 13 | 17 | 15 | 17 | 1 | 4 | 2 | 2 | 2 | **BIR/HDR** |
| **MCM4** | 16 | 19 | 17 | 20 | 19 | 24 | 21 | 22 | 17 | 22 | 1 | 3 | 2 | 1 | 4 | **BIR/HDR** |
| **MCM6** | 12 | 13 | 10 | 14 | 13 | 12 | 14 | 15 | 14 | 20 | 2 | 4 | 5 | 3 | 5 | **BIR/HDR** |
| **MCM7** | 25 | 24 | 23 | 24 | 23 | 26 | 25 | 23 | 25 | 26 | 5 | 9 | 4 | 3 | 12 | **BIR/HDR** |
| **NPM1** | 10 | 11 | 10 | 11 | 10 | 11 | 11 | 10 | 10 | 10 | 7 | 9 | 7 | 7 | 7 | **Other** |
| **PARP1** | 43 | 43 | 45 | 45 | 43 | 46 | 46 | 47 | 43 | 46 | 14 | 27 | 21 | 25 | 21 | **HDR/NER/BER** |
| **PRKDC** | 74 | 84 | 85 | 84 | 82 | 92 | 87 | 89 | 85 | 91 | 20 | 41 | 27 | 23 | 43 | **NHEJ** |
| **RPS27A;UBA52; UBB;UBC** | 8 | 6 | 8 | 5 | 7 | 5 | 5 | 8 | 6 | 5 | 4 | 4 | 7 | 5 | 4 | **ICLR/NER** |
| **SMC1A** | 11 | 14 | 13 | 20 | 16 | 24 | 22 | 23 | 18 | 23 | 2 | 4 | 2 | 1 | 7 | **Cohesin complex** |
| **SSRP1** | 9 | 9 | 10 | 8 | 10 | 11 | 9 | 10 | 10 | 9 | 3 | 3 | 4 | 4 | 3 | **FACT complex** |
| **XRCC5** | 18 | 19 | 17 | 18 | 20 | 21 | 20 | 22 | 20 | 21 | 1 | 8 | 4 | 2 | 4 | **NHEJ** |
| **XRCC6** | 17 | 17 | 17 | 16 | 20 | 14 | 17 | 18 | 17 | 17 | 3 | 8 | 5 | 7 | 2 | **NEHJ** |
| **DDX1** | 13 | 12 | 12 | 11 | 12 | 15 | 14 | 15 | 14 | 15 | 2 | 3 | 1 | 3 | 5 | **Other** |
| **FEN1** | 4 | 4 | 5 | 5 | 5 | 5 | 6 | 5 | 6 | 6 | 1 | 3 | 3 | 2 | 3 | **HDR/BER** |
| **NONO** | 17 | 15 | 16 | 17 | 14 | 18 | 18 | 19 | 18 | 18 | 7 | 10 | 10 | 9 | 10 | **Other** |
| **OTUB1** | 6 | 8 | 7 | 6 | 5 | 8 | 7 | 5 | 7 | 8 | 1 | 3 | 4 | 3 | 2 | **Other** |
| **RPS3** | 18 | 18 | 18 | 18 | 18 | 18 | 18 | 18 | 18 | 18 | 8 | 14 | 11 | 11 | 13 | **Other** |
| **SFPQ** | 15 | 16 | 16 | 13 | 15 | 14 | 15 | 15 | 14 | 16 | 5 | 9 | 6 | 6 | 5 | **HDR** |
| **SMC3** | 18 | 18 | 18 | 17 | 21 | 27 | 21 | 28 | 23 | 25 | 0 | 4 | 3 | 1 | 3 | **Cohesin complex** |
| **SUPT16H** | 21 | 21 | 19 | 18 | 22 | 20 | 22 | 19 | 21 | 17 | 0 | 5 | 5 | 4 | 4 | **FACT complex** |

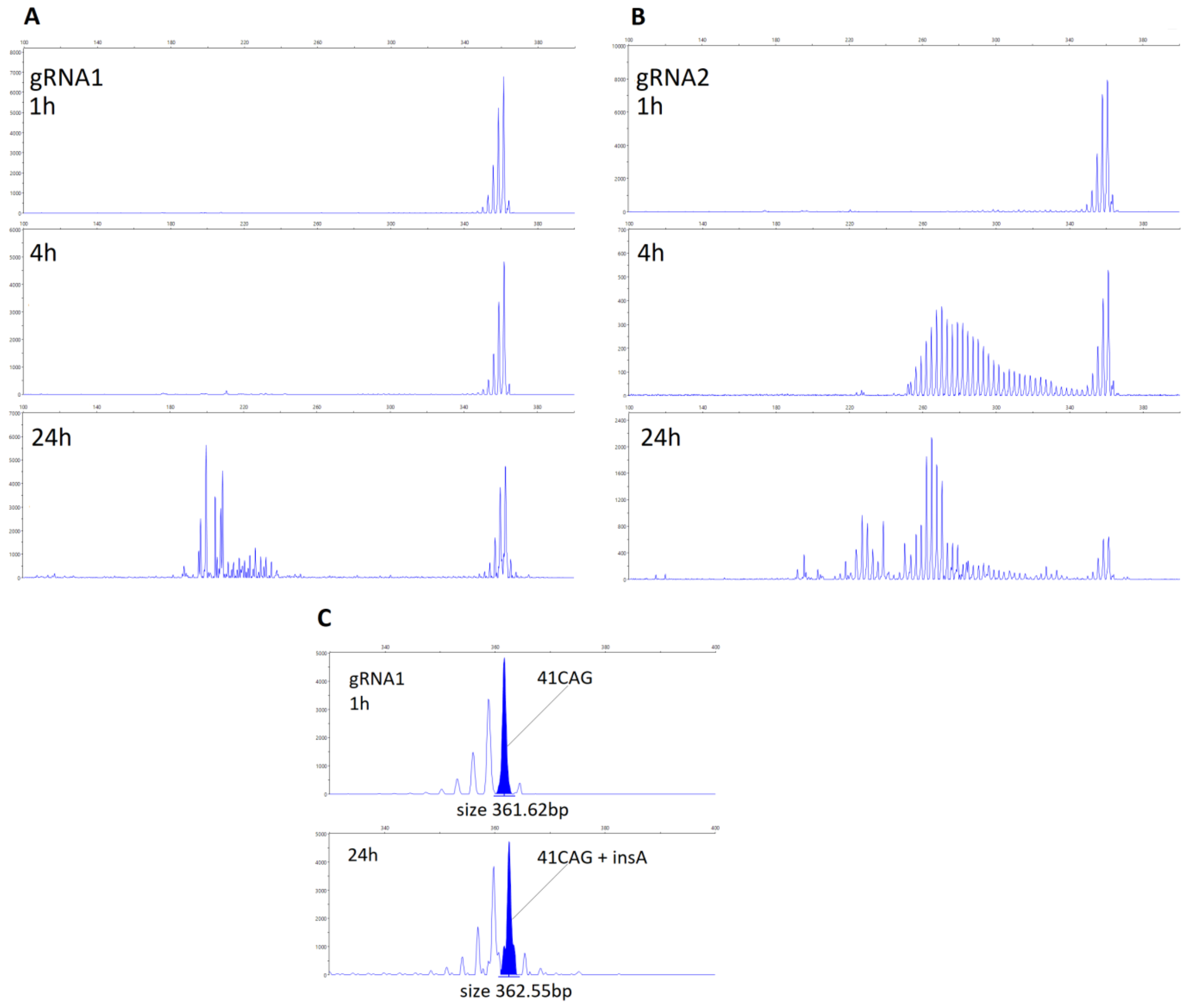


**Supplementary Figure S1**. PCR products of the CAG locus in the HTT gene resolved by capillary electrophoresis. A) Cells were collected at given timepoints after electroporation with RNP containing gRNA1 or B) gRNA2. C) Insertion of A detected 24 h posttransfection in cells edited with gRNA1.

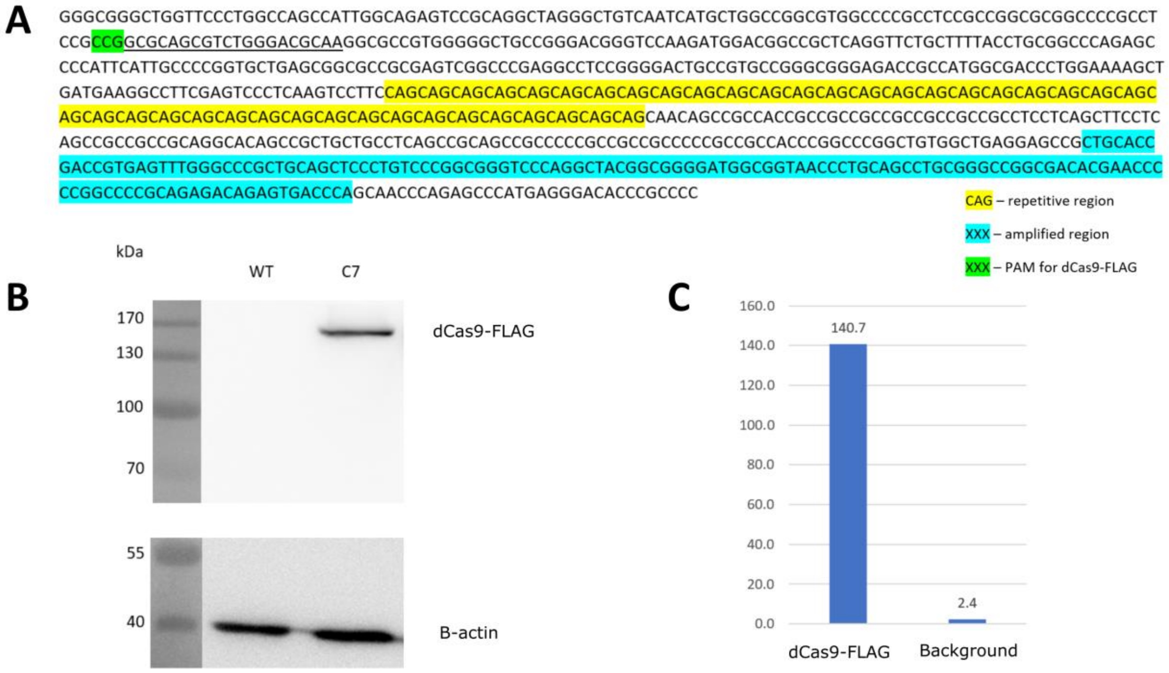


**Supplementary Figure S2.** EnChIP validation. **A)** Sequence of the region containing the CAG repetitive tract in the *HTT* gene. The site complementary to the gRNA guiding dCas9-FLAG is underlined. The PAM sequence is highlighted in green. The sequence amplified during qPCR is highlighted in blue. **B)** Expression of dCas9-FLAG after lentiviral transduction. C7 – HEK293T-41CAG clone expressing dCas9-FLAG. Cell lysates were subjected to Western blot analysis with an anti-Cas9 antibody. β-Actin was used as a loading control. WT – HEK293T-41CAG cells not subjected to lentiviral transduction. **C)** Isolation of the *HTT* CAG repetitive locus by enChIP with dCas9-FLAG. qPCR analysis of enrichment was performed. The locus encoding the β-actin gene was used as a background control.

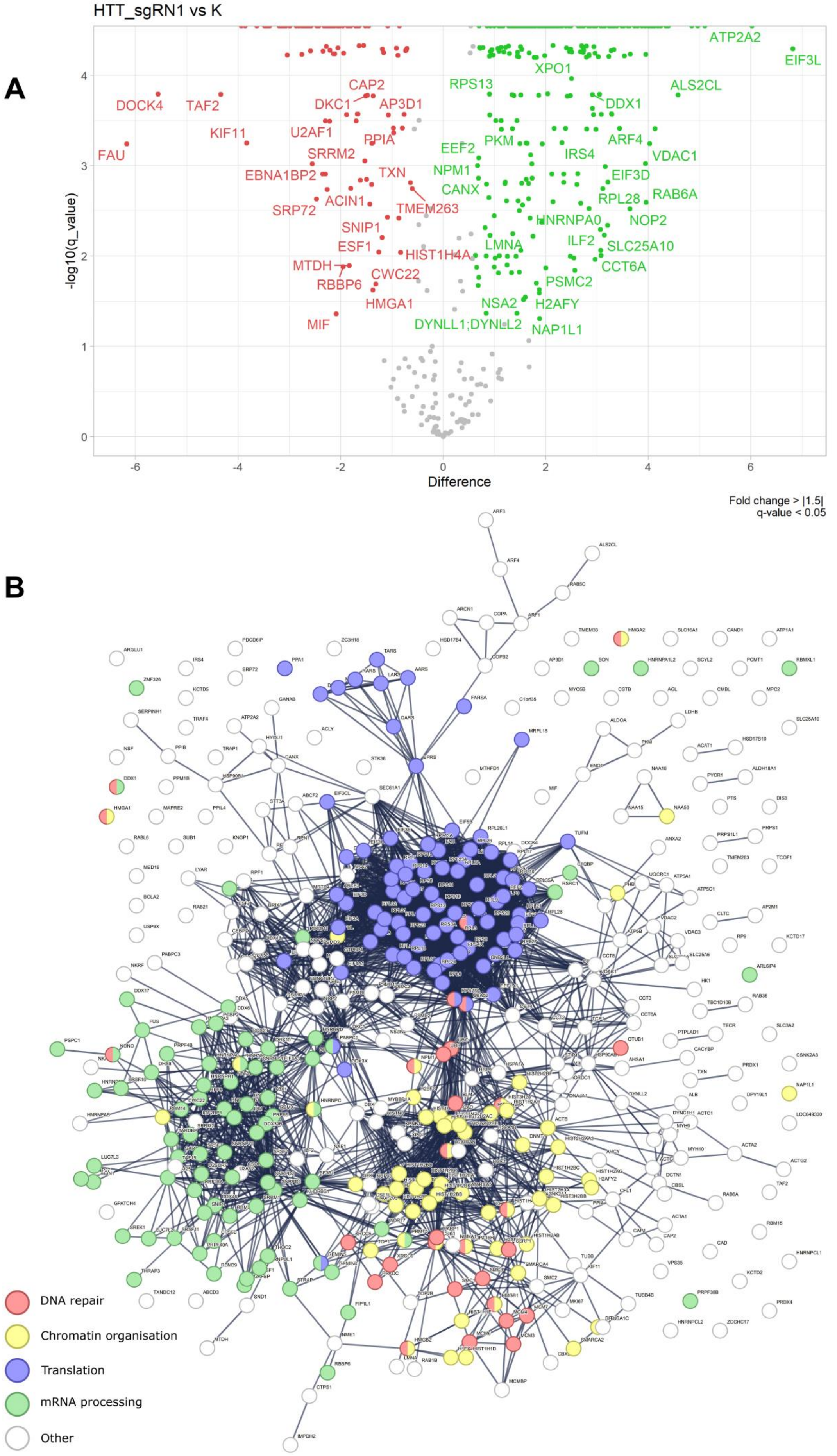
A
HTT_sgRN1 vs K
-log10(q_value)
Difference
Fold change > |1.5|
q-value < 0.05
DOCK4
TAF2
FAU
KIF11
CAP2
DKC1
AP3D1
U2AF1
PPIA
SRRM2
EBNA1BP2
TXN
ACIN1
SRP72
TMEM263
SNIP1
ESF1
HIST1H4A
MTDH
RBBP6
CWC22
HMGA1
MIF
ATP2A2
EIF3L
XPO1
RPS13
ALS2CL
DDX1
PKM
ARF4
EEF2
IRS4
VDAC1
NPM1
EIF3D
CANX
RAB6A
RPL28
NOP2
HNRNPA0
ILF2
LMNA
SLC25A10
CCT6A
PSMC2
NSA2
H2AFY
DYNLL1;DYNLL2
NAP1L1
B
DNA repair
Chromatin organisation
Translation
mRNA processing
Other

**Supplementary Figure S3 A)** Volcano plot presenting the distribution of all the proteins identified in enChIP-MS analysis after introduction of DSBs by using HTT_sgRNA1. The “Difference” parameter is the difference between the mean of log10(protein spectrum intensity) of the experimental groups. The proteins characterized by |fold change| >= 1.5 and q-value < 0.05 were considered significant and are marked in red (downregulated) or green (upregulated). **B)** A STRING network of proteins identified in enChIP-MS analysis after introduction of DSB by using HTT_sgRNA1. Proteins exhibiting low fold change (|fold change| < 1.5) or high q-value (q-value > 0.05) were filtered out. The STRING confidence level was set to the highest (0.900). The proteins that were enriched in the gene ontology category of DNA repair were analysed further (see main text).

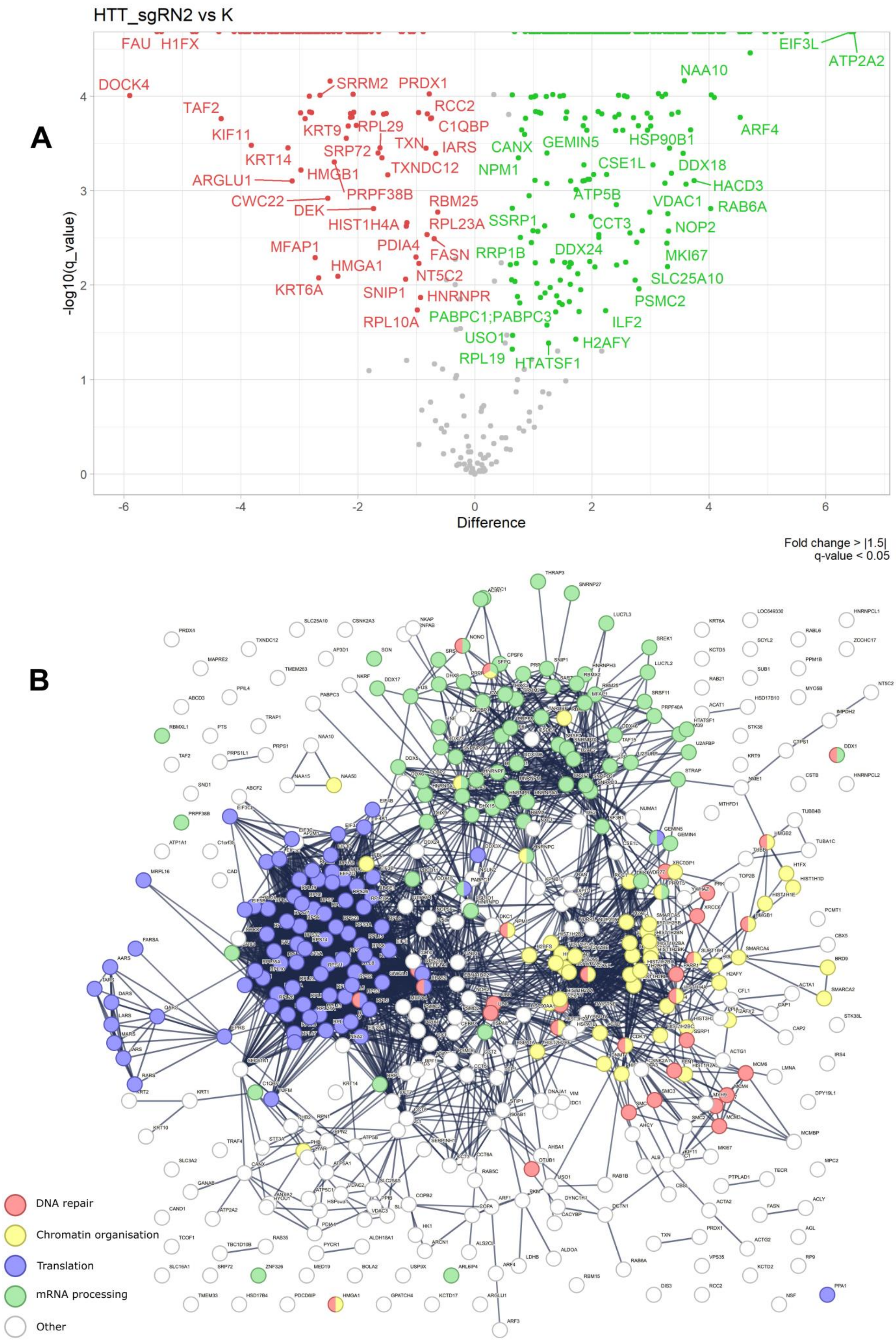
A
HTT_sgRN2 vs K
FAU
H1FX
DOCK4
TAF2
KIF11
KRT14
ARGLU1
CWC22
DEK
MFAP1
KRT6A
HMGA1
SRRM2
PRDX1
RCC2
C1QBP
KRT9
RPL29
TXN
IARS
SRP72
HMGB1
TXNDC12
PRPF38B
RBM25
HIST1H4A
RPL23A
PDIA4
FASN
NT5C2
SNIP1
HNRNPR
RPL10A
EIF3L
ATP2A2
NAA10
ARF4
CANX
GEMIN5
HSP90B1
NPM1
CSE1L
DDX18
HACD3
ATP5B
VDAC1
RAB6A
SSRP1
CCT3
NOP2
RRP1B
DDX24
MKI67
SLC25A10
PSMC2
PABPC1;PABPC3
ILF2
USO1
H2AFY
RPL19
HTATSF1
-log10(q_value)
Difference
Fold change > |1.5|
q-value < 0.05
B
DNA repair
Chromatin organisation
Translation
mRNA processing
Other

**Supplementary Figure S4 A)** Volcano plot presenting the distribution of all the proteins identified in enChIP-MS analysis after introduction of DSBs by using HTT_sgRNA2. The "Difference" parameter is the difference between the mean of log10(protein spectrum intensity) of the experimental groups. The proteins characterized by |fold change| >= 1.5 and q-value < 0.05 were considered significant and are marked in red (decreased levels) or green (increased levels). **B)** A STRING network of proteins identified in enChIP-MS analysis after introduction of DSB by using HTT_sgRNA2. Proteins exhibiting low fold change (|fold change| < 1.5) or high q-value (q-value > 0.05) were filtered out. The STRING confidence level was set to the highest (0.900). The proteins that were enriched in the gene ontology category of DNA repair were analysed further (see main text).

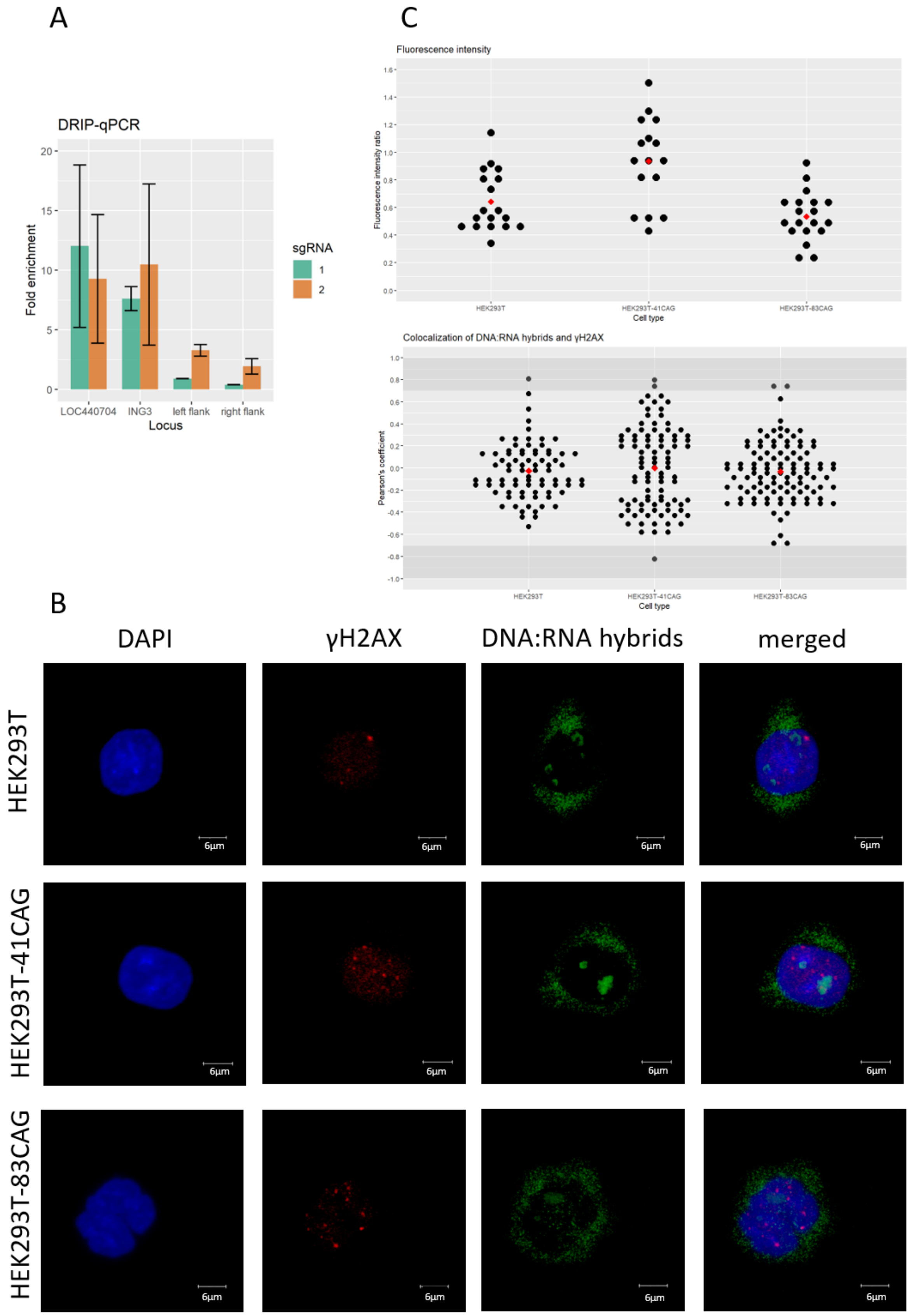
A
DRIP-qPCR
Fold enrichment
Locus
LOC440704
ING3
left flank
right flank
sgRNA
1
2
C
Fluorescence intensity
Fluorescence intensity ratio
Cell type
HEK293T
HEK293T-41CAG
HEK293T-83CAG
Colocalization of DNA:RNA hybrids and γH2AX
Pearson's coefficient
B
DAPI
γH2AX
DNA:RNA hybrids
merged
HEK293T
HEK293T-41CAG
HEK293T-83CAG
6μm

**Supplementary Figure S5.** Presence of DNA:RNA hybrids in the process of DSB repair at the *HTT* locus. A) The enrichment of DNA:RNA hybrids determined by DRIP-qPCR in the positive control locus (LOC440704 and ING3) and in the regions flanking the CAG repeat tract in the *HTT* locus after induction of DSBs using HTT_sgRNA1 and HTT_sgRNA2, respectively. DNA:RNA hybrids are 2-fold enriched in the *HTT* locus after inducing DSBs with HTT_sgRNA2. No significant enrichment of DNA:RNA hybrids was observed after inducing DSBs with HTT_sgRNA1. The data shown represent the mean ± SD (n=2). B) Representative images of nuclei of HEK293T, HEK293T-41CAG and HEK293T-83CAG cells after inducing DSBs with HTT_gRNA2 with foci of γH2AX (red) and RNA:DNA hybrids (green). Scale bar = 6 µm. C) Graphs representing Pearson's coefficient for colocalization of γH2AX and RNA:DNA hybrid foci (n=79, 94, and 105, respectively) and the ratio of the fluorescence intensity from DNA:RNA hybrids to the fluorescence from γH2AX (n=18, 16, and 19, respectively) within nuclei of HEK293T, HEK293T-41CAG and HEK293T-83CAG cells.
