## Supplemental file 1 for "CRISPR/Cas9-induced double-strand breaks in huntingtin locus lead to CAG repeat contraction through the extensive DNA end resection and homology-mediated repair"

| T#ID | CAG | WT Sequence | RGEN Treated Sequence | Length | Count | Type |
|---|---|---|---|---|---|---|
| 109 | 40.0 | GCTGATGAAGGCCTTCGAGTCCCT**CAAGTCCTTCC**-AGCAGCAGCAGCAGCAGCAGCAGCAGCAGCAGCAGCAGCAGCAGCAGCAGCAGCAGCAGCAGCAGCAGCAGCAGCAGCAGCAGCAGCAGCAGCAGCAGCAGCAGCAGCAGCAGCAGCAGCAGCAGCAG**CAACAGCCG**CCACCGCCGCCGCCGCCGCCGCCGCCTCCTCAGCTTCCTCAGCCGCCGCCGCAGGCACAGCCGCTGCTGCCTCAGC | GCTGATGAAGGCCTTCGAGTCCCT**CAAGTCCTTCC**AAGCAGCAGCAGCAGCAGCAGCAGCAGCAGCAGCAGCAGCAGCAGCAGCAGCAGCAGCAGCAGCAGCAGCAGCAGCAGCAGCAGCAGCAGCAGCAGCAGCAGCAGCAGCAGCAGCAGCAGCAGCAGCAG**CAACAGCCG**CCACCGCCGCCGCCGCCGCCGCCGCCTCCTCAGCTTCCTCAGCCGCCGCCGCAGGCACAGCCGCTGCTGCCTCAGC | 242 | 23396 | INS |
| 125 | 39.0 | GCTGATGAAGGCCTTCGAGTCCCT**CAAGTCCTTCC**AGCAGCAGCAGCAGCAGCAGCAGCAGCAGCAGCAGCAGCAGCAGCAGCAGCAGCAGCAGCAGCAGCAGCAGCAGCAGCAGCAGCAGCAGCAGCAGCAGCAGCAGCAGCAGCAGCAGCAGCAGCAGCAG**CAACAGCCG**CCACCGCCGCCGCCGCCGCCGCCGCCTCCTCAGCTTCCTCAGCCGCCGCCGCAGGCACAGCCGCTGCTGCCTCAGC | GCTGATGAAGGCCTTCGAGTCCCT**CAAGTCCTTCC**A--AGCAGCAGCAGCAGCAGCAGCAGCAGCAGCAGCAGCAGCAGCAGCAGCAGCAGCAGCAGCAGCAGCAGCAGCAGCAGCAGCAGCAGCAGCAGCAGCAGCAGCAGCAGCAGCAGCAGCAGCAG**CAACAGCCG**CCACCGCCGCCGCCGCCGCCGCCGCCTCCTCAGCTTCCTCAGCCGCCGCCGCAGGCACAGCCGCTGCTGCCTCAGC | 242 | 15100 | DEL |
| 1300 | 0.0 | GCTGATGAAGGCCTTCGAGTCCCT**CAAGTCCTTCC**AGCAGCAGCAGCAGCAGCAGCAGCAGCAGCAGCAGCAGCAGCAGCAGCAGCAGCAGCAGCAGCAGCAGCAGCAGCAGCAGCAGCAGCAGCAGCAGCAGCAGCAGCAGCAGCAGCAGCAGCAGCAGCAG**CAACAGCCG**CCACCGCCGCCGCCGCCGCCGCCGCCTCCTCAGCTTCCTCAGCCGCCGCCGCAGGCACAGCCGCTGCTGCCTCAGC | GCTGATGAAGGCCTTCGAGTCCCTCAAGTCCTT----------------------------------------------------------------------------------------------------------------------------------------------CCGCCGCCGCCGCCGCCTCCTCAGCTTCCTCAGCCGCCGCCGCAGGCACAGCCGCTGCTGCCTCAGC | 242 | 8849 | DEL |
| 1335 | 0.0 | GCTGATGAAGGCCTTCGAGTCCCT**CAAGTCCTTCC**AGCAGCAGCAGCAGCAGCAGCAGCAGCAGCAGCAGCAGCAGCAGCAGCAGCAGCAGCAGCAGCAGCAGCAGCAGCAGCAGCAGCAGCAGCAGCAGCAGCAGCAGCAGCAGCAGCAGCAGCAGCAGCAG**CAACAGCCG**CCACCGCCGCCGCCGCCGCCGCCGCCTCCTCAGCTTCCTCAGCCGCCGCCGCAGGCACAGCCGCTGCTGCCTCAGC | GCTGATGAAGGCCTTCGAGTCCCTCAAGT--------------------------------------------------------------------------------------------------------------------------------------------------CCGCCGCCGCCGCCGCCTCCTCAGCTTCCTCAGCCGCCGCCGCAGGCACAGCCGCTGCTGCCTCAGC | 242 | 7905 | DEL |
| 1351 | 0.0 | GCTGATGAAGGCCTTCGAGTCCCT**CAAGTCCTTCC**AGCAGCAGCAGCAGCAGCAGCAGCAGCAGCAGCAGCAGCAGCAGCAGCAGCAGCAGCAGCAGCAGCAGCAGCAGCAGCAGCAGCAGCAGCAGCAGCAGCAGCAGCAGCAGCAGCAGCAGCAGCAGCAG**CAACAGCCG**CCACCGCCGCCGCCGCCGCCGCCGCCTCCTCAGCTTCCTCAGCCGCCGCCGCAGGCACAGCCGCTGCTGCCTCAGC | GCTGATGAAGGCCTTCGAGTCCCTCAAGTCCTT----------------------------------------------------------------------------------------------------------------------------------------------------CCGGCGCCGCCTCCTCAGCTTCCTCAGCCGCCGCCGCAGGCACAGCCGCTGCTGCCTCAGC | 242 | 7867 | DEL |
| 1492 | 0.0 | GCTGATGAAGGCCTTCGAGTCCCT**CAAGTCCTTCC**AGCAGCAGCAGCAGCAGCAGCAGCAGCAGCAGCAGCAGCAGCAGCAGCAGCAGCAGCAGCAGCAGCAGCAGCAGCAGCAGCAGCAGCAGCAGCAGCAGCAGCAGCAGCAGCAGCAGCAGCAGCAGCAG**CAACAGCCG**CCACCGCCGCCGCCGCCGCCGCCGCCTCCTCAGCTTCCTCAGCCGCCGCCGCAGGCACAGCCGCTGCTGCCTCAGC | GCTGATGAAGGCCTTCGAGTC----------------------------------------------------------------------------------------------------------------------------------------------------------------------------CCTCAGCTTCCTCAGCCGCCGCCGCAGGCACAGCCGCTGCTGCCTCAGC | 242 | 7625 | DEL |
| 1243 | 0.0 | GCTGATGAAGGCCTTCGAGTCCCT**CAAGTCCTTCC**AGCAGCAGCAGCAGCAGCAGCAGCAGCAGCAGCAGCAGCAGCAGCAGCAGCAGCAGCAGCAGCAGCAGCAGCAGCAGCAGCAGCAGCAGCAGCAGCAGCAGCAGCAGCAGCAGCAGCAGCAGCAGCAG**CAACAGCCG**CCACCGCCGCCGCCGCCGCCGCCGCCTCCTCAGCTTCCTCAGCCGCCGCCGCAGGCACAGCCGCTGCTGCCTCAGC | GCTGATGAAGGCCTTCGAGTCCCTCAAGTCCTT----------------------------------------------------------------------------------------------------------------------------------------CCGCCGCCGCCGCCGCCGCCGCCTCCTCAGCTTCCTCAGCCGCCGCCGCAGGCACAGCCGCTGCTGCCTCAGC | 242 | 6340 | DEL |
| 1214 | 0.0 | GCTGATGAAGGCCTTCGAGTCCCT**CAAGTCCTTCC**AGCAGCAGCAGCAGCAGCAGCAGCAGCAGCAGCAGCAGCAGCAGCAGCAGCAGCAGCAGCAGCAGCAGCAGCAGCAGCAGCAGCAGCAGCAGCAGCAGCAGCAGCAGCAGCAGCAGCAGCAGCAGCAG**CAACAGCCG**CCACCGCCGCCGCCGCCGCCGCCGCCTCCTCAGCTTCCTCAGCCGCCGCCGCAGGCACAGCCGCTGCTGCCTCAGC | GCTGATGAAGGCCTTCGAGTCCCTCAAGTCCTT-------------------------------------------------------------------------------------------------------------------------------------CCACCGCCGCCGCCGCCGCCGCCGCCTCCTCAGCTTCCTCAGCCGCCGCCGCAGGCACAGCCGCTGCTGCCTCAGC | 242 | 6280 | DEL |
| 1256 | 0.0 | GCTGATGAAGGCCTTCGAGTCCCT**CAAGTCCTTCC**AGCAGCAGCAGCAGCAGCAGCAGCAGCAGCAGCAGCAGCAGCAGCAGCAGCAGCAGCAGCAGCAGCAGCAGCAGCAGCAGCAGCAGCAGCAGCAGCAGCAGCAGCAGCAGCAGCAGCAGCAGCAGCAG**CAACAGCCG**CCACCGCCGCCGCCGCCGCCGCCGCCTCCTCAGCTTCCTCAGCCGCCGCCGCAGGCACAGCCGCTGCTGCCTCAGC | GCTGATGAAGGCCTTCGAGTCCCTCAAGT-----------------------------------------------------------------------------------------------------------------------------------------CCACCGCCGCCGCCGCCGCCGCCGCCTCCTCAGCTTCCTCAGCCGCCGCCGCAGGCACAGCCGCTGCTGCCTCAGC | 242 | 6097 | DEL |
| 168 | 38.0 | GCTGATGAAGGCCTTCGAGTCCCT**CAAGTCCTTCC**AGCAGCAGCAGCAGCAGCAGCAGCAGCAGCAGCAGCAGCAGCAGCAGCAGCAGCAGCAGCAGCAGCAGCAGCAGCAGCAGCAGCAGCAGCAGCAGCAGCAGCAGCAGCAGCAGCAGCAGCAGCAGCAG**CAACAGCCG**CCACCGCCGCCGCCGCCGCCGCCGCCTCCTCAGCTTCCTCAGCCGCCGCCGCAGGCACAGCCGCTGCTGCCTCAGC | GCTGATGAAGGCCTTCGAGTCCCT**CAAGTCCTTCC**A-----AGCAGCAGCAGCAGCAGCAGCAGCAGCAGCAGCAGCAGCAGCAGCAGCAGCAGCAGCAGCAGCAGCAGCAGCAGCAGCAGCAGCAGCAGCAGCAGCAGCAGCAGCAGCAGCAGCAG**CAACAGCCG**CCACCGCCGCCGCCGCCGCCGCCGCCTCCTCAGCTTCCTCAGCCGCCGCCGCAGGCACAGCCGCTGCTGCCTCAGC | 242 | 5817 | DEL |
| 116 | 41.0 | GCTGATGAAGGCCTTCGAGTCCCT**CAAGTCCTTCC**AGCAGCAGCAGCAGCAGCAGCAGCAGCAGCAGCAGCAGCAGCAGCAGCAGCAGCAGCAGCAGCAGCAGCAGCAGCAGCAGCAGCAGCAGCAGCAGCAGCAGCAGCAGCAGCAGCAGCAGCAGCAGCAG**CAACAGCCG**CCACCGCCGCCGCCGCCGCCGCCGCCTCCTCAGCTTCCTCAGCCGCCGCCGCAGGCACAGCCGCTGCTGCCTCAGC | GCTGATGAAGGCCTTCGAGTCCCT**CAAGTCCTTCC**AGCAGCAGCAGCAGCAGCAGCAGCAGCAGCAGCAGCAGCAGCAGCAGCAGCAGCAGCAGCAGCAGCAGCAGCAGCAGCAGCAGCAGCAGCAGCAGCAGCAGCAGCAGCAGCAGCAGCAGCAGCAGCAG**CAACAGCCG**CCACCGCCGCCGCCGCCGCCGCCGCCTCCTCAGCTTCCTCAGCCGCCGCCGCAGGCACAGCCGCTGCTGCCTCAGC | 242 | 5776 | WT/SUB |
| 1276 | 0.0 | GCTGATGAAGGCCTTCGAGTCCCT**CAAGTCCTTCC**AGCAGCAGCAGCAGCAGCAGCAGCAGCAGCAGCAGCAGCAGCAGCAGCAGCAGCAGCAGCAGCAGCAGCAGCAGCAGCAGCAGCAGCAGCAGCAGCAGCAGCAGCAGCAGCAGCAGCAGCAGCAGCAG**CAACAGCCG**CCACCGCCGCCGCCGCCGCCGCCGCCTCCTCAGCTTCCTCAGCCGCCGCCGCAGGCACAGCCGCTGCTGCCTCAGC | GCTGATGAAGGCCTTCGAGTCCCTCAAGTCCTT-------------------------------------------------------------------------------------------------------------------------------------------CCGCCGCCGCCGCCGCCGCCTCCTCAGCTTCCTCAGCCGCCGCCGCAGGCACAGCCGCTGCTGCCTCAGC | 242 | 5326 | DEL |
| 1325 | 0.0 | GCTGATGAAGGCCTTCGAGTCCCT**CAAGTCCTTCC**AGCAGCAGCAGCAGCAGCAGCAGCAGCAGCAGCAGCAGCAGCAGCAGCAGCAGCAGCAGCAGCAGCAGCAGCAGCAGCAGCAGCAGCAGCAGCAGCAGCAGCAGCAGCAGCAGCAGCAGCAGCAGCAG**CAACAGCCG**CCACCGCCGCCGCCGCCGCCGCCGCCTCCTCAGCTTCCTCAGCCGCCGCCGCAGGCACAGCCGCTGCTGCCTCAGC | GCTGATGAAGGCCTTCGAGTCCCTCAAGTCCTT-------------------------------------------------------------------------------------------------------------------------------------------------CCGCCGCCGCCGCCTCCTCAGCTTCCTCAGCCGCCGCCGCAGGCACAGCCGCTGCTGCCTCAGC | 242 | 5079 | DEL |
| 1307 | 0.0 | GCTGATGAAGGCCTTCGAGTCCCT**CAAGTCCTTCC**AGCAGCAGCAGCAGCAGCAGCAGCAGCAGCAGCAGCAGCAGCAGCAGCAGCAGCAGCAGCAGCAGCAGCAGCAGCAGCAGCAGCAGCAGCAGCAGCAGCAGCAGCAGCAGCAGCAGCAGCAGCAGCAG**CAACAGCCG**CCACCGCCGCCGCCGCCGCCGCCGCCTCCTCAGCTTCCTCAGCCGCCGCCGCAGGCACAGCCGCTGCTGCCTCAGC | GCTGATGAAGGCCTTCGAGTCCCTCAAGTC-----------------------------------------------------------------------------------------------------------------------------------------------TCAAGCCGCCGCCGCCGCCTCCTCAGCTTCCTCAGCCGCCGCCGCAGGCACAGCCGCTGCTGCCTCAGC | 242 | 4892 | DEL |
| 1455 | 0.0 | GCTGATGAAGGCCTTCGAGTCCCT**CAAGTCCTTCC**AGCAGCAGCAGCAGCAGCAGCAGCAGCAGCAGCAGCAGCAGCAGCAGCAGCAGCAGCAGCAGCAGCAGCAGCAGCAGCAGCAGCAGCAGCAGCAGCAGCAGCAGCAGCAGCAGCAGCAGCAGCAGCAG**CAACAGCCG**CCACCGCCGCCGCCGCCGCCGCCGCCTCCTCAGCTTCCTCAGCCGCCGCCGCAGGCACAGCCGCTGCTGCCTCAGC | GCTGATGAAGGCCTTCGAGTCCCTCAAG--------------------------------------------------------------------------------------------------------------------------------------------------------------------TCCTCAGCTTCCTCAGCCGCCGCCGCAGGCACAGCCGCTGCTGCCTCAGC | 242 | 4424 | DEL |
| 1285 | 0.0 | GCTGATGAAGGCCTTCGAGTCCCT**CAAGTCCTTCC**AGCAGCAGCAGCAGCAGCAGCAGCAGCAGCAGCAGCAGCAGCAGCAGCAGCAGCAGCAGCAGCAGCAGCAGCAGCAGCAGCAGCAGCAGCAGCAGCAGCAGCAGCAGCAGCAGCAGCAGCAGCAGCAG**CAACAGCCG**CCACCGCCGCCGCCGCCGCCGCCGCCTCCTCAGCTTCCTCAGCCGCCGCCGCAGGCACAGCCGCTGCTGCCTCAGC | GCTGATGAAGGCCTTCGAGTCCCTCAAGT--------------------------------------------------------------------------------------------------------------------------------------------CCGCCGCCGCCGCCGCCGCCGCCTCCTCAGCTTCCTCAGCCGCCGCCGCAGGCACAGCCGCTGCTGCCTCAGC | 242 | 4176 | DEL |
| 1501 | 0.0 | GCTGATGAAGGCCTTCGAGTCCCT**CAAGTCCTTCC**AGCAGCAGCAGCAGCAGCAGCAGCAGCAGCAGCAGCAGCAGCAGCAGCAGCAGCAGCAGCAGCAGCAGCAGCAGCAGCAGCAGCAGCAGCAGCAGCAGCAGCAGCAGCAGCAGCAGCAGCAGCAGCAG**CAACAGCCG**CCACCGCCGCCGCCGCCGCCGCCGCCTCCTCAGCTTCCTCAGCCGCCGCCGCAGGCACAGCCGCTGCTGCCTCAGC | GCTGATGAAGGCCTTCGAGTCCCTCAAG-----------------------------------------------------------------------------------------------------------------------------------------------------------------------------TCCTCAGCCGCCGCCGCAGGCACAGCCGCTGCTGCCTCAGC | 242 | 3928 | DEL |
| 1220 | 0.0 | GCTGATGAAGGCCTTCGAGTCCCT**CAAGTCCTTCC**AGCAGCAGCAGCAGCAGCAGCAGCAGCAGCAGCAGCAGCAGCAGCAGCAGCAGCAGCAGCAGCAGCAGCAGCAGCAGCAGCAGCAGCAGCAGCAGCAGCAGCAGCAGCAGCAGCAGCAGCAGCAGCAG**CAACAGCCG**CCACCGCCGCCGCCGCCGCCGCCGCCTCCTCAGCTTCCTCAGCCGCCGCCGCAGGCACAGCCGCTGCTGCCTCAGC | GCTGATGAAGGCCTTCGAGTCCCTCAAGT--------------------------------------------------------------------------------------------------------------------------------------CCGCCACCGCCGCCGCCGCCGCCGCCGCCTCCTCAGCTTCCTCAGCCGCCGCCGCAGGCACAGCCGCTGCTGCCTCAGC | 242 | 3466 | DEL |
| 134 | 40.0 | GCTGATGAAGGCCTTCGAGTCCCT**CAAGTCCTTCC**AGCAGCAGCAGCAGCAGCAGCAGCAGCAGCAGCAGCAGCAGCAGCAGCAGCAGCAGCAGCAGCAGCAGCAGCAGCAGCAGCAGCAGCAGCAGCAGCAGCAGCAGCAGCAGCAGCAGCAGCAGCAGCAG**CAACAGCCG**CCACCGCCGCCGCCGCCGCCGCCGCCTCCTCAGCTTCCTCAGCCGCCGCCGCAGGCACAGCCGCTGCTGCCTCAGC | GCTGATGAAGGCCTTCGAGTCCCTCAAGTCCTTC---CAGCAGCAGCAGCAGCAGCAGCAGCAGCAGCAGCAGCAGCAGCAGCAGCAGCAGCAGCAGCAGCAGCAGCAGCAGCAGCAGCAGCAGCAGCAGCAGCAGCAGCAGCAGCAGCAGCAGCAGCAGCAG**CAACAGCCG**CCACCGCCGCCGCCGCCGCCGCCGCCTCCTCAGCTTCCTCAGCCGCCGCCGCAGGCACAGCCGCTGCTGCCTCAGC | 242 | 3211 | DEL |
| 92 | 41.0 | GCTGATGAAGGCCTTCGAGTCCCTCAAGTCCTTC----CAGCAGCAGCAGCAGCAGCAGCAGCAGCAGCAGCAGCAGCAGCAGCAGCAGCAGCAGCAGCAGCAGCAGCAGCAGCAGCAGCAGCAGCAGCAGCAGCAGCAGCAGCAGCAGCAGCAGCAGCAGCAGCAGCAG**CAACAGCCG**CCACCGCCGCCGCCGCCGCCGCCGCCTCCTCAGCTTCCTCAGCCGCCGCCGCAGGCACAGCCGCTGCTGCCTCAGC | GCTGATGAAGGCCTTCGAGTCCCT**CAAGTCCTTCC**AAGCAGCAGCAGCAGCAGCAGCAGCAGCAGCAGCAGCAGCAGCAGCAGCAGCAGCAGCAGCAGCAGCAGCAGCAGCAGCAGCAGCAGCAGCAGCAGCAGCAGCAGCAGCAGCAGCAGCAGCAGCAGCAGCAG**CAACAGCCG**CCACCGCCGCCGCCGCCGCCGCCGCCTCCTCAGCTTCCTCAGCCGCCGCCGCAGGCACAGCCGCTGCTGCCTCAGC | 242 | 3076 | INS |
| 1359 | 0.0 | GCTGATGAAGGCCTTCGAGTCCCT**CAAGTCCTTCC**AGCAGCAGCAGCAGCAGCAGCAGCAGCAGCAGCAGCAGCAGCAGCAGCAGCAGCAGCAGCAGCAGCAGCAGCAGCAGCAGCAGCAGCAGCAGCAGCAGCAGCAGCAGCAGCAGCAGCAGCAGCAGCAG**CAACAGCCG**CCACCGCCGCCGCCGCCGCCGCCGCCTCCTCAGCTTCCTCAGCCGCCGCCGCAGGCACAGCCGCTGCTGCCTCAGC | GCTGATGAAGGCCTTCGAGTCCCTCAAGT-----------------------------------------------------------------------------------------------------------------------------------------------------CCGCCGCCGCCGCCTCCTCAGCTTCCTCAGCCGCCGCCGCAGGCACAGCCGCTGCTGCCTCAGC | 242 | 3068 | DEL |
| 1233 | 0.0 | GCTGATGAAGGCCTTCGAGTCCCT**CAAGTCCTTCC**AGCAGCAGCAGCAGCAGCAGCAGCAGCAGCAGCAGCAGCAGCAGCAGCAGCAGCAGCAGCAGCAGCAGCAGCAGCAGCAGCAGCAGCAGCAGCAGCAGCAGCAGCAGCAGCAGCAGCAGCAGCAGCAG**CAACAGCCG**CCACCGCCGCCGCCGCCGCCGCCGCCTCCTCAGCTTCCTCAGCCGCCGCCGCAGGCACAGCCGCTGCTGCCTCAGC | GCTGATGAAGGCCTTCGAGTCCCTCA---------------------------------------------------------------------------------------------------------------------------------------AGCCGCCACCGCCGCCGCCGCCGCCGCCGCCTCCTCAGCTTCCTCAGCCGCCGCCGCAGGCACAGCCGCTGCTGCCTCAGC | 242 | 3048 | DEL |
| 1452 | 0.0 | GCTGATGAAGGCCTTCGAGTCCCT**CAAGTCCTTCC**AGCAGCAGCAGCAGCAGCAGCAGCAGCAGCAGCAGCAGCAGCAGCAGCAGCAGCAGCAGCAGCAGCAGCAGCAGCAGCAGCAGCAGCAGCAGCAGCAGCAGCAGCAGCAGCAGCAGCAGCAGCAGCAG**CAACAGCCG**CCACCGCCGCCGCCGCCGCCGCCGCCTCCTCAGCTTCCTCAGCCGCCGCCGCAGGCACAGCCGCTGCTGCCTCAGC | GCTGATGAAGGCCTTCGAGTCCCTCAAGTC-------------------------------------------------------------------------------------------------------------------------------------------------------------------CCTCAGCTTCCTCAGCCGCCGCCGCAGGCACAGCCGCTGCTGCCTCAGC | 242 | 2953 | DEL |
| 1366 | 0.0 | GCTGATGAAGGCCTTCGAGTCCCT**CAAGTCCTTCC**AGCAGCAGCAGCAGCAGCAGCAGCAGCAGCAGCAGCAGCAGCAGCAGCAGCAGCAGCAGCAGCAGCAGCAGCAGCAGCAGCAGCAGCAGCAGCAGCAGCAGCAGCAGCAGCAGCAGCAGCAGCAGCAG**CAACAGCCG**CCACCGCCGCCGCCGCCGCCGCCGCCTCCTCAGCTTCCTCAGCCGCCGCCGCAGGCACAGCCGCTGCTGCCTCAGC | GCTGATGAAGGCCTTCGAGTCCCTCAAGT------------------------------------------------------------------------------------------------------------------------------------------------------CGCCGCCGCCGCCTCCTCAGCTTCCTCAGCCGCCGCCGCAGGCACAGCCGCTGCTGCCTCAGC | 242 | 2950 | DEL |
| 1443 | 0.0 | GCTGATGAAGGCCTTCGAGTCCCT**CAAGTCCTTCC**AGCAGCAGCAGCAGCAGCAGCAGCAGCAGCAGCAGCAGCAGCAGCAGCAGCAGCAGCAGCAGCAGCAGCAGCAGCAGCAGCAGCAGCAGCAGCAGCAGCAGCAGCAGCAGCAGCAGCAGCAGCAGCAG**CAACAGCCG**CCACCGCCGCCGCCGCCGCCGCCGCCTCCTCAGCTTCCTCAGCCGCCGCCGCAGGCACAGCCGCTGCTGCCTCAGC | GCTGATGAAGGCCTTCGAGTCCCT**CAAGTCCTTCC**-----------------------------------------------------------------------------------------------------------------------------------------------------------------CTGCTTCCTCAGCCGCCGCCGCAGGCACAGCCGCTGCTGCCTCAGC | 242 | 2674 | DEL |
| 231 | 37.0 | GCTGATGAAGGCCTTCGAGTCCCT**CAAGTCCTTCC**AGCAGCAGCAGCAGCAGCAGCAGCAGCAGCAGCAGCAGCAGCAGCAGCAGCAGCAGCAGCAGCAGCAGCAGCAGCAGCAGCAGCAGCAGCAGCAGCAGCAGCAGCAGCAGCAGCAGCAGCAGCAGCAG**CAACAGCCG**CCACCGCCGCCGCCGCCGCCGCCGCCTCCTCAGCTTCCTCAGCCGCCGCCGCAGGCACAGCCGCTGCTGCCTCAGC | GCTGATGAAGGCCTTCGAGTCCCT**CAAGTCCTTCC**A--------AGCAGCAGCAGCAGCAGCAGCAGCAGCAGCAGCAGCAGCAGCAGCAGCAGCAGCAGCAGCAGCAGCAGCAGCAGCAGCAGCAGCAGCAGCAGCAGCAGCAGCAGCAGCAG**CAACAGCCG**CCACCGCCGCCGCCGCCGCCGCCGCCTCCTCAGCTTCCTCAGCCGCCGCCGCAGGCACAGCCGCTGCTGCCTCAGC | 242 | 2324 | DEL |
| 1266 | 0.0 | GCTGATGAAGGCCTTCGAGTCCCT**CAAGTCCTTCC**AGCAGCAGCAGCAGCAGCAGCAGCAGCAGCAGCAGCAGCAGCAGCAGCAGCAGCAGCAGCAGCAGCAGCAGCAGCAGCAGCAGCAGCAGCAGCAGCAGCAGCAGCAGCAGCAGCAGCAGCAGCAGCAG**CAACAGCCG**CCACCGCCGCCGCCGCCGCCGCCGCCTCCTCAGCTTCCTCAGCCGCCGCCGCAGGCACAGCCGCTGCTGCCTCAGC | GCTGATGAAGGCCTTCGAGTCCCT**CAAGTCCTTCC**------------------------------------------------------------------------------------------------------------------------------------------GACTCGCGGCGCCGCCGCCTCCTCAGCTTCCTCAGCCGCCGCCGCAGGCACAGCCGCTGCTGCCTCAGC | 242 | 2254 | DEL |
| 1387 | 0.0 | GCTGATGAAGGCCTTCGAGTCCCT**CAAGTCCTTCC**AGCAGCAGCAGCAGCAGCAGCAGCAGCAGCAGCAGCAGCAGCAGCAGCAGCAGCAGCAGCAGCAGCAGCAGCAGCAGCAGCAGCAGCAGCAGCAGCAGCAGCAGCAGCAGCAGCAGCAGCAGCAGCAG**CAACAGCCG**CCACCGCCGCCGCCGCCGCCGCCGCCTCCTCAGCTTCCTCAGCCGCCGCCGCAGGCACAGCCGCTGCTGCCTCAGC | GCTGATGAAGGCCTTCGAGTCCCTCAAGT---------------------------------------------------------------------------------------------------------------------------------------------------------CGCCGCCGCCTCCTCAGCTTCCTCAGCCGCCGCCGCAGGCACAGCCGCTGCTGCCTCAGC | 242 | 2134 | DEL |
| 1108 | 0.0 | GCTGATGAAGGCCTTCGAGTCCCT**CAAGTCCTTCC**AGCAGCAGCAGCAGCAGCAGCAGCAGCAGCAGCAGCAGCAGCAGCAGCAGCAGCAGCAGCAGCAGCAGCAGCAGCAGCAGCAGCAGCAGCAGCAGCAGCAGCAGCAGCAGCAGCAGCAGCAGCAGCAG**CAACAGCCG**CCACCGCCGCCGCCGCCGCCGCCGCCTCCTCAGCTTCCTCAGCCGCCGCCGCAGGCACAGCCGCTGCTGCCTCAGC | GCTGATGAAGGCCTTCGAGTCCCTCAAGTCCT-------------------------------------------------------------------------------------------------------------------------CAAG**CAACAGCCG**CCACCGCCGCCGCCGCCGCCGCCGCCTCCTCAGCTTCCTCAGCCGCCGCCGCAGGCACAGCCGCTGCTGCCTCAGC | 242 | 2003 | DEL |
| 1409 | 0.0 | GCTGATGAAGGCCTTCGAGTCCCT**CAAGTCCTTCC**AGCAGCAGCAGCAGCAGCAGCAGCAGCAGCAGCAGCAGCAGCAGCAGCAGCAGCAGCAGCAGCAGCAGCAGCAGCAGCAGCAGCAGCAGCAGCAGCAGCAGCAGCAGCAGCAGCAGCAGCAGCAGCAG**CAACAGCCG**CCACCGCCGCCGCCGCCGCCGCCGCCTCCTCAGCTTCCTCAGCCGCCGCCGCAGGCACAGCCGCTGCTGCCTCAGC | GCTGATGAAGGCCTTCGAGTCCCTCAA------------------------------------------------------------------------------------------------------------------------------------------------------------GCCGCCGCCTCCTCAGCTTCCTCAGCCGCCGCCGCAGGCACAGCCGCTGCTGCCTCAGC | 242 | 1917 | DEL |
| 1317 | 0.0 | GCTGATGAAGGCCTTCGAGTCCCT**CAAGTCCTTCC**AGCAGCAGCAGCAGCAGCAGCAGCAGCAGCAGCAGCAGCAGCAGCAGCAGCAGCAGCAGCAGCAGCAGCAGCAGCAGCAGCAGCAGCAGCAGCAGCAGCAGCAGCAGCAGCAGCAGCAGCAGCAGCAG**CAACAGCCG**CCACCGCCGCCGCCGCCGCCGCCGCCTCCTCAGCTTCCTCAGCCGCCGCCGCAGGCACAGCCGCTGCTGCCTCAGC | GCTGATGAAGGCCTTCGAGTCCCTCAACT------------------------------------------------------------------------------------------------------------------------------------------------CGAGGCCGCCGCCGCCGCCTCCTCAGCTTCCTCAGCCGCCGCCGCAGGCACAGCCGCTGCTGCCTCAGC | 242 | 1912 | DEL |
| 1182 | 0.0 | GCTGATGAAGGCCTTCGAGTCCCT**CAAGTCCTTCC**AGCAGCAGCAGCAGCAGCAGCAGCAGCAGCAGCAGCAGCAGCAGCAGCAGCAGCAGCAGCAGCAGCAGCAGCAGCAGCAGCAGCAGCAGCAGCAGCAGCAGCAGCAGCAGCAGCAGCAGCAGCAGCAG**CAACAGCCG**CCACCGCCGCCGCCGCCGCCGCCGCCTCCTCAGCTTCCTCAGCCGCCGCCGCAGGCACAGCCGCTGCTGCCTCAGC | GCTGATGAAGGCCTTCGAGTCCCTCA---------------------------------------------------------------------------------------------------------------------------------AG**CAACAGCCG**CCACCGCCGCCGCCGCCGCCGCCGCCTCCTCAGCTTCCTCAGCCGCCGCCGCAGGCACAGCCGCTGCTGCCTCAGC | 242 | 1837 | DEL |
| 107 | 41.0 | GCTGATGAAGGCCTTCGAGTCCCTCAAGTCCTTC--CAGCAGCAGCAGCAGCAGCAGCAGCAGCAGCAGCAGCAGCAGCAGCAGCAGCAGCAGCAGCAGCAGCAGCAGCAGCAGCAGCAGCAGCAGCAGCAGCAGCAGCAGCAGCAGCAGCAGCAGCAGCAGCAGCAG**CAACAGCCG**CCACCGCCGCCGCCGCCGCCGCCGCCTCCTCAGCTTCCTCAGCCGCCGCCGCAGGCACAGCCGCTGCTGCCTCAGC | GCTGATGAAGGCCTTCGAGTCCCT**CAAGTCCTTCC**ACAGCAGCAGCAGCAGCAGCAGCAGCAGCAGCAGCAGCAGCAGCAGCAGCAGCAGCAGCAGCAGCAGCAGCAGCAGCAGCAGCAGCAGCAGCAGCAGCAGCAGCAGCAGCAGCAGCAGCAGCAGCAGCAG**CAACAGCCG**CCACCGCCGCCGCCGCCGCCGCCGCCTCCTCAGCTTCCTCAGCCGCCGCCGCAGGCACAGCCGCTGCTGCCTCAGC | 242 | 1756 | INS |
| 122 | 40.0 | GCTGATGAAGGCCTTCGAGTCCCT**CAAGTCCTTCC**AGCAGCAGCAGCAGCAGCAGCAGCAGCAGCAGCAGCAGCAGCAGCAGCAGCAGCAGCAGCAGCAGCAGCAGCAGCAGCAGCAGCAGCAGCAGCAGCAGCAGCAGCAGCAGCAGCAGCAGCAGCAGCAG**CAACAGCCG**CCACCGCCGCCGCCGCCGCCGCCGCCTCCTCAGCTTCCTCAGCCGCCGCCGCAGGCACAGCCGCTGCTGCCTCAGC | GCTGATGAAGGCCTTCGAGTCCCT**CAAGTCCTTCC**A-CAGCAGCAGCAGCAGCAGCAGCAGCAGCAGCAGCAGCAGCAGCAGCAGCAGCAGCAGCAGCAGCAGCAGCAGCAGCAGCAGCAGCAGCAGCAGCAGCAGCAGCAGCAGCAGCAGCAGCAGCAG**CAACAGCCG**CCACCGCCGCCGCCGCCGCCGCCGCCTCCTCAGCTTCCTCAGCCGCCGCCGCAGGCACAGCCGCTGCTGCCTCAGC | 242 | 1555 | DEL |
| 207 | 39.0 | GCTGATGAAGGCCTTCGAGTCCCT**CAAGTCCTTCC**AGCAGCAGCAGCAGCAGCAGCAGCAGCAGCAGCAGCAGCAGCAGCAGCAGCAGCAGCAGCAGCAGCAGCAGCAGCAGCAGCAGCAGCAGCAGCAGCAGCAGCAGCAGCAGCAGCAGCAGCAGCAGCAG**CAACAGCCG**CCACCGCCGCCGCCGCCGCCGCCGCCTCCTCAGCTTCCTCAGCCGCCGCCGCAGGCACAGCCGCTGCTGCCTCAGC | GCTGATGAAGGCCTTCGAGTCCCTCAAGTCCTTC------CAGCAGCAGCAGCAGCAGCAGCAGCAGCAGCAGCAGCAGCAGCAGCAGCAGCAGCAGCAGCAGCAGCAGCAGCAGCAGCAGCAGCAGCAGCAGCAGCAGCAGCAGCAGCAGCAGCAG**CAACAGCCG**CCACCGCCGCCGCCGCCGCCGCCGCCTCCTCAGCTTCCTCAGCCGCCGCCGCAGGCACAGCCGCTGCTGCCTCAGC | 242 | 1388 | DEL |
| 1345 | 0.0 | | | | | |

GCTGATGAAGGCCTTCGAGTCCCT**CAAGTCCTTCC**AGCAGCAGCAGCAGCAGCAGCAGCAGCAGCAGCAGCAGCAGCAGCAGCAGCAGCAGCAGCAGCAGCAGCAGCAGCAGCAGCAGCAGCAGCAGCAGCAGCAGCAGCAGCAGCAGCAGCAGCAGCAG**CAACAGCCG**CCACCGCCGCCGCCGCCGCCGCCGCCTCCTCAGCTTCCTCAGCCGCCGCCGCAGGCACAGCCGCTGCTGCCTCAGC
GCTGATGAAGGCCTTCGAGTCCCTCAAGTCCTTC---------------------------------------------------------------------------------------------------------------------------------------------------CCGCCGCCGCCTCCTCAGCTTCCTCAGCCGCCGCCGCAGGCACAGCCGCTGCTGCCTCAGC 242 1269 DEL
1131 0.0
GCTGATGAAGGCCTTCGAGTCCCT**CAAGTCCTTCC**AGCAGCAGCAGCAGCAGCAGCAGCAGCAGCAGCAGCAGCAGCAGCAGCAGCAGCAGCAGCAGCAGCAGCAGCAGCAGCAGCAGCAGCAGCAGCAGCAGCAGCAGCAGCAGCAGCAGCAGCAGCAG**CAACAGCCG**CCACCGCCGCCGCCGCCGCCGCCGCCTCCTCAGCTTCCTCAGCCGCCGCCGCAGGCACAGCCGCTGCTGCCTCAGC
GCTGATGAAGGCCTTCGAGTCCCTCAAG----------------------------------------------------------------------------------------------------------------------------GAGAG**CAACAGCCG**CCACCGCCGCCGCCGCCGCCGCCGCCTCCTCAGCTTCCTCAGCCGCCGCCGCAGGCACAGCCGCTGCTGCCTCAGC 242 1259 DEL
1550 0.0
GCTGATGAAGGCCTTCGAGTCCCT**CAAGTCCTTCC**AGCAGCAGCAGCAGCAGCAGCAGCAGCAGCAGCAGCAGCAGCAGCAGCAGCAGCAGCAGCAGCAGCAGCAGCAGCAGCAGCAGCAGCAGCAGCAGCAGCAGCAGCAGCAGCAGCAGCAGCAGCAG**CAACAGCCG**CCACCGCCGCCGCCGCCGCCGCCGCCTCCTCAGCTTCCTCAGCCGCCGCCGCAGGCACAGCCGCTGCTGCCTCAGC
GCTGATGAAGGCCTTCGAGTCCCTCAAGTC--------------------------------------------------------------------------------------------------------------------------------------------------------------------------------------AGCCGCAGGCACAGCCGCTGCTGCCTCAGC 242 1247 DEL
1291 0.0
GCTGATGAAGGCCTTCGAGTCCCT**CAAGTCCTTCC**AGCAGCAGCAGCAGCAGCAGCAGCAGCAGCAGCAGCAGCAGCAGCAGCAGCAGCAGCAGCAGCAGCAGCAGCAGCAGCAGCAGCAGCAGCAGCAGCAGCAGCAGCAGCAGCAGCAGCAGCAGCAG**CAACAGCCG**CCACCGCCGCCGCCGCCGCCGCCGCCTCCTCAGCTTCCTCAGCCGCCGCCGCAGGCACAGCCGCTGCTGCCTCAGC
GCTGATGAAGGCCTTCGAGTCCCT**CAAGTCCTTCC**A---------------------------------------------------------------------------------------------------------------------------------------------GCCGCCGCCGCCGCCTCCTCAGCTTCCTCAGCCGCCGCCGCAGGCACAGCCGCTGCTGCCTCAGC 242 1200 DEL
1098 0.0
GCTGATGAAGGCCTTCGAGTCCCT**CAAGTCCTTCC**AGCAGCAGCAGCAGCAGCAGCAGCAGCAGCAGCAGCAGCAGCAGCAGCAGCAGCAGCAGCAGCAGCAGCAGCAGCAGCAGCAGCAGCAGCAGCAGCAGCAGCAGCAGCAGCAGCAGCAGCAGCAG**CAACAGCCG**CCACCGCCGCCGCCGCCGCCGCCGCCTCCTCAGCTTCCTCAGCCGCCGCCGCAGGCACAGCCGCTGCTGCCTCAGC
GCTGATGAAGGCCTTCGAGTCCCTCAAGTCACCGCA-----------------------------------------------------------------------------------------------------------------------AG**CAACAGCCG**CCACCGCCGCCGCCGCCGCCGCCGCCTCCTCAGCTTCCTCAGCCGCCGCCGCAGGCACAGCCGCTGCTGCCTCAGC 242 1197 DEL
1190 0.0
GCTGATGAAGGCCTTCGAGTCCCT**CAAGTCCTTCC**AGCAGCAGCAGCAGCAGCAGCAGCAGCAGCAGCAGCAGCAGCAGCAGCAGCAGCAGCAGCAGCAGCAGCAGCAGCAGCAGCAGCAGCAGCAGCAGCAGCAGCAGCAGCAGCAGCAGCAGCAGCAG**CAACAGCCG**CCACCGCCGCCGCCGCCGCCGCCGCCTCCTCAGCTTCCTCAGCCGCCGCCGCAGGCACAGCCGCTGCTGCCTCAGC
GCTGATGAAGGCCTTCGAGTCCCTCAAGTC----------------------------------------------------------------------------------------------------------------------------------CAGCCGCCACCGCCGCCGCCGCCGCCGCCGCCTCCTCAGCTTCCTCAGCCGCCGCCGCAGGCACAGCCGCTGCTGCCTCAGC 242 1140 DEL
1589 0.0
GCTGATGAAGGCCTTCGAGTCCCT**CAAGTCCTTCC**AGCAGCAGCAGCAGCAGCAGCAGCAGCAGCAGCAGCAGCAGCAGCAGCAGCAGCAGCAGCAGCAGCAGCAGCAGCAGCAGCAGCAGCAGCAGCAGCAGCAGCAGCAGCAGCAGCAGCAGCAGCAG**CAACAGCCG**CCACCGCCGCCGCCGCCGCCGCCGCCTCCTCAGC
GCTGATGAAGGCCTTCGAGTCCCT**CAAGTCCTTCC**A---------------------------------------------------------------------------------------------------------------------------------------------------GCCGCTGCTGCCTCAGC 200 1090 DEL
1502 0.0
GCTGATGAAGGCCTTCGAGTCCCT**CAAGTCCTTCC**AGCAGCAGCAGCAGCAGCAGCAGCAGCAGCAGCAGCAGCAGCAGCAGCAGCAGCAGCAGCAGCAGCAGCAGCAGCAGCAGCAGCAGCAGCAGCAGCAGCAGCAGCAGCAGCAGCAGCAGCAGCAG**CAACAGCCG**CCACCGCCGCCGCCGCCGCCGCCGCCTCCTCAGCTTCCTCAGCCGCCGCCGCAGGCACAGCCGCTGCTGCCTCAGC
GCTGATGAAGGCCTTCGAGTCCCT**CAAGTCCTTCC**-----------------------------------------------------------------------------------------------------------------------------------------------------------------------------CCGCGGCCGCAGGCACAGCCGCTGCTGCCTCAGC 242 1085 DEL
1404 0.0
GCTGATGAAGGCCTTCGAGTCCCT**CAAGTCCTTCC**AGCAGCAGCAGCAGCAGCAGCAGCAGCAGCAGCAGCAGCAGCAGCAGCAGCAGCAGCAGCAGCAGCAGCAGCAGCAGCAGCAGCAGCAGCAGCAGCAGCAGCAGCAGCAGCAGCAGCAGCAGCAG**CAACAGCCG**CCACCGCCGCCGCCGCCGCCGCCGCCTCCTCAGCTTCCTCAGCCGCCGCCGCAGGCACAGCCGCTGCTGCCTCAGC
GCTGATGAAGGCCTTCGAGTCCCTCAAGTCCTT-----------------------------------------------------------------------------------------------------------------------------------------------------------CGCCTCCTCAGCTTCCTCAGCCGCCGCCGCAGGCACAGCCGCTGCTGCCTCAGC 242 1004 DEL
273 36.0
GCTGATGAAGGCCTTCGAGTCCCT**CAAGTCCTTCC**AGCAGCAGCAGCAGCAGCAGCAGCAGCAGCAGCAGCAGCAGCAGCAGCAGCAGCAGCAGCAGCAGCAGCAGCAGCAGCAGCAGCAGCAGCAGCAGCAGCAGCAGCAGCAGCAGCAGCAGCAGCAG**CAACAGCCG**CCACCGCCGCCGCCGCCGCCGCCGCCTCCTCAGCTTCCTCAGCCGCCGCCGCAGGCACAGCCGCTGCTGCCTCAGC
GCTGATGAAGGCCTTCGAGTCCCT**CAAGTCCTTCC**A-----------AGCAGCAGCAGCAGCAGCAGCAGCAGCAGCAGCAGCAGCAGCAGCAGCAGCAGCAGCAGCAGCAGCAGCAGCAGCAGCAGCAGCAGCAGCAGCAGCAGCAGCAGCAGCAG**CAACAGCCG**CCACCGCCGCCGCCGCCGCCGCCGCCTCCTCAGCTTCCTCAGCCGCCGCCGCAGGCACAGCCGCTGCTGCCTCAGC 242 984 DEL
155 39.0
GCTGATGAAGGCCTTCGAGTCCCT**CAAGTCCTTCC**AGCAGCAGCAGCAGCAGCAGCAGCAGCAGCAGCAGCAGCAGCAGCAGCAGCAGCAGCAGCAGCAGCAGCAGCAGCAGCAGCAGCAGCAGCAGCAGCAGCAGCAGCAGCAGCAGCAGCAGCAGCAG**CAACAGCCG**CCACCGCCGCCGCCGCCGCCGCCGCCTCCTCAGCTTCCTCAGCCGCCGCCGCAGGCACAGCCGCTGCTGCCTCAGC
GCTGATGAAGGCCTTCGAGTCCCT**CAAGTCCTTCC**A----CAGCAGCAGCAGCAGCAGCAGCAGCAGCAGCAGCAGCAGCAGCAGCAGCAGCAGCAGCAGCAGCAGCAGCAGCAGCAGCAGCAGCAGCAGCAGCAGCAGCAGCAGCAGCAGCAGCAGCAG**CAACAGCCG**CCACCGCCGCCGCCGCCGCCGCCGCCTCCTCAGCTTCCTCAGCCGCCGCCGCAGGCACAGCCGCTGCTGCCTCAGC 242 970 DEL
1171 0.0
GCTGATGAAGGCCTTCGAGTCCCT**CAAGTCCTTCC**AGCAGCAGCAGCAGCAGCAGCAGCAGCAGCAGCAGCAGCAGCAGCAGCAGCAGCAGCAGCAGCAGCAGCAGCAGCAGCAGCAGCAGCAGCAGCAGCAGCAGCAGCAGCAGCAGCAGCAGCAGCAG**CAACAGCCG**CCACCGCCGCCGCCGCCGCCGCCGCCTCCTCAGCTTCCTCAGCCGCCGCCGCAGGCACAGCCGCTGCTGCCTCAGC
GCTGATGAAGGCCTTCGAGTCCCT**CAAGTCCTTCC**A--------------------------------------------------------------------------------------------------------------------------------AGCCACCGCCGCCGCCGCCGCCGCCGCCTCCTCAGCTTCCTCAGCCGCCGCCGCAGGCACAGCCGCTGCTGCCTCAGC 242 948 DEL
1576 0.0
GCTGATGAAGGCCTTCGAGTCCCT**CAAGTCCTTCC**AGCAGCAGCAGCAGCAGCAGCAGCAGCAGCAGCAGCAGCAGCAGCAGCAGCAGCAGCAGCAGCAGCAGCAGCAGCAGCAGCAGCAGCAGCAGCAGCAGCAGCAGCAGCAGCAGCAGCAGCAGCAG**CAACAGCCG**CCACCGCCGCCGCCGCCGCCGCCGCCTCCTCAGC
GCTGATGAAGGCCTTCGAGTCCCTCAAGTCCTT-------------------------------------------------------------------------------------------------------------------------------------------------CCGCCGCCGCCGCCTCCTCAGC 200 945 DEL
74 42.0
GCTGATGAAGGCCTTCGAGTCCCTCAAGTCCTTC-------CAGCAGCAGCAGCAGCAGCAGCAGCAGCAGCAGCAGCAGCAGCAGCAGCAGCAGCAGCAGCAGCAGCAGCAGCAGCAGCAGCAGCAGCAGCAGCAGCAGCAGCAGCAGCAGCAGCAGCAGCAGCAGCAG**CAACAGCCG**CCACCGCCGCCGCCGCCGCCGCCGCCTCCTCAGCTTCCTCAGCCGCCGCCGCAGGCACAGCCGCTGCTGCCTCAGC
GCTGATGAAGGCCTTCGAGTCCCT**CAAGTCCTTCC**AAGCAGCAGCAGCAGCAGCAGCAGCAGCAGCAGCAGCAGCAGCAGCAGCAGCAGCAGCAGCAGCAGCAGCAGCAGCAGCAGCAGCAGCAGCAGCAGCAGCAGCAGCAGCAGCAGCAGCAGCAGCAGCAGCAG**CAACAGCCG**CCACCGCCGCCGCCGCCGCCGCCGCCTCCTCAGCTTCCTCAGCCGCCGCCGCAGGCACAGCCGCTGCTGCCTCAGC 242 940 INS
1318 0.0
GCTGATGAAGGCCTTCGAGTCCCT**CAAGTCCTTCC**AGCAGCAGCAGCAGCAGCAGCAGCAGCAGCAGCAGCAGCAGCAGCAGCAGCAGCAGCAGCAGCAGCAGCAGCAGCAGCAGCAGCAGCAGCAGCAGCAGCAGCAGCAGCAGCAGCAGCAGCAGCAG**CAACAGCCG**CCACCGCCGCCGCCGCCGCCGCCGCCTCCTCAGCTTCCTCAGCCGCCGCCGCAGGCACAGCCGCTGCTGCCTCAGC
GCTGATGAAGGCCTTCGAGTCCCT**CAAGTCCTTCC**A------------------------------------------------------------------------------------------------------------------------------------------------GCCGCTGCTGCCTCCTCAGCTTCCTCAGCCGCCGCCGCAGGCACAGCCGCTGCTGCCTCAGC 242 930 DEL
1067 0.0
GCTGATGAAGGCCTTCGAGTCCCT**CAAGTCCTTCC**AGCAGCAGCAGCAGCAGCAGCAGCAGCAGCAGCAGCAGCAGCAGCAGCAGCAGCAGCAGCAGCAGCAGCAGCAGCAGCAGCAGCAGCAGCAGCAGCAGCAGCAGCAGCAGCAGCAGCAGCAGCAG**CAACAGCCG**CCACCGCCGCCGCCGCCGCCGCCGCCTCCTCAGCTTCCTCAGCCGCCGCCGCAGGCACAGCCGCTGCTGCCTCAGC
GCTGATGAAGGCCTTCGAGTCCCTCAAGTCCTT------------------------------------------------------------------------------------------------------------------GAGGTGGTGG**CAACAGCCG**CCACCGCCGCCGCCGCCGCCGCCGCCTCCTCAGCTTCCTCAGCCGCCGCCGCAGGCACAGCCGCTGCTGCCTCAGC 242 881 DEL
256 39.0
GCTGATGAAGGCCTTCGAGTCCCT**CAAGTCCTTCC**AGCAGCAGCAGCAGCAGCAGCAGCAGCAGCAGCAGCAGCAGCAGCAGCAGCAGCAGCAGCAGCAGCAGCAGCAGCAGCAGCAGCAGCAGCAGCAGCAGCAGCAGCAGCAGCAGCAGCAGCAGCAG**CAACAGCCG**CCACCGCCGCCGCCGCCGCCGCCGCCTCCTCAGCTTCCTCAGCCGCCGCCGCAGGCACAGCCGCTGCTGCCTCAGC
GCTGATGAAGGCCTTCGAGTCCCTCAAGT---------AGCAGCAGCAGCAGCAGCAGCAGCAGCAGCAGCAGCAGCAGCAGCAGCAGCAGCAGCAGCAGCAGCAGCAGCAGCAGCAGCAGCAGCAGCAGCAGCAGCAGCAGCAGCAGCAGCAGCAGCAG**CAACAGCCG**CCACCGCCGCCGCCGCCGCCGCCGCCTCCTCAGCTTCCTCAGCCGCCGCCGCAGGCACAGCCGCTGCTGCCTCAGC 242 855 DEL
1161 0.0
GCTGATGAAGGCCTTCGAGTCCCT**CAAGTCCTTCC**AGCAGCAGCAGCAGCAGCAGCAGCAGCAGCAGCAGCAGCAGCAGCAGCAGCAGCAGCAGCAGCAGCAGCAGCAGCAGCAGCAGCAGCAGCAGCAGCAGCAGCAGCAGCAGCAGCAGCAGCAGCAG**CAACAGCCG**CCACCGCCGCCGCCGCCGCCGCCGCCTCCTCAGCTTCCTCAGCCGCCGCCGCAGGCACAGCCGCTGCTGCCTCAGC
GCTGATGAAGGCCTTCGAGTCCCT-------------------------------------------------------------------------------------------------------------------------------CAGCAG**CAACAGCCG**CCACCGCCGCCGCCGCCGCCGCCGCCTCCTCAGCTTCCTCAGCCGCCGCCGCAGGCACAGCCGCTGCTGCCTCAGC 242 844 DEL
1088 0.0
GCTGATGAAGGCCTTCGAGTCCCT**CAAGTCCTTCC**AGCAGCAGCAGCAGCAGCAGCAGCAGCAGCAGCAGCAGCAGCAGCAGCAGCAGCAGCAGCAGCAGCAGCAGCAGCAGCAGCAGCAGCAGCAGCAGCAGCAGCAGCAGCAGCAGCAGCAGCAGCAG**CAACAGCCG**CCACCGCCGCCGCCGCCGCCGCCGCCTCCTCAGCTTCCTCAGCCGCCGCCGCAGGCACAGCCGCTGCTGCCTCAGC
GCTGATGAAGGCCTTCGAGTCCCTCAAGTCCTTC---------------------------------------------------------------------------------------------------------------------CAGCAG**CAACAGCCG**CCACCGCCGCCGCCGCCGCCGCCGCCTCCTCAGCTTCCTCAGCCGCCGCCGCAGGCACAGCCGCTGCTGCCTCAGC 242 750 DEL
1383 0.0
GCTGATGAAGGCCTTCGAGTCCCT**CAAGTCCTTCC**AGCAGCAGCAGCAGCAGCAGCAGCAGCAGCAGCAGCAGCAGCAGCAGCAGCAGCAGCAGCAGCAGCAGCAGCAGCAGCAGCAGCAGCAGCAGCAGCAGCAGCAGCAGCAGCAGCAGCAGCAGCAG**CAACAGCCG**CCACCGCCGCCGCCGCCGCCGCCGCCTCCTCAGCTTCCTCAGCCGCCGCCGCAGGCACAGCCGCTGCTGCCTCAGC
GCTGATGAAGGCCTTCGAGTCCCTCAAGT--------------------------------------------------------------------------------------------------------------------------------------------------------CCGCCGCCGCCTCCTCAGCTTCCTCAGCCGCCGCCGCAGGCACAGCCGCTGCTGCCTCAGC 242 747 DEL
1183 0.0
GCTGATGAAGGCCTTCGAGTCCCT**CAAGTCCTTCC**AGCAGCAGCAGCAGCAGCAGCAGCAGCAGCAGCAGCAGCAGCAGCAGCAGCAGCAGCAGCAGCAGCAGCAGCAGCAGCAGCAGCAGCAGCAGCAGCAGCAGCAGCAGCAGCAGCAGCAGCAGCAG**CAACAGCCG**CCACCGCCGCCGCCGCCGCCGCCGCCTCCTCAGCTTCCTCAGCCGCCGCCGCAGGCACAGCCGCTGCTGCCTCAGC
GCTGATGAAGGCCTTCGAGTCCCT**CAAGTCCTTCC**A---------------------------------------------------------------------------------------------------------------------------------GCCACCGCCGCCGCCGCCGCCGCCGCCTCCTCAGCTTCCTCAGCCGCCGCCGCAGGCACAGCCGCTGCTGCCTCAGC 242 723 DEL
1267 0.0
GCTGATGAAGGCCTTCGAGTCCCT**CAAGTCCTTCC**AGCAGCAGCAGCAGCAGCAGCAGCAGCAGCAGCAGCAGCAGCAGCAGCAGCAGCAGCAGCAGCAGCAGCAGCAGCAGCAGCAGCAGCAGCAGCAGCAGCAGCAGCAGCAGCAGCAGCAGCAGCAG**CAACAGCCG**CCACCGCCGCCGCCGCCGCCGCCGCCTCCTCAGCTTCCTCAGCCGCCGCCGCAGGCACAGCCGCTGCTGCCTCAGC
GCTGATGAAGGCCTTCGAGTCCCTCAAGTCCTTCAA------------------------------------------------------------------------------------------------------------------------------------------GCCGCCGCCGCCGCCGCCTCCTCAGCTTCCTCAGCCGCCGCCGCAGGCACAGCCGCTGCTGCCTCAGC 242 687 DEL
1050 0.0
GCTGATGAAGGCCTTCGAGTCCCTCAAGTCCT----TCCAGCAGCAGCAGCAGCAGCAGCAGCAGCAGCAGCAGCAGCAGCAGCAGCAGCAGCAGCAGCAGCAGCAGCAGCAGCAGCAGCAGCAGCAGCAGCAGCAGCAGCAGCAGCAGCAGCAGCAGCAGCAG**CAACAGCCG**CCACCGCCGCCGCCGCCGCCGCCGCCTCCTCAGCTTCCTCAGCCGCCGCCGCAGGCACAGCCGCTGCTGCCTCAGC
GCTGATGAAGGCCTTCGAGTCCCTCAAGTCCTCAAGTCCCTCA--------------------------------------------------------------------------------------------------------------------AG**CAACAGCCG**CCACCGCCGCCGCCGCCGCCGCCGCCTCCTCAGCTTCCTCAGCCGCCGCCGCAGGCACAGCCGCTGCTGCCTCAGC 242 669 INS+DEL
1234 0.0
GCTGATGAAGGCCTTCGAGTCCCT**CAAGTCCTTCC**AGCAGCAGCAGCAGCAGCAGCAGCAGCAGCAGCAGCAGCAGCAGCAGCAGCAGCAGCAGCAGCAGCAGCAGCAGCAGCAGCAGCAGCAGCAGCAGCAGCAGCAGCAGCAGCAGCAGCAGCAGCAG**CAACAGCCG**CCACCGCCGCCGCCGCCGCCGCCGCCTCCTCAGCTTCCTCAGCCGCCGCCGCAGGCACAGCCGCTGCTGCCTCAGC
GCTGATGAAGGCCTTCGAGTCCCT**CAAGTCCTTCC**A---------------------------------------------------------------------------------------------------------------------------------------GCCGCCGCCGCCGCCGCCGCCTCCTCAGCTTCCTCAGCCGCCGCCGCAGGCACAGCCGCTGCTGCCTCAGC 242 665 DEL
1608 0.0
GCTGATGAAGGCCTTCGAGTCCCT**CAAGTCCTTCC**AGCAGCAGCAGCAGCAGCAGCAGCAGCAGCAGCAGCAGCAGCAGCAGCAGCAGCAGCAGCAGCAGCAGCAGCAGCAGCAGCAGCAGCAGCAGCAGCAGCAGCAGCAGCAGCAGCAGCAGCAGCAG**CAACAGCCG**CCACCGCCGCCGCCGCCGCCGCCGCCTCCTCAGCTTCCTCAGCCGCCGCCGCAGGCACAGCCGCTGCTGCCTCAGC
GCTGATGAAGGCCTTCGAGTC-------------------------------------------------------------------------------------------------------------------------------------------------------------------------------------------------CCGCAGGCACAGCCGCTGCTGCCTCAGC 242 647 DEL
796 15.0
GCTGATGAAGGCCTTCGAGTCCCT**CAAGTCCTTCC**AGCAGCAGCAGCAGCAGCAGCAGCAGCAGCAGCAGCAGCAGCAGCAGCAGCAGCAGCAGCAGCAGCAGCAGCAGCAGCAGCAGCAGCAGCAGCAGCAGCAGCAGCAGCAGCAGCAGCAGCAGCAG**CAACAGCCG**CCACCGCCGCCGCCGCCGCCGCCGCCTCCTCAGCTTCCTCAGCCGCCGCCGCAGGCACAGCCGCTGCTGCCTCAGC
GCTGATGAAGGCCTTCGAGTCCCT**CAAGTCCTTCC**A--------------------------------------------------------------------------AGCAGCAGCAGCAGCAGCAGCAGCAGCAGCAGCAGCAGCAGCAGCAGCAG**CAACAGCCG**CCACCGCCGCCGCCGCCGCCGCCGCCTCCTCAGCTTCCTCAGCCGCCGCCGCAGGCACAGCCGCTGCTGCCTCAGC 242 641 DEL
102 42.0
GCTGATGAAGGCCTTCGAGTCCCTCAAGTCCTTC---CAGCAGCAGCAGCAGCAGCAGCAGCAGCAGCAGCAGCAGCAGCAGCAGCAGCAGCAGCAGCAGCAGCAGCAGCAGCAGCAGCAGCAGCAGCAGCAGCAGCAGCAGCAGCAGCAGCAGCAGCAGCAG**CAACAGCCG**CCACCGCCGCCGCCGCCGCCGCCGCCTCCTCAGCTTCCTCAGCCGCCGCCGCAGGCACAGCCGCTGCTGCCTCAGC
GCTGATGAAGGCCTTCGAGTCCCT**CAAGTCCTTCC**AGCAGCAGCAGCAGCAGCAGCAGCAGCAGCAGCAGCAGCAGCAGCAGCAGCAGCAGCAGCAGCAGCAGCAGCAGCAGCAGCAGCAGCAGCAGCAGCAGCAGCAGCAGCAGCAGCAGCAGCAGCAGCAG**CAACAGCCG**CCACCGCCGCCGCCGCCGCCGCCGCCTCCTCAGCTTCCTCAGCCGCCGCCGCAGGCACAGCCGCTGCTGCCTCAGC 242 632 INS
1489 0.0
GCTGATGAAGGCCTTCGAGTCCCT**CAAGTCCTTCC**AGCAGCAGCAGCAGCAGCAGCAGCAGCAGCAGCAGCAGCAGCAGCAGCAGCAGCAGCAGCAGCAGCAGCAGCAGCAGCAGCAGCAGCAGCAGCAGCAGCAGCAGCAGCAGCAGCAGCAGCAGCAG**CAACAGCCG**CCACCGCCGCCGCCGCCGCCGCCGCCTCCTCAGCTTCCTCAGCCGCCGCCGCAGGCACAGCCGCTGCTGCCTCAGC
GCTGATGAAGGCCTTCGAGTCCCTCA---------------------------------------------------------------------------------------------------------------------------------------------------------------------------AGCTTCCTCAGCCGCCGCCGCAGGCACAGCCGCTGCTGCCTCAGC 242 630 DEL
900 13.0
GCTGATGAAGGCCTTCGAGTCCCT**CAAGTCCTTCC**AGCAGCAGCAGCAGCAGCAGCAGCAGCAGCAGCAGCAGCAGCAGCAGCAGCAGCAGCAGCAGCAGCAGCAGCAGCAGCAGCAGCAGCAGCAGCAGCAGCAGCAGCAGCAGCAGCAGCAGCAGCAG**CAACAGCCG**CCACCGCCGCCGCCGCCGCCGCCGCCTCCTCAGCTTCCTCAGCCGCCGCCGCAGGCACAGCCGCTGCTGCCTCAGC
GCTGATGAAGGCCTTCGAGTCCCTCAAGTC----------------------------------------------------------------------------------------CAGCAGCAGCAGCAGCAGCAGCAGCAGCAGCAGCAGCAG**CAACAGCCG**CCACCGCCGCCGCCGCCGCCGCCGCCTCCTCAGCTTCCTCAGCCGCCGCCGCAGGCACAGCCGCTGCTGCCTCAGC 242 624 DEL
1152 0.0
GCTGATGAAGGCCTTCGAGTCCCT**CAAGTCCTTCC**AGCAGCAGCAGCAGCAGCAGCAGCAGCAGCAGCAGCAGCAGCAGCAGCAGCAGCAGCAGCAGCAGCAGCAGCAGCAGCAGCAGCAGCAGCAGCAGCAGCAGCAGCAGCAGCAGCAGCAGCAGCAG**CAACAGCCG**CCACCGCCGCCGCCGCCGCCGCCGCCTCCTCAGCTTCCTCAGCCGCCGCCGCAGGCACAGCCGCTGCTGCCTCAGC
GCTGATGAAGGCCTTCGAGTCCCTCAAGTCCTTC------------------------------------------------------------------------------------------------------------------------------CAGCCGCCACCGCCGCCGCCGCCGCCGCCGCCTCCTCAGCTTCCTCAGCCGCCGCCGCAGGCACAGCCGCTGCTGCCTCAGC 242 622 DEL
222 38.0
GCTGATGAAGGCCTTCGAGTCCCT**CAAGTCCTTCC**AGCAGCAGCAGCAGCAGCAGCAGCAGCAGCAGCAGCAGCAGCAGCAGCAGCAGCAGCAGCAGCAGCAGCAGCAGCAGCAGCAGCAGCAGCAGCAGCAGCAGCAGCAGCAGCAGCAGCAGCAGCAG**CAACAGCCG**CCACCGCCGCCGCCGCCGCCGCCGCCTCCTCAGCTTCCTCAGCCGCCGCCGCAGGCACAGCCGCTGCTGCCTCAGC
GCTGATGAAGGCCTTCGAGTCCCT**CAAGTCCTTCC**A-------CAGCAGCAGCAGCAGCAGCAGCAGCAGCAGCAGCAGCAGCAGCAGCAGCAGCAGCAGCAGCAGCAGCAGCAGCAGCAGCAGCAGCAGCAGCAGCAGCAGCAGCAGCAGCAGCAG**CAACAGCCG**CCACCGCCGCCGCCGCCGCCGCCGCCTCCTCAGCTTCCTCAGCCGCCGCCGCAGGCACAGCCGCTGCTGCCTCAGC 242 621 DEL
1126 2.0
GCTGATGAAGGCCTTCGAGTCCCT**CAAGTCCTTCC**AGCAGCAGCAGCAGCAGCAGCAGCAGCAGCAGCAGCAGCAGCAGCAGCAGCAGCAGCAGCAGCAGCAGCAGCAGCAGCAGCAGCAGCAGCAGCAGCAGCAGCAGCAGCAGCAGCAGCAGCAGCAG**CAACAGCCG**CCACCGCCGCCGCCGCCGCCGCCGCCTCCTCAGCTTCCTCAGCCGCCGCCGCAGGCACAGCCGCTGCTGCCTCAGC
GCTGATGAAGGCCTTCGAGTCCCTCA---------------------------------------------------------------------------------------------------------------------------AGCAGCAG**CAACAGCCG**CCACCGCCGCCGCCGCCGCCGCCGCCTCCTCAGCTTCCTCAGCCGCCGCCGCAGGCACAGCCGCTGCTGCCTCAGC 242 616 DEL
1080 4.0
GCTGATGAAGGCCTTCGAGTCCCT**CAAGTCCTTCC**AGCAGCAGCAGCAGCAGCAGCAGCAGCAGCAGCAGCAGCAGCAGCAGCAGCAGCAGCAGCAGCAGCAGCAGCAGCAGCAGCAGCAGCAGCAGCAGCAGCAGCAGCAGCAGCAGCAGCAGCAGCAG**CAACAGCCG**CCACCGCCGCCGCCGCCGCCGCCGCCTCCTCAGCTTCCTCAGCCGCCGCCGCAGGCACAGCCGCTGCTGCCTCAGC
GCTGATGAAGGCCTTCGAGTCCCTCAAGT--------------------------------------------------------------------------------------------------------------------CAGCAGCAGCAG**CAACAGCCG**CCACCGCCGCCGCCGCCGCCGCCGCCTCCTCAGCTTCCTCAGCCGCCGCCGCAGGCACAGCCGCTGCTGCCTCAGC 242 610 DEL
1162 0.0
GCTGATGAAGGCCTTCGAGTCCCT**CAAGTCCTTCC**AGCAGCAGCAGCAGCAGCAGCAGCAGCAGCAGCAGCAGCAGCAGCAGCAGCAGCAGCAGCAGCAGCAGCAGCAGCAGCAGCAGCAGCAGCAGCAGCAGCAGCAGCAGCAGCAGCAGCAGCAGCAG**CAACAGCCG**CCACCGCCGCCGCCGCCGCCGCCGCCTCCTCAGCTTCCTCAGCCGCCGCCGCAGGCACAGCCGCTGCTGCCTCAGC
GCTGATGAAGGCCTTCGAGTCCCT**CAAGTCCTTCC**-------------------------------------------------------------------------------------------------------------------------------GCCGCCACCGCCGCCGCCGCCGCCGCCGCCTCCTCAGCTTCCTCAGCCGCCGCCGCAGGCACAGCCGCTGCTGCCTCAGC 242 606 DEL
439 33.0
GCTGATGAAGGCCTTCGAGTCCCT**CAAGTCCTTCC**AGCAGCAGCAGCAGCAGCAGCAGCAGCAGCAGCAGCAGCAGCAGCAGCAGCAGCAGCAGCAGCAGCAGCAGCAGCAGCAGCAGCAGCAGCAGCAGCAGCAGCAGCAGCAGCAGCAGCAGCAGCAG**CAACAGCCG**CCACCGCCGCCGCCGCCGCCGCCGCCTCCTCAGCTTCCTCAGCCGCCGCCGCAGGCACAGCCGCTGCTGCCTCAGC
GCTGATGAAGGCCTTCGAGTCCCTCAAGT-----------------------------CAGCAGCAGCAGCAGCAGCAGCAGCAGCAGCAGCAGCAGCAGCAGCAGCAGCAGCAGCAGCAGCAGCAGCAGCAGCAGCAGCAGCAGCAGCAGCAGCAGCAG**CAACAGCCG**CCACCGCCGCCGCCGCCGCCGCCGCCTCCTCAGCTTCCTCAGCCGCCGCCGCAGGCACAGCCGCTGCTGCCTCAGC 242 590 DEL
1208 0.0
GCTGATGAAGGCCTTCGAGTCCCT**CAAGTCCTTCC**AGCAGCAGCAGCAGCAGCAGCAGCAGCAGCAGCAGCAGCAGCAGCAGCAGCAGCAGCAGCAGCAGCAGCAGCAGCAGCAGCAGCAGCAGCAGCAGCAGCAGCAGCAGCAGCAGCAGCAGCAGCAG**CAACAGCCG**CCACCGCCGCCGCCGCCGCCGCCGCCTCCTCAGCTTCCTCAGCCGCCGCCGCAGGCACAGCCGCTGCTGCCTCAGC
GCTGATGAAGGCCTTCGAGTCCCT**CAAGTCCTTCC**A------------------------------------------------------------------------------------------------------------------------------------ACCGCGGCCGCCGCCGCCGCCGCCTCCTCAGCTTCCTCAGCCGCCGCCGCAGGCACAGCCGCTGCTGCCTCAGC 242 568 DEL

715 22.0
GCTGATGAAGGCCTTCGAGTCCCT**CAAGTCCTTCC**AGCAGCAGCAGCAGCAGCAGCAGCAGCAGCAGCAGCAGCAGCAGCAGCAGCAGCAGCAGCAGCAGCAGCAGCAGCAGCAGCAGCAGCAGCAGCAGCAGCAGCAGCAGCAGCAGCAGCAGCAGCAGCAG**CAACAGCCG**CCACCGCCGCCGCCGCCGCCGCCTCCTCAGCTTCCTCAGCCGCCGCCGCAGGCACAGCCGCTGCTGCCTCAGC
GCTGATGAAGGCCTTCGAGTCCCTCA---------------------------------------------------------------AGCAGCAGCAGCAGCAGCAGCAGCAGCAGCAGCAGCAGCAGCAGCAGCAGCAGCAGCAGCAGCAGCAGCAGCAG**CAACAGCCG**CCACCGCCGCCGCCGCCGCCGCCTCCTCAGCTTCCTCAGCCGCCGCCGCAGGCACAGCCGCTGCTGCCTCAGC 242 546 DEL
88 42.0
GCTGATGAAGGCCTTCGAGTCCCTCAAGTCCTTC-----CAGCAGCAGCAGCAGCAGCAGCAGCAGCAGCAGCAGCAGCAGCAGCAGCAGCAGCAGCAGCAGCAGCAGCAGCAGCAGCAGCAGCAGCAGCAGCAGCAGCAGCAGCAGCAGCAGCAGCAGCAGCAG**CAACAGCCG**CCACCGCCGCCGCCGCCGCCGCCTCCTCAGCTTCCTCAGCCGCCGCCGCAGGCACAGCCGCTGCTGCCTCAGC
GCTGATGAAGGCCTTCGAGTCCCT**CAAGTCCTTCC**ACAGCAGCAGCAGCAGCAGCAGCAGCAGCAGCAGCAGCAGCAGCAGCAGCAGCAGCAGCAGCAGCAGCAGCAGCAGCAGCAGCAGCAGCAGCAGCAGCAGCAGCAGCAGCAGCAGCAGCAGCAGCAGCAG**CAACAGCCG**CCACCGCCGCCGCCGCCGCCGCCTCCTCAGCTTCCTCAGCCGCCGCCGCAGGCACAGCCGCTGCTGCCTCAGC 242 523 INS
1433 0.0
GCTGATGAAGGCCTTCGAGTCCCT**CAAGTCCTTCC**AGCAGCAGCAGCAGCAGCAGCAGCAGCAGCAGCAGCAGCAGCAGCAGCAGCAGCAGCAGCAGCAGCAGCAGCAGCAGCAGCAGCAGCAGCAGCAGCAGCAGCAGCAGCAGCAGCAGCAGCAGCAGCAG**CAACAGCCG**CCACCGCCGCCGCCGCCGCCGCCTCCTCAGCTTCCTCAGCCGCCGCCGCAGGCACAGCCGCTGCTGCCTCAGC
GCTGATGAAGGCCTTCGAGTCC---------------------------------------------------------------------------------------------------------------------------------------------------------------CCGCCGCCGCCTCCTCAGCTTCCTCAGCCGCCGCCGCAGGCACAGCCGCTGCTGCCTCAGC 242 520 DEL
859 12.0
GCTGATGAAGGCCTTCGAGTCCCT**CAAGTCCTTCC**AGCAGCAGCAGCAGCAGCAGCAGCAGCAGCAGCAGCAGCAGCAGCAGCAGCAGCAGCAGCAGCAGCAGCAGCAGCAGCAGCAGCAGCAGCAGCAGCAGCAGCAGCAGCAGCAGCAGCAGCAGCAGCAG**CAACAGCCG**CCACCGCCGCCGCCGCCGCCGCCTCCTCAGCTTCCTCAGCCGCCGCCGCAGGCACAGCCGCTGCTGCCTCAGC
GCTGATGAAGGCCTTCGAGTCCCT**CAAGTCCTTCC**A-----------------------------------------------------------------------------------AGCAGCAGCAGCAGCAGCAGCAGCAGCAGCAGCAGCAGCAGCAG**CAACAGCCG**CCACCGCCGCCGCCGCCGCCGCCTCCTCAGCTTCCTCAGCCGCCGCCGCAGGCACAGCCGCTGCTGCCTCAGC 242 509 DEL
1292 0.0
GCTGATGAAGGCCTTCGAGTCCCT**CAAGTCCTTCC**AGCAGCAGCAGCAGCAGCAGCAGCAGCAGCAGCAGCAGCAGCAGCAGCAGCAGCAGCAGCAGCAGCAGCAGCAGCAGCAGCAGCAGCAGCAGCAGCAGCAGCAGCAGCAGCAGCAGCAGCAGCAGCAG**CAACAGCCG**CCACCGCCGCCGCCGCCGCCGCCTCCTCAGCTTCCTCAGCCGCCGCCGCAGGCACAGCCGCTGCTGCCTCAGC
GCTGATGAAGGCCTTCGAGTCCCTCAA------------------------------------------------------------------------------------------------------------------------------------------------ACCGCCGCCGCCGCCGCCGCCTCCTCAGCTTCCTCAGCCGCCGCCGCAGGCACAGCCGCTGCTGCCTCAGC 242 464 DEL
1470 0.0
GCTGATGAAGGCCTTCGAGTCCCT**CAAGTCCTTCC**AGCAGCAGCAGCAGCAGCAGCAGCAGCAGCAGCAGCAGCAGCAGCAGCAGCAGCAGCAGCAGCAGCAGCAGCAGCAGCAGCAGCAGCAGCAGCAGCAGCAGCAGCAGCAGCAGCAGCAGCAGCAGCAG**CAACAGCCG**CCACCGCCGCCGCCGCCGCCGCCTCCTCAGCTTCCTCAGCCGCCGCCGCAGGCACAGCCGCTGCTGCCTCAGC
GCTGATGAAGGCCTTCGAGTCC----------------------------------------------------------------------------------------------------------------------------------------------------------------------CCTCCTCAGCTTCCTCAGCCGCCGCCGCAGGCACAGCCGCTGCTGCCTCAGC 242 458 DEL
1244 0.0
GCTGATGAAGGCCTTCGAGTCCCT**CAAGTCCTTCC**AGCAGCAGCAGCAGCAGCAGCAGCAGCAGCAGCAGCAGCAGCAGCAGCAGCAGCAGCAGCAGCAGCAGCAGCAGCAGCAGCAGCAGCAGCAGCAGCAGCAGCAGCAGCAGCAGCAGCAGCAGCAGCAG**CAACAGCCG**CCACCGCCGCCGCCGCCGCCGCCTCCTCAGCTTCCTCAGCCGCCGCCGCAGGCACAGCCGCTGCTGCCTCAGC
GCTGATGAAGGCCTTCGAGTCCC-------------------------------------------------------------------------------------------------------------------------------------------ACAGCCGCCACCGCCGCCGCCGCCGCCGCCTCCTCAGCTTCCTCAGCCGCCGCCGCAGGCACAGCCGCTGCTGCCTCAGC 242 456 DEL
308 35.0
GCTGATGAAGGCCTTCGAGTCCCT**CAAGTCCTTCC**AGCAGCAGCAGCAGCAGCAGCAGCAGCAGCAGCAGCAGCAGCAGCAGCAGCAGCAGCAGCAGCAGCAGCAGCAGCAGCAGCAGCAGCAGCAGCAGCAGCAGCAGCAGCAGCAGCAGCAGCAGCAGCAG**CAACAGCCG**CCACCGCCGCCGCCGCCGCCGCCTCCTCAGCTTCCTCAGCCGCCGCCGCAGGCACAGCCGCTGCTGCCTCAGC
GCTGATGAAGGCCTTCGAGTCCCT**CAAGTCCTTCC**A--------------AGCAGCAGCAGCAGCAGCAGCAGCAGCAGCAGCAGCAGCAGCAGCAGCAGCAGCAGCAGCAGCAGCAGCAGCAGCAGCAGCAGCAGCAGCAGCAGCAGCAGCAGCAGCAGCAG**CAACAGCCG**CCACCGCCGCCGCCGCCGCCGCCTCCTCAGCTTCCTCAGCCGCCGCCGCAGGCACAGCCGCTGCTGCCTCAGC 242 451 DEL
266 41.0
GCTGATGAAGGCCTTCGAGTCCCT**CAAGTCCTTCC**AGCAGCAGCAGCAGCAGCAGCAGCAGCAGCAGCAGCAGCAGCAGCAGCAGCAGCAGCAGCAGCAGCAGCAGCAGCAGCAGCAGCAGCAGCAGCAGCAGCAGCAGCAGCAGCAGCAGCAGCAGCAGCAG**CAACAGCCG**CCACCGCCGCCGCCGCCGCCGCCTCCTCAGCTTCCTCAGCCGCCGCCGCAGGCACAGCCGCTGCTGCCTCAGC
GCTGATGAAGGCCTTCGAGTCCCT----------CAGCAGCAGCAGCAGCAGCAGCAGCAGCAGCAGCAGCAGCAGCAGCAGCAGCAGCAGCAGCAGCAGCAGCAGCAGCAGCAGCAGCAGCAGCAGCAGCAGCAGCAGCAGCAGCAGCAGCAGCAGCAGCAG**CAACAGCCG**CCACCGCCGCCGCCGCCGCCGCCTCCTCAGCTTCCTCAGCCGCCGCCGCAGGCACAGCCGCTGCTGCCTCAGC 242 442 DEL
771 16.0
GCTGATGAAGGCCTTCGAGTCCCT**CAAGTCCTTCC**AGCAGCAGCAGCAGCAGCAGCAGCAGCAGCAGCAGCAGCAGCAGCAGCAGCAGCAGCAGCAGCAGCAGCAGCAGCAGCAGCAGCAGCAGCAGCAGCAGCAGCAGCAGCAGCAGCAGCAGCAGCAGCAG**CAACAGCCG**CCACCGCCGCCGCCGCCGCCGCCTCCTCAGCTTCCTCAGCCGCCGCCGCAGGCACAGCCGCTGCTGCCTCAGC
GCTGATGAAGGCCTTCGAGTCCCT**CAAGTCCTTCC**A-----------------------------------------------------------------------AGCAGCAGCAGCAGCAGCAGCAGCAGCAGCAGCAGCAGCAGCAGCAGCAGCAGCAG**CAACAGCCG**CCACCGCCGCCGCCGCCGCCGCCTCCTCAGCTTCCTCAGCCGCCGCCGCAGGCACAGCCGCTGCTGCCTCAGC 242 428 DEL
301 36.0
GCTGATGAAGGCCTTCGAGTCCCT**CAAGTCCTTCC**AGCAGCAGCAGCAGCAGCAGCAGCAGCAGCAGCAGCAGCAGCAGCAGCAGCAGCAGCAGCAGCAGCAGCAGCAGCAGCAGCAGCAGCAGCAGCAGCAGCAGCAGCAGCAGCAGCAGCAGCAGCAGCAG**CAACAGCCG**CCACCGCCGCCGCCGCCGCCGCCTCCTCAGCTTCCTCAGCCGCCGCCGCAGGCACAGCCGCTGCTGCCTCAGC
GCTGATGAAGGCCTTCGAGTCCCT**CAAGTCCTTCC**A-------------CAGCAGCAGCAGCAGCAGCAGCAGCAGCAGCAGCAGCAGCAGCAGCAGCAGCAGCAGCAGCAGCAGCAGCAGCAGCAGCAGCAGCAGCAGCAGCAGCAGCAGCAGCAGCAGCAG**CAACAGCCG**CCACCGCCGCCGCCGCCGCCGCCTCCTCAGCTTCCTCAGCCGCCGCCGCAGGCACAGCCGCTGCTGCCTCAGC 242 422 DEL
414 31.0
GCTGATGAAGGCCTTCGAGTCCCT**CAAGTCCTTCC**AGCAGCAGCAGCAGCAGCAGCAGCAGCAGCAGCAGCAGCAGCAGCAGCAGCAGCAGCAGCAGCAGCAGCAGCAGCAGCAGCAGCAGCAGCAGCAGCAGCAGCAGCAGCAGCAGCAGCAGCAGCAGCAG**CAACAGCCG**CCACCGCCGCCGCCGCCGCCGCCTCCTCAGCTTCCTCAGCCGCCGCCGCAGGCACAGCCGCTGCTGCCTCAGC
GCTGATGAAGGCCTTCGAGTCCCT**CAAGTCCTTCC**A--------------------------AGCAGCAGCAGCAGCAGCAGCAGCAGCAGCAGCAGCAGCAGCAGCAGCAGCAGCAGCAGCAGCAGCAGCAGCAGCAGCAGCAGCAGCAGCAGCAGCAG**CAACAGCCG**CCACCGCCGCCGCCGCCGCCGCCTCCTCAGCTTCCTCAGCCGCCGCCGCAGGCACAGCCGCTGCTGCCTCAGC 242 421 DEL
1115 0.0
GCTGATGAAGGCCTTCGAGTCCCT**CAAGTCCTTCC**AGCAGCAGCAGCAGCAGCAGCAGCAGCAGCAGCAGCAGCAGCAGCAGCAGCAGCAGCAGCAGCAGCAGCAGCAGCAGCAGCAGCAGCAGCAGCAGCAGCAGCAGCAGCAGCAGCAGCAGCAGCAGCAG**CAACAGCCG**CCACCGCCGCCGCCGCCGCCGCCTCCTCAGCTTCCTCAGCCGCCGCCGCAGGCACAGCCGCTGCTGCCTCAGC
GCTGATGAAGGCCTTCGAGTCCCTCAAGT-----------------------------------------------------------------------------------------------------------------------------CACCACCTCAAGCCGCCACCGCCGCCGCCGCCGCCGCCTCCTCAGCTTCCTCAGCCGCCGCCGCAGGCACAGCCGCTGCTGCCTCAGC 242 418 DEL
337 34.0
GCTGATGAAGGCCTTCGAGTCCCT**CAAGTCCTTCC**AGCAGCAGCAGCAGCAGCAGCAGCAGCAGCAGCAGCAGCAGCAGCAGCAGCAGCAGCAGCAGCAGCAGCAGCAGCAGCAGCAGCAGCAGCAGCAGCAGCAGCAGCAGCAGCAGCAGCAGCAGCAGCAG**CAACAGCCG**CCACCGCCGCCGCCGCCGCCGCCTCCTCAGCTTCCTCAGCCGCCGCCGCAGGCACAGCCGCTGCTGCCTCAGC
GCTGATGAAGGCCTTCGAGTCCCT**CAAGTCCTTCC**A-----------------AGCAGCAGCAGCAGCAGCAGCAGCAGCAGCAGCAGCAGCAGCAGCAGCAGCAGCAGCAGCAGCAGCAGCAGCAGCAGCAGCAGCAGCAGCAGCAGCAGCAGCAGCAG**CAACAGCCG**CCACCGCCGCCGCCGCCGCCGCCTCCTCAGCTTCCTCAGCCGCCGCCGCAGGCACAGCCGCTGCTGCCTCAGC 242 412 DEL
464 29.0
GCTGATGAAGGCCTTCGAGTCCCT**CAAGTCCTTCC**AGCAGCAGCAGCAGCAGCAGCAGCAGCAGCAGCAGCAGCAGCAGCAGCAGCAGCAGCAGCAGCAGCAGCAGCAGCAGCAGCAGCAGCAGCAGCAGCAGCAGCAGCAGCAGCAGCAGCAGCAGCAGCAG**CAACAGCCG**CCACCGCCGCCGCCGCCGCCGCCTCCTCAGCTTCCTCAGCCGCCGCCGCAGGCACAGCCGCTGCTGCCTCAGC
GCTGATGAAGGCCTTCGAGTCCCT**CAAGTCCTTCC**A--------------------------------AGCAGCAGCAGCAGCAGCAGCAGCAGCAGCAGCAGCAGCAGCAGCAGCAGCAGCAGCAGCAGCAGCAGCAGCAGCAGCAGCAGCAGCAG**CAACAGCCG**CCACCGCCGCCGCCGCCGCCGCCTCCTCAGCTTCCTCAGCCGCCGCCGCAGGCACAGCCGCTGCTGCCTCAGC 242 404 DEL
1221 0.0
GCTGATGAAGGCCTTCGAGTCCCT**CAAGTCCTTCC**AGCAGCAGCAGCAGCAGCAGCAGCAGCAGCAGCAGCAGCAGCAGCAGCAGCAGCAGCAGCAGCAGCAGCAGCAGCAGCAGCAGCAGCAGCAGCAGCAGCAGCAGCAGCAGCAGCAGCAGCAGCAGCAG**CAACAGCCG**CCACCGCCGCCGCCGCCGCCGCCTCCTCAGCTTCCTCAGCCGCCGCCGCAGGCACAGCCGCTGCTGCCTCAGC
GCTGATGAAGGCCTTCGAGT-----------------------------------------------------------------------------------------------------------------------------------------CAG**CAACAGCCG**CCACCGCCGCCGCCGCCGCCGCCTCCTCAGCTTCCTCAGCCGCCGCCGCAGGCACAGCCGCTGCTGCCTCAGC 242 393 DEL
1346 0.0
GCTGATGAAGGCCTTCGAGTCCCT**CAAGTCCTTCC**AGCAGCAGCAGCAGCAGCAGCAGCAGCAGCAGCAGCAGCAGCAGCAGCAGCAGCAGCAGCAGCAGCAGCAGCAGCAGCAGCAGCAGCAGCAGCAGCAGCAGCAGCAGCAGCAGCAGCAGCAGCAGCAG**CAACAGCCG**CCACCGCCGCCGCCGCCGCCGCCTCCTCAGCTTCCTCAGCCGCCGCCGCAGGCACAGCCGCTGCTGCCTCAGC
GCTGATGAAGGCCTTCGAGTCCCTCAA------------------------------------------------------------------------------------------------------------------------------------------------------GCCGCCGCCGCCGCCTCCTCAGCTTCCTCAGCCGCCGCCGCAGGCACAGCCGCTGCTGCCTCAGC 242 370 DEL
536 26.0
GCTGATGAAGGCCTTCGAGTCCCT**CAAGTCCTTCC**AGCAGCAGCAGCAGCAGCAGCAGCAGCAGCAGCAGCAGCAGCAGCAGCAGCAGCAGCAGCAGCAGCAGCAGCAGCAGCAGCAGCAGCAGCAGCAGCAGCAGCAGCAGCAGCAGCAGCAGCAGCAGCAG**CAACAGCCG**CCACCGCCGCCGCCGCCGCCGCCTCCTCAGCTTCCTCAGCCGCCGCCGCAGGCACAGCCGCTGCTGCCTCAGC
GCTGATGAAGGCCTTCGAGTCCCT**CAAGTCCTTCC**A-----------------------------------------AGCAGCAGCAGCAGCAGCAGCAGCAGCAGCAGCAGCAGCAGCAGCAGCAGCAGCAGCAGCAGCAGCAGCAGCAGCAGCAGCAG**CAACAGCCG**CCACCGCCGCCGCCGCCGCCGCCTCCTCAGCTTCCTCAGCCGCCGCCGCAGGCACAGCCGCTGCTGCCTCAGC 242 368 DEL
362 33.0
GCTGATGAAGGCCTTCGAGTCCCT**CAAGTCCTTCC**AGCAGCAGCAGCAGCAGCAGCAGCAGCAGCAGCAGCAGCAGCAGCAGCAGCAGCAGCAGCAGCAGCAGCAGCAGCAGCAGCAGCAGCAGCAGCAGCAGCAGCAGCAGCAGCAGCAGCAGCAGCAGCAG**CAACAGCCG**CCACCGCCGCCGCCGCCGCCGCCTCCTCAGCTTCCTCAGCCGCCGCCGCAGGCACAGCCGCTGCTGCCTCAGC
GCTGATGAAGGCCTTCGAGTCCCT**CAAGTCCTTCC**A--------------------AGCAGCAGCAGCAGCAGCAGCAGCAGCAGCAGCAGCAGCAGCAGCAGCAGCAGCAGCAGCAGCAGCAGCAGCAGCAGCAGCAGCAGCAGCAGCAGCAGCAGCAG**CAACAGCCG**CCACCGCCGCCGCCGCCGCCGCCTCCTCAGCTTCCTCAGCCGCCGCCGCAGGCACAGCCGCTGCTGCCTCAGC 242 366 DEL
286 29.0
GCTGATGAAGGCCTTCGAGTCCCT**CAAGTCCTTCC**AGCAGCAGCAGCAGCAGCAGCAGCAGCAGCAGCAGCAGCAGCAGCAGCAGCAGCAGCAGCAGCAGCAGCAGCAGCAGCAGCAGCAGCAGCAGCAGCAGCAGCAGCAGCAGCAGCAGCAGCAGCAGCAG**CAACAGCCG**CCACCGCCGCCGCCGCCGCCGCCTCCTCAGCTTCCTCAGCCGCCGCCGCAGGCACAGCCGCTGCTGCCTCAGC
GCTGATGAAGGCCTTCGAGTC------------CCAGCAGCAGCAGCAGCAGCAGCAGCAGTAGCAGCAGCAGCAGCAGCAGCAGCAGCAGCAGCAGCAGCAGCAGCAGCAGCAGCAGCAGCAGCAGCAGCAGCAGCAGCAGCAGCAGCAG**CAACAGCCG**CCACCGCCGCCGCCGCCGCCGCCTCCTCAGCTTCCTCAGCCGCCGCCGCAGGCACAGCCGCTGCTGCCTCAGC 242 362 DEL
1257 0.0
GCTGATGAAGGCCTTCGAGTCCCT**CAAGTCCTTCC**AGCAGCAGCAGCAGCAGCAGCAGCAGCAGCAGCAGCAGCAGCAGCAGCAGCAGCAGCAGCAGCAGCAGCAGCAGCAGCAGCAGCAGCAGCAGCAGCAGCAGCAGCAGCAGCAGCAGCAGCAGCAGCAG**CAACAGCCG**CCACCGCCGCCGCCGCCGCCGCCTCCTCAGCTTCCTCAGCCGCCGCCGCAGGCACAGCCGCTGCTGCCTCAGC
GCTGATGAAGGCCTTCGAGTCCCTCAAGTCCTTCAGTC--------------------------------------------------------------------------------------------------------------------------------------------CCGCCGCCGCCGCCTCCTCAGCTTCCTCAGCCGCCGCCGCAGGCACAGCCGCTGCTGCCTCAGC 242 357 DEL
956 7.0
GCTGATGAAGGCCTTCGAGTCCCT**CAAGTCCTTCC**AGCAGCAGCAGCAGCAGCAGCAGCAGCAGCAGCAGCAGCAGCAGCAGCAGCAGCAGCAGCAGCAGCAGCAGCAGCAGCAGCAGCAGCAGCAGCAGCAGCAGCAGCAGCAGCAGCAGCAGCAGCAGCAG**CAACAGCCG**CCACCGCCGCCGCCGCCGCCGCCTCCTCAGCTTCCTCAGCCGCCGCCGCAGGCACAGCCGCTGCTGCCTCAGC
GCTGATGAAGGCCTTCGAGTCCCT**CAAGTCCTTCC**A--------------------------------------------------------------------------------------------------AGCAGCAGCAGCAGCAGCAGCAGCAG**CAACAGCCG**CCACCGCCGCCGCCGCCGCCGCCTCCTCAGCTTCCTCAGCCGCCGCCGCAGGCACAGCCGCTGCTGCCTCAGC 242 354 DEL
918 9.0
GCTGATGAAGGCCTTCGAGTCCCT**CAAGTCCTTCC**AGCAGCAGCAGCAGCAGCAGCAGCAGCAGCAGCAGCAGCAGCAGCAGCAGCAGCAGCAGCAGCAGCAGCAGCAGCAGCAGCAGCAGCAGCAGCAGCAGCAGCAGCAGCAGCAGCAGCAGCAGCAGCAG**CAACAGCCG**CCACCGCCGCCGCCGCCGCCGCCTCCTCAGCTTCCTCAGCCGCCGCCGCAGGCACAGCCGCTGCTGCCTCAGC
GCTGATGAAGGCCTTCGAGTCCCT**CAAGTCCTTCC**A--------------------------------------------------------------------------------------------AGCAGCAGCAGCAGCAGCAGCAGCAGCAGCAG**CAACAGCCG**CCACCGCCGCCGCCGCCGCCGCCTCCTCAGCTTCCTCAGCCGCCGCCGCAGGCACAGCCGCTGCTGCCTCAGC 242 354 DEL
658 21.0
GCTGATGAAGGCCTTCGAGTCCCT**CAAGTCCTTCC**AGCAGCAGCAGCAGCAGCAGCAGCAGCAGCAGCAGCAGCAGCAGCAGCAGCAGCAGCAGCAGCAGCAGCAGCAGCAGCAGCAGCAGCAGCAGCAGCAGCAGCAGCAGCAGCAGCAGCAGCAGCAGCAG**CAACAGCCG**CCACCGCCGCCGCCGCCGCCGCCTCCTCAGCTTCCTCAGCCGCCGCCGCAGGCACAGCCGCTGCTGCCTCAGC
GCTGATGAAGGCCTTCGAGTCCCT**CAAGTCCTTCC**A--------------------------------------------------------AGCAGCAGCAGCAGCAGCAGCAGCAGCAGCAGCAGCAGCAGCAGCAGCAGCAGCAGCAGCAGCAGCAGCAG**CAACAGCCG**CCACCGCCGCCGCCGCCGCCGCCTCCTCAGCTTCCTCAGCCGCCGCCGCAGGCACAGCCGCTGCTGCCTCAGC 242 351 DEL
465 40.0
GCTGATGAAGGCCTTCGAGTCCCT**CAAGTCCTTCC**-AGCAGCAGCAGCAGCAGCAGCAGCAGCAGCAGCAGCAGCAGCAGCAGCAGCAGCAGCAGCAGCAGCAGCAGCAGCAGCAGCAGCAGCAGCAGCAGCAGCAGCAGCAGCAGCAGCAGCAGCAGCAGCAG**CAACAGCCG**CCACCGCCGCCGCCGCCGCCGCCTCCTCAGCTTCCTCAGC
GCTGATGAAGGCCTTCGAGTCCCT**CAAGTCCTTCC**AAGCAGCAGCAGCAGCAGCAGCAGCAGCAGCAGCAGCAGCAGCAGCAGCAGCAGCAGCAGCAGCAGCAGCAGCAGCAGCAGCAGCAGCAGCAGCAGCAGCAGCAGCAGCAGCAGCAGCAGCAGCAGCAG**CAACAGCCG**CCACCGCCGCCGCCGCCGCCGCCTCCTCAGCAGCCTCAGC 209 344 INS
787 12.0
GCTGATGAAGGCCTTCGAGTCCCT**CAAGTCCTTCC**AGCAGCAGCAGCAGCAGCAGCAGCAGCAGCAGCAGCAGCAGCAGCAGCAGCAGCAGCAGCAGCAGCAGCAGCAGCAGCAGCAGCAGCAGCAGCAGCAGCAGCAGCAGCAGCAGCAGCAGCAGCAGCAG**CAACAGCCG**CCACCGCCGCCGCCGCCGCCGCCTCCTCAGCTTCCTCAGCCGCCGCCGCAGGCACAGCCGCTGCTGCCTCAGC
GCTGATGAAGGCCTTCGAGTCCCT**CAAGTCCTTCC**A-------------------------------------------------------------------------CAGCAGCTGCTGCAGCAGCAGCAGCAGCAGCAGCAGCAGCAGCAGCAGCAG**CAACAGCCG**CCACCGCCGCCGCCGCCGCCGCCTCCTCAGCTTCCTCAGCCGCCGCCGCAGGCACAGCCGCTGCTGCCTCAGC 242 343 DEL
349 33.0
GCTGATGAAGGCCTTCGAGTCCCT**CAAGTCCTTCC**AGCAGCAGCAGCAGCAGCAGCAGCAGCAGCAGCAGCAGCAGCAGCAGCAGCAGCAGCAGCAGCAGCAGCAGCAGCAGCAGCAGCAGCAGCAGCAGCAGCAGCAGCAGCAGCAGCAGCAGCAGCAGCAG**CAACAGCCG**CCACCGCCGCCGCCGCCGCCGCCTCCTCAGCTTCCTCAGCCGCCGCCGCAGGCACAGCCGCTGCTGCCTCAGC
GCTGATGAAGGCCTTCGAGTCCCT**CAAGTCCTTCC**AGC------------------TGCAGCAGCAGCAGCAGCAGCAGCAGCAGCAGCAGCAGCAGCAGCAGCAGCAGCAGCAGCAGCAGCAGCAGCAGCAGCAGCAGCAGCAGCAGCAGCAGCAGCAGCAG**CAACAGCCG**CCACCGCCGCCGCCGCCGCCGCCTCCTCAGCTTCCTCAGCCGCCGCCGCAGGCACAGCCGCTGCTGCCTCAGC 242 343 DEL
1308 0.0
GCTGATGAAGGCCTTCGAGTCCCT**CAAGTCCTTCC**AGCAGCAGCAGCAGCAGCAGCAGCAGCAGCAGCAGCAGCAGCAGCAGCAGCAGCAGCAGCAGCAGCAGCAGCAGCAGCAGCAGCAGCAGCAGCAGCAGCAGCAGCAGCAGCAGCAGCAGCAGCAGCAG**CAACAGCCG**CCACCGCCGCCGCCGCCGCCGCCTCCTCAGCTTCCTCAGCCGCCGCCGCAGGCACAGCCGCTGCTGCCTCAGC
GCTGATGAAGGCCTTCGAGTCCCT--------------------------------------------------------------------------------------------------------------------------------------------------CACCGCCGCCGCCGCCGCCGCCTCCTCAGCTTCCTCAGCCGCCGCCGCAGGCACAGCCGCTGCTGCCTCAGC 242 337 DEL
938 8.0
GCTGATGAAGGCCTTCGAGTCCCT**CAAGTCCTTCC**AGCAGCAGCAGCAGCAGCAGCAGCAGCAGCAGCAGCAGCAGCAGCAGCAGCAGCAGCAGCAGCAGCAGCAGCAGCAGCAGCAGCAGCAGCAGCAGCAGCAGCAGCAGCAGCAGCAGCAGCAGCAGCAG**CAACAGCCG**CCACCGCCGCCGCCGCCGCCGCCTCCTCAGCTTCCTCAGCCGCCGCCGCAGGCACAGCCGCTGCTGCCTCAGC
GCTGATGAAGGCCTTCGAGTCCCT**CAAGTCCTTCC**A-----------------------------------------------------------------------------------------------AGCAGCAGCAGCAGCAGCAGCAGCAGCAG**CAACAGCCG**CCACCGCCGCCGCCGCCGCCGCCTCCTCAGCTTCCTCAGCCGCCGCCGCAGGCACAGCCGCTGCTGCCTCAGC 242 313 DEL
490 28.0
GCTGATGAAGGCCTTCGAGTCCCT**CAAGTCCTTCC**AGCAGCAGCAGCAGCAGCAGCAGCAGCAGCAGCAGCAGCAGCAGCAGCAGCAGCAGCAGCAGCAGCAGCAGCAGCAGCAGCAGCAGCAGCAGCAGCAGCAGCAGCAGCAGCAGCAGCAGCAGCAGCAG**CAACAGCCG**CCACCGCCGCCGCCGCCGCCGCCTCCTCAGCTTCCTCAGCCGCCGCCGCAGGCACAGCCGCTGCTGCCTCAGC
GCTGATGAAGGCCTTCGAGTCCCT**CAAGTCCTTCC**A-----------------------------------AGCAGCAGCAGCAGCAGCAGCAGCAGCAGCAGCAGCAGCAGCAGCAGCAGCAGCAGCAGCAGCAGCAGCAGCAGCAGCAGCAGCAGCAGCAG**CAACAGCCG**CCACCGCCGCCGCCGCCGCCGCCTCCTCAGCTTCCTCAGCCGCCGCCGCAGGCACAGCCGCTGCTGCCTCAGC 242 311 DEL
391 32.0
GCTGATGAAGGCCTTCGAGTCCCT**CAAGTCCTTCC**AGCAGCAGCAGCAGCAGCAGCAGCAGCAGCAGCAGCAGCAGCAGCAGCAGCAGCAGCAGCAGCAGCAGCAGCAGCAGCAGCAGCAGCAGCAGCAGCAGCAGCAGCAGCAGCAGCAGCAGCAGCAGCAG**CAACAGCCG**CCACCGCCGCCGCCGCCGCCGCCTCCTCAGCTTCCTCAGCCGCCGCCGCAGGCACAGCCGCTGCTGCCTCAGC
GCTGATGAAGGCCTTCGAGTCCCT**CAAGTCCTTCC**A-----------------------AGCAGCAGCAGCAGCAGCAGCAGCAGCAGCAGCAGCAGCAGCAGCAGCAGCAGCAGCAGCAGCAGCAGCAGCAGCAGCAGCAGCAGCAGCAGCAGCAGCAG**CAACAGCCG**CCACCGCCGCCGCCGCCGCCGCCTCCTCAGCTTCCTCAGCCGCCGCCGCAGGCACAGCCGCTGCTGCCTCAGC 242 311 DEL
1127 0.0
GCTGATGAAGGCCTTCGAGTCCCT**CAAGTCCTTCC**AGCAGCAGCAGCAGCAGCAGCAGCAGCAGCAGCAGCAGCAGCAGCAGCAGCAGCAGCAGCAGCAGCAGCAGCAGCAGCAGCAGCAGCAGCAGCAGCAGCAGCAGCAGCAGCAGCAGCAGCAGCAGCAG**CAACAGCCG**CCACCGCCGCCGCCGCCGCCGCCTCCTCAGCTTCCTCAGCCGCCGCCGCAGGCACAGCCGCTGCTGCCTCAGC
GCTGATGAAGGCCTTCGAGTCCCTCAAGTCCTTC------------------------------------------------------------------------------------------------------------------------------**CAACAGCCG**CCACCGCCGCCGCCGCCGCCGCCTCCTCAGCTTCCTCAGCCGCCGCCGCAGGCACAGCCGCTGCTGCCTCAGC 242 309 DEL
537 40.0
GCTGATGAAGGCCTTCGAGTCCCT**CAAGTCCTTCC**-AGCAGCAGCAGCAGCAGCAGCAGCAGCAGCAGCAGCAGCAGCAGCAGCAGCAGCAGCAGCAGCAGCAGCAGCAGCAGCAGCAGCAGCAGCAGCAGCAGCAGCAGCAGCAGCAGCAGCAGCAGCAGCAG**CAACAGCCG**CCACCGCCGCCGCCGCCGCCGCCTCCTCAGC
GCTGATGAAGGCCTTCGAGTCCCT**CAAGTCCTTCC**AAGCAGCAGCAGCAGCAGCAGCAGCAGCAGCAGCAGCAGCAGCAGCAGCAGCAGCAGCAGCAGCAGCAGCAGCAGCAGCAGCAGCAGCAGCAGCAGCAGCAGCAGCAGCAGCAGCAGCAGCAGCAGCAG**CAACAGCCG**CCACCGCCGCCGCCGCCGCCGCCGCCTCAGC 200 304 INS
1493 0.0
GCTGATGAAGGCCTTCGAGTCCCT**CAAGTCCTTCC**AGCAGCAGCAGCAGCAGCAGCAGCAGCAGCAGCAGCAGCAGCAGCAGCAGCAGCAGCAGCAGCAGCAGCAGCAGCAGCAGCAGCAGCAGCAGCAGCAGCAGCAGCAGCAGCAGCAGCAGCAGCAGCAG**CAACAGCCG**CCACCGCCGCCGCCGCCGCCGCCTCCTCAGCTTCCTCAGCCGCCGCCGCAGGCACAGCCGCTGCTGCCTCAGC
GCTGATGAAGGCCTTCGAGTCCCT**CAAGTCCTTCC**--------------------------------------------------------------------------------------------------------------------------------------------------------------------------------GCCGCCGCCGCAGGCACAGCCGCTGCTGCCTCAGC 242 303 DEL
66 43.0
GCTGATGAAGGCCTTCGAGTCCCTCAAGTCCTTC----------CAGCAGCAGCAGCAGCAGCAGCAGCAGCAGCAGCAGCAGCAGCAGCAGCAGCAGCAGCAGCAGCAGCAGCAGCAGCAGCAGCAGCAGCAGCAGCAGCAGCAGCAGCAGCAGCAGCAGCAGCAG**CAACAGCCG**CCACCGCCGCCGCCGCCGCCGCCTCCTCAGCTTCCTCAGCCGCCGCCGCAGGCACAGCCGCTGCTGCCTCAGC
GCTGATGAAGGCCTTCGAGTCCCT**CAAGTCCTTCC**AAGCAGCAGCAGCAGCAGCAGCAGCAGCAGCAGCAGCAGCAGCAGCAGCAGCAGCAGCAGCAGCAGCAGCAGCAGCAGCAGCAGCAGCAGCAGCAGCAGCAGCAGCAGCAGCAGCAGCAGCAGCAGCAGCAGCAGCAGCAG**CAACAGCCG**CCACCGCCGCCGCCGCCGCCGCCTCCTCAGCTTCCTCAGCCGCCGCCGCAGGCACAGCCGCTGCTGCCTCAGC 242 294 INS

1434 0.0
GCTGATGAAGGCCTTCGAGTCCCT**CAAGTCCTTCC**AGCAGCAGCAGCAGCAGCAGCAGCAGCAGCAGCAGCAGCAGCAGCAGCAGCAGCAGCAGCAGCAGCAGCAGCAGCAGCAGCAGCAGCAGCAGCAGCAGCAGCAGCAGCAGCAGCAGCAGCAGCAGCAG**CAACAGCCG**CCACCGCCGCCGCCGCCGCCGCCGCCTCCTCAGCTTCCTCAGCCGCCGCCGCAGGCACAGCCGCTGCTGCCTCAGC
GCTGATGAAGGCCTTCGAGTCCCTCAAGT---------------------------------------------------------------------------------------------------------------------------------------------------------------CGCCTCCTCAGCTTCCTCAGCCGCCGCCGCAGGCACAGCCGCTGCTGCCTCAGC 242 291 DEL
1427 0.0
GCTGATGAAGGCCTTCGAGTCCCT**CAAGTCCTTCC**AGCAGCAGCAGCAGCAGCAGCAGCAGCAGCAGCAGCAGCAGCAGCAGCAGCAGCAGCAGCAGCAGCAGCAGCAGCAGCAGCAGCAGCAGCAGCAGCAGCAGCAGCAGCAGCAGCAGCAGCAGCAGCAG**CAACAGCCG**CCACCGCCGCCGCCGCCGCCGCCGCCTCCTCAGCTTCCTCAGCCGCCGCCGCAGGCACAGCCGCTGCTGCCTCAGC
GCTGATGAAGGCCTTCGAGTCCCTCAAGT--------------------------------------------------------------------------------------------------------------------------------------------------------------CCGCCTCCTCAGCTTCCTCAGCCGCCGCCGCAGGCACAGCCGCTGCTGCCTCAGC 242 289 DEL
1163 5.0
GCTGATGAAGGCCTTCGAGTCCCT**CAAGTCCTTCC**AGCAGCAGCAGCAGCAGCAGCAGCAGCAGCAGCAGCAGCAGCAGCAGCAGCAGCAGCAGCAGCAGCAGCAGCAGCAGCAGCAGCAGCAGCAGCAGCAGCAGCAGCAGCAGCAGCAGCAGCAGCAGCAG**CAACAGCCG**CCACCGCCGCCGCCGCCGCCGCCGCCTCCTCAGCTTCCTCAGCCGCCGCCGCAGGCACAGCCGCTGCTGCCTCAGC
GCTGATGAAGGCCTTCGAGTCCCT**CAAGTCCTTCC**A-------------------------------------------------------------------------------------------------------------CAGCAGCAGCA---------------------GCCGCCGCCGCCGCCTCCTCAGCTTCCTCAGCCGCCGCCGCAGGCACAGCCGCTGCTGCCTCAGC 242 287 DEL
644 25.0
GCTGATGAAGGCCTTCGAGTCCCT**CAAGTCCTTCC**AGCAGCAGCAGCAGCAGCAGCAGCAGCAGCAGCAGCAGCAGCAGCAGCAGCAGCAGCAGCAGCAGCAGCAGCAGCAGCAGCAGCAGCAGCAGCAGCAGCAGCAGCAGCAGCAGCAGCAGCAGCAGCAG**CAACAGCCG**CCACCGCCGCCGCCGCCGCCGCCGCCTCCTCAGCTTCCTCAGCCGCCGCCGCAGGCACAGCCGCTGCTGCCTCAGC
GCTGATGAAGGCCTTCGAGTCCCTCA------------------------------------------------------AGCAGCAGCAGCAGCAGCAGCAGCAGCAGCAGCAGCAGCAGCAGCAGCAGCAGCAGCAGCAGCAGCAGCAGCAGCAGCAGCAG**CAACAGCCG**CCACCGCCGCCGCCGCCGCCGCCGCCTCCTCAGCTTCCTCAGCCGCCGCCGCAGGCACAGCCGCTGCTGCCTCAGC 242 286 DEL
749 17.0
GCTGATGAAGGCCTTCGAGTCCCT**CAAGTCCTTCC**AGCAGCAGCAGCAGCAGCAGCAGCAGCAGCAGCAGCAGCAGCAGCAGCAGCAGCAGCAGCAGCAGCAGCAGCAGCAGCAGCAGCAGCAGCAGCAGCAGCAGCAGCAGCAGCAGCAGCAGCAGCAGCAG**CAACAGCCG**CCACCGCCGCCGCCGCCGCCGCCGCCTCCTCAGCTTCCTCAGCCGCCGCCGCAGGCACAGCCGCTGCTGCCTCAGC
GCTGATGAAGGCCTTCGAGTCCCT**CAAGTCCTTCC**A--------------------------------------------------------------------AGCAGCAGCAGCAGCAGCAGCAGCAGCAGCAGCAGCAGCAGCAGCAGCAGCAG**CAACAGCCG**CCACCGCCGCCGCCGCCGCCGCCGCCTCCTCAGCTTCCTCAGCCGCCGCCGCAGGCACAGCCGCTGCTGCCTCAGC 242 281 DEL
945 11.0
GCTGATGAAGGCCTTCGAGTCCCT**CAAGTCCTTCC**AGCAGCAGCAGCAGCAGCAGCAGCAGCAGCAGCAGCAGCAGCAGCAGCAGCAGCAGCAGCAGCAGCAGCAGCAGCAGCAGCAGCAGCAGCAGCAGCAGCAGCAGCAGCAGCAGCAGCAGCAGCAGCAG**CAACAGCCG**CCACCGCCGCCGCCGCCGCCGCCGCCTCCTCAGCTTCCTCAGCCGCCGCCGCAGGCACAGCCGCTGCTGCCTCAGC
GCTGATGAAGGCCTTCGAGTCCCTCA------------------------------------------------------------------------------------------------AGCAGCAGCAGCAGCAGCAGCAGCAGCAGCAGCAG**CAACAGCCG**CCACCGCCGCCGCCGCCGCCGCCGCCTCCTCAGCTTCCTCAGCCGCCGCCGCAGGCACAGCCGCTGCTGCCTCAGC 242 278 DEL
491 39.0
GCTGATGAAGGCCTTCGAGTCCCT**CAAGTCCTTCC**AGCAGCAGCAGCAGCAGCAGCAGCAGCAGCAGCAGCAGCAGCAGCAGCAGCAGCAGCAGCAGCAGCAGCAGCAGCAGCAGCAGCAGCAGCAGCAGCAGCAGCAGCAGCAGCAGCAGCAGCAGCAGCAG**CAACAGCCG**CCACCGCCGCCGCCGCCGCCGCCGCCTCCTCAGCTTCCTCAGC
GCTGATGAAGGCCTTCGAGTCCCT**CAAGTCCTTCC**A--AGCAGCAGCAGCAGCAGCAGCAGCAGCAGCAGCAGCAGCAGCAGCAGCAGCAGCAGCAGCAGCAGCAGCAGCAGCAGCAGCAGCAGCAGCAGCAGCAGCAGCAGCAGCAGCAGCAGCAGCAGCAG**CAACAGCCG**CCACCGCCGCCGCCGCCGCCGCCGCCTCCTCAGCAGCCTCAGC 209 275 DEL
1068 3.0
GCTGATGAAGGCCTTCGAGTCCCT**CAAGTCCTTC**CAGCAGCAGCAGCAGCAGCAGCAGCAGCAGCAGCAGCAGCAGCAGCAGCAGCAGCAGCAGCAGCAGCAGCAGCAGCAGCAGCAGCAGCAGCAGCAGCAGCAGCAGCAGCAGCAGCAGCAGCAGCAGCAG**CAACAGCCG**CCACCGCCGCCGCCGCCGCCGCCGCCTCCTCAGCTTCCTCAGCCGCCGCCGCAGGCACAGCCGCTGCTGCCTCAGC
GCTGATGAAGGCCTTCGAGTCCCTCAAGTCCTTC------------------------------------------------------------------------------------------------------------------CAGCAGCAG**CAACAGCCG**CCACCGCCGCCGCCGCCGCCGCCGCCTCCTCAGCTTCCTCAGCCGCCGCCGCAGGCACAGCCGCTGCTGCCTCAGC 242 264 DEL
1509 0.0
GCTGATGAAGGCCTTCGAGTCCCT**CAAGTCCTTCC**AGCAGCAGCAGCAGCAGCAGCAGCAGCAGCAGCAGCAGCAGCAGCAGCAGCAGCAGCAGCAGCAGCAGCAGCAGCAGCAGCAGCAGCAGCAGCAGCAGCAGCAGCAGCAGCAGCAGCAGCAGCAGCAG**CAACAGCCG**CCACCGCCGCCGCCGCCGCCGCCGCCTCCTCAGCTTCCTCAGCCGCCGCCGCAGGCACAGCCGCTGCTGCCTCAGC
GCTGATGAAGGCCTTCGAGTCCCTCAAGTCCTT-------------------------------------------------------------------------------------------------------------------------------------------------------------------------------CCGCCGCCGCAGGCACAGCCGCTGCTGCCTCAGC 242 256 DEL
510 27.0
GCTGATGAAGGCCTTCGAGTCCCT**CAAGTCCTTCC**AGCAGCAGCAGCAGCAGCAGCAGCAGCAGCAGCAGCAGCAGCAGCAGCAGCAGCAGCAGCAGCAGCAGCAGCAGCAGCAGCAGCAGCAGCAGCAGCAGCAGCAGCAGCAGCAGCAGCAGCAGCAGCAG**CAACAGCCG**CCACCGCCGCCGCCGCCGCCGCCGCCTCCTCAGCTTCCTCAGCCGCCGCCGCAGGCACAGCCGCTGCTGCCTCAGC
GCTGATGAAGGCCTTCGAGTCCCT**CAAGTCCTTCC**A--------------------------------------AGCAGCAGCAGCAGCAGCAGCAGCAGCAGCAGCAGCAGCAGCAGCAGCAGCAGCAGCAGCAGCAGCAGCAGCAGCAGCAGCAGCAG**CAACAGCCG**CCACCGCCGCCGCCGCCGCCGCCGCCTCCTCAGCTTCCTCAGCCGCCGCCGCAGGCACAGCCGCTGCTGCCTCAGC 242 251 DEL
817 17.0
GCTGATGAAGGCCTTCGAGTCCCT**CAAGTCCTTCC**AGCAGCAGCAGCAGCAGCAGCAGCAGCAGCAGCAGCAGCAGCAGCAGCAGCAGCAGCAGCAGCAGCAGCAGCAGCAGCAGCAGCAGCAGCAGCAGCAGCAGCAGCAGCAGCAGCAGCAGCAGCAGCAG**CAACAGCCG**CCACCGCCGCCGCCGCCGCCGCCGCCTCCTCAGCTTCCTCAGCCGCCGCCGCAGGCACAGCCGCTGCTGCCTCAGC
GCTGATGAAGGCCTTCGAGTCCCTCAAGT-----------------------------------------------------------------------------CAGCAGCAGCAGCAGCAGCAGCAGCAGCAGCAGCAGCAGCAGCAGCAGCAG**CAACAGCCG**CCACCGCCGCCGCCGCCGCCGCCGCCTCCTCAGCTTCCTCAGCCGCCGCCGCAGGCACAGCCGCTGCTGCCTCAGC 242 249 DEL
1514 0.0
GCTGATGAAGGCCTTCGAGTCCCT**CAAGTCCTTCC**AGCAGCAGCAGCAGCAGCAGCAGCAGCAGCAGCAGCAGCAGCAGCAGCAGCAGCAGCAGCAGCAGCAGCAGCAGCAGCAGCAGCAGCAGCAGCAGCAGCAGCAGCAGCAGCAGCAGCAGCAGCAGCAG**CAACAGCCG**CCACCGCCGCCGCCGCCGCCGCCGCCTCCTCAGCTTCCTCAGCCGCCGCCGCAGGCACAGCCGCTGCTGCCTCAGC
GCTGATGAAGGCCTTCGAGTCCCTCAAGTCC------------------------------------------------------------------------------------------------------------------------------------------------------------------------------------------------GCCGCCGCCGCAGGCACAGCCGCTGCTGCCTCAGC 242 241 DEL
834 13.0
GCTGATGAAGGCCTTCGAGTCCCT**CAAGTCCTTCC**AGCAGCAGCAGCAGCAGCAGCAGCAGCAGCAGCAGCAGCAGCAGCAGCAGCAGCAGCAGCAGCAGCAGCAGCAGCAGCAGCAGCAGCAGCAGCAGCAGCAGCAGCAGCAGCAGCAGCAGCAGCAGCAG**CAACAGCCG**CCACCGCCGCCGCCGCCGCCGCCGCCTCCTCAGCTTCCTCAGCCGCCGCCGCAGGCACAGCCGCTGCTGCCTCAGC
GCTGATGAAGGCCTTCGAGTCCCT**CAAGTCCTTCC**A--------------------------------------------------------------------------------AGCAGCAGCAGCAGCAGCAGCAGCAGCAGCAGCAGCAGCAG**CAACAGCCG**CCACCGCCGCCGCCGCCGCCGCCGCCTCCTCAGCTTCCTCAGCCGCCGCCGCAGGCACAGCCGCTGCTGCCTCAGC 242 240 DEL
1132 0.0
GCTGATGAAGGCCTTCGAGTCCCT**CAAGTCCTTCC**AGCAGCAGCAGCAGCAGCAGCAGCAGCAGCAGCAGCAGCAGCAGCAGCAGCAGCAGCAGCAGCAGCAGCAGCAGCAGCAGCAGCAGCAGCAGCAGCAGCAGCAGCAGCAGCAGCAGCAGCAGCAGCAG**CAACAGCCG**CCACCGCCGCCGCCGCCGCCGCCGCCTCCTCAGCTTCCTCAGCCGCCGCCGCAGGCACAGCCGCTGCTGCCTCAGC
GCTGATGAAGGCCTTCGAGTCCCT**CAAGTCCTTCC**----------------------------------------------------------------------------------------------------------------------------ACAGCCGCCACCGCCGCCGCCGCCGCCGCCGCCTCCTCAGCTTCCTCAGCCGCCGCCGCAGGCACAGCCGCTGCTGCCTCAGC 242 240 DEL
934 13.0
GCTGATGAAGGCCTTCGAGTCCCT**CAAGTCCTTCC**AGCAGCAGCAGCAGCAGCAGCAGCAGCAGCAGCAGCAGCAGCAGCAGCAGCAGCAGCAGCAGCAGCAGCAGCAGCAGCAGCAGCAGCAGCAGCAGCAGCAGCAGCAGCAGCAGCAGCAGCAGCAGCAG**CAACAGCCG**CCACCGCCGCCGCCGCCGCCGCCGCCTCCTCAGCTTCCTCAGCCGCCGCCGCAGGCACAGCCGCTGCTGCCTCAGC
GCTGATGAAGGCCTTCGAGTCCCT----------------------------------------------------------------------------------------------CAGCAGCAGCAGCAGCAGCAGCAGCAGCAGCAGCAGCAG**CAACAGCCG**CCACCGCCGCCGCCGCCGCCGCCGCCTCCTCAGCTTCCTCAGCCGCCGCCGCAGGCACAGCCGCTGCTGCCTCAGC 242 232 DEL
561 25.0
GCTGATGAAGGCCTTCGAGTCCCT**CAAGTCCTTCC**AGCAGCAGCAGCAGCAGCAGCAGCAGCAGCAGCAGCAGCAGCAGCAGCAGCAGCAGCAGCAGCAGCAGCAGCAGCAGCAGCAGCAGCAGCAGCAGCAGCAGCAGCAGCAGCAGCAGCAGCAGCAGCAG**CAACAGCCG**CCACCGCCGCCGCCGCCGCCGCCGCCTCCTCAGCTTCCTCAGCCGCCGCCGCAGGCACAGCCGCTGCTGCCTCAGC
GCTGATGAAGGCCTTCGAGTCCCT**CAAGTCCTTCC**A--------------------------------------------AGCAGCAGCAGCAGCAGCAGCAGCAGCAGCAGCAGCAGCAGCAGCAGCAGCAGCAGCAGCAGCAGCAGCAGCAGCAGCAG**CAACAGCCG**CCACCGCCGCCGCCGCCGCCGCCGCCTCCTCAGCTTCCTCAGCCGCCGCCGCAGGCACAGCCGCTGCTGCCTCAGC 242 219 DEL
503 32.0
GCTGATGAAGGCCTTCGAGTCCCT**CAAGTCCTTCC**AGCAGCAGCAGCAGCAGCAGCAGCAGCAGCAGCAGCAGCAGCAGCAGCAGCAGCAGCAGCAGCAGCAGCAGCAGCAGCAGCAGCAGCAGCAGCAGCAGCAGCAGCAGCAGCAGCAGCAGCAGCAGCAG**CAACAGCCG**CCACCGCCGCCGCCGCCGCCGCCGCCTCCTCAGCTTCCTCAGCCGCCGCCGCAGGCACAGCCGCTGCTGCCTCAGC
GCTGATGAAGGCCTTCGAGTCCCT-------------------------------------CAGCAGCAGCAGCAGCAGCAGCAGCAGCAGCAGCAGCAGCAGCAGCAGCAGCAGCAGCAGCAGCAGCAGCAGCAGCAGCAGCAGCAGCAGCAGCAG**CAACAGCCG**CCACCGCCGCCGCCGCCGCCGCCGCCTCCTCAGCTTCCTCAGCCGCCGCCGCAGGCACAGCCGCTGCTGCCTCAGC 242 215 DEL
708 19.0
GCTGATGAAGGCCTTCGAGTCCCT**CAAGTCCTTCC**AGCAGCAGCAGCAGCAGCAGCAGCAGCAGCAGCAGCAGCAGCAGCAGCAGCAGCAGCAGCAGCAGCAGCAGCAGCAGCAGCAGCAGCAGCAGCAGCAGCAGCAGCAGCAGCAGCAGCAGCAGCAGCAG**CAACAGCCG**CCACCGCCGCCGCCGCCGCCGCCGCCTCCTCAGCTTCCTCAGCCGCCGCCGCAGGCACAGCCGCTGCTGCCTCAGC
GCTGATGAAGGCCTTCGAGTCCCT**CAAGTCCTTCC**A--------------------------------------------------------------AGCAGCAGCAGCAGCAGCAGCAGCAGCAGCAGCAGCAGCAGCAGCAGCAGCAGCAGCAGCAG**CAACAGCCG**CCACCGCCGCCGCCGCCGCCGCCGCCTCCTCAGCTTCCTCAGCCGCCGCCGCAGGCACAGCCGCTGCTGCCTCAGC 242 209 DEL
330 35.0
GCTGATGAAGGCCTTCGAGTCCCT**CAAGTCCTTCC**AGCAGCAGCAGCAGCAGCAGCAGCAGCAGCAGCAGCAGCAGCAGCAGCAGCAGCAGCAGCAGCAGCAGCAGCAGCAGCAGCAGCAGCAGCAGCAGCAGCAGCAGCAGCAGCAGCAGCAGCAGCAGCAG**CAACAGCCG**CCACCGCCGCCGCCGCCGCCGCCGCCTCCTCAGCTTCCTCAGCCGCCGCCGCAGGCACAGCCGCTGCTGCCTCAGC
GCTGATGAAGGCCTTCGAGTCCCT**CAAGTCCTTCC**A----------------CAGCAGCAGCAGCAGCAGCAGCAGCAGCAGCAGCAGCAGCAGCAGCAGCAGCAGCAGCAGCAGCAGCAGCAGCAGCAGCAGCAGCAGCAGCAGCAGCAGCAGCAG**CAACAGCCG**CCACCGCCGCCGCCGCCGCCGCCGCCTCCTCAGCTTCCTCAGCCGCCGCCGCAGGCACAGCCGCTGCTGCCTCAGC 242 209 DEL
1478 0.0
GCTGATGAAGGCCTTCGAGTCCCT**CAAGTCCTTCC**AGCAGCAGCAGCAGCAGCAGCAGCAGCAGCAGCAGCAGCAGCAGCAGCAGCAGCAGCAGCAGCAGCAGCAGCAGCAGCAGCAGCAGCAGCAGCAGCAGCAGCAGCAGCAGCAGCAGCAGCAGCAGCAG**CAACAGCCG**CCACCGCCGCCGCCGCCGCCGCCGCCTCCTCAGCTTCCTCAGCCGCCGCCGCAGGCACAGCCGCTGCTGCCTCAGC
GCTGATGAAGGCCTTCGAGTC-----------------------------------------------------------------------------------------------------------------------------------------------------------------------CCTCCTCAGCTTCCTCAGCCGCCGCCGCAGGCACAGCCGCTGCTGCCTCAGC 242 207 DEL
384 32.0
GCTGATGAAGGCCTTCGAGTCCCT**CAAGTCCTTCC**AGCAGCAGCAGCAGCAGCAGCAGCAGCAGCAGCAGCAGCAGCAGCAGCAGCAGCAGCAGCAGCAGCAGCAGCAGCAGCAGCAGCAGCAGCAGCAGCAGCAGCAGCAGCAGCAGCAGCAGCAGCAGCAG**CAACAGCCG**CCACCGCCGCCGCCGCCGCCGCCGCCTCCTCAGCTTCCTCAGCCGCCGCCGCAGGCACAGCCGCTGCTGCCTCAGC
GCTGATGAAGGCCTTCGAGTCCCT**CAAGTCCTTCC**AG----------------------AGCAGCAGCAGCAGCAGCAGCAGCAGCAGCAGCAGCAGCAGCAGCAGCAGCAGCAGCAGCAGCAGCAGCAGCAGCAGCAGCAGCAGCAGCAGCAGCAG**CAACAGCCG**CCACCGCCGCCGCCGCCGCCGCCGCCTCCTCAGCTTCCTCAGCCGCCGCCGCAGGCACAGCCGCTGCTGCCTCAGC 242 200 DEL
1479 0.0
GCTGATGAAGGCCTTCGAGTCCCT**CAAGTCCTTCC**AGCAGCAGCAGCAGCAGCAGCAGCAGCAGCAGCAGCAGCAGCAGCAGCAGCAGCAGCAGCAGCAGCAGCAGCAGCAGCAGCAGCAGCAGCAGCAGCAGCAGCAGCAGCAGCAGCAGCAGCAGCAGCAG**CAACAGCCG**CCACCGCCGCCGCCGCCGCCGCCGCCTCCTCAGCTTCCTCAGCCGCCGCCGCAGGCACAGCCGCTGCTGCCTCAGC
GCTGATGAAGGCCTTCGAGTCCCT**CAAGTCCTTCC**GCC-----------------------------------------------------------------------------------------------------------------------------------------------------------------------GCCGCCGCCGCAGGCACAGCCGCTGCTGCCTCAGC 242 195 DEL
683 20.0
GCTGATGAAGGCCTTCGAGTCCCT**CAAGTCCTTCC**AGCAGCAGCAGCAGCAGCAGCAGCAGCAGCAGCAGCAGCAGCAGCAGCAGCAGCAGCAGCAGCAGCAGCAGCAGCAGCAGCAGCAGCAGCAGCAGCAGCAGCAGCAGCAGCAGCAGCAGCAGCAGCAG**CAACAGCCG**CCACCGCCGCCGCCGCCGCCGCCGCCTCCTCAGCTTCCTCAGCCGCCGCCGCAGGCACAGCCGCTGCTGCCTCAGC
GCTGATGAAGGCCTTCGAGTCCCT**CAAGTCCTTCC**A-----------------------------------------------------------AGCAGCAGCAGCAGCAGCAGCAGCAGCAGCAGCAGCAGCAGCAGCAGCAGCAGCAGCAGCAGCAG**CAACAGCCG**CCACCGCCGCCGCCGCCGCCGCCGCCTCCTCAGCTTCCTCAGCCGCCGCCGCAGGCACAGCCGCTGCTGCCTCAGC 242 194 DEL
913 12.0
GCTGATGAAGGCCTTCGAGTCCCT**CAAGTCCTTCC**AGCAGCAGCAGCAGCAGCAGCAGCAGCAGCAGCAGCAGCAGCAGCAGCAGCAGCAGCAGCAGCAGCAGCAGCAGCAGCAGCAGCAGCAGCAGCAGCAGCAGCAGCAGCAGCAGCAGCAGCAGCAGCAG**CAACAGCCG**CCACCGCCGCCGCCGCCGCCGCCGCCTCCTCAGCTTCCTCAGCCGCCGCCGCAGGCACAGCCGCTGCTGCCTCAGC
GCTGATGAAGGCCTTCGAGTCCCTCAAGTC-------------------------------------------------------------------------------------------CAGCAGCAGCAGCAGCAGCAGCAGCAGCAGCAGCAG**CAACAGCCG**CCACCGCCGCCGCCGCCGCCGCCGCCTCCTCAGCTTCCTCAGCCGCCGCCGCAGGCACAGCCGCTGCTGCCTCAGC 242 193 DEL
482 31.0
GCTGATGAAGGCCTTCGAGTCCCT**CAAGTCCTTCC**AGCAGCAGCAGCAGCAGCAGCAGCAGCAGCAGCAGCAGCAGCAGCAGCAGCAGCAGCAGCAGCAGCAGCAGCAGCAGCAGCAGCAGCAGCAGCAGCAGCAGCAGCAGCAGCAGCAGCAGCAGCAGCAG**CAACAGCCG**CCACCGCCGCCGCCGCCGCCGCCGCCTCCTCAGCTTCCTCAGCCGCCGCCGCAGGCACAGCCGCTGCTGCCTCAGC
GCTGATGAAGGCCTTCGAGTCCCTCAAGTC----------------------------------CAGCAGCAGCAGCAGCAGCAGCAGCAGCAGCAGCAGCAGCAGCAGCAGCAGCAGCAGCAGCAGCAGCAGCAGCAGCAGCAGCAGCAGCAGCAG**CAACAGCCG**CCACCGCCGCCGCCGCCGCCGCCGCCTCCTCAGCTTCCTCAGCCGCCGCCGCAGGCACAGCCGCTGCTGCCTCAGC 242 192 DEL
374 32.0
GCTGATGAAGGCCTTCGAGTCCCT**CAAGTCCTTCC**AGCAGCAGCAGCAGCAGCAGCAGCAGCAGCAGCAGCAGCAGCAGCAGCAGCAGCAGCAGCAGCAGCAGCAGCAGCAGCAGCAGCAGCAGCAGCAGCAGCAGCAGCAGCAGCAGCAGCAGCAGCAGCAG**CAACAGCCG**CCACCGCCGCCGCCGCCGCCGCCGCCTCCTCAGCTTCCTCAGCCGCCGCCGCAGGCACAGCCGCTGCTGCCTCAGC
GCTGATGAAGGCCTTCGAGTCCCT**CAAGTCCTTCC**AGC---------------------TGCAGCAGCAGCAGCAGCAGCAGCAGCAGCAGCAGCAGCAGCAGCAGCAGCAGCAGCAGCAGCAGCAGCAGCAGCAGCAGCAGCAGCAGCAGCAGCAG**CAACAGCCG**CCACCGCCGCCGCCGCCGCCGCCGCCTCCTCAGCTTCCTCAGCCGCCGCCGCAGGCACAGCCGCTGCTGCCTCAGC 242 189 DEL
584 24.0
GCTGATGAAGGCCTTCGAGTCCCT**CAAGTCCTTCC**AGCAGCAGCAGCAGCAGCAGCAGCAGCAGCAGCAGCAGCAGCAGCAGCAGCAGCAGCAGCAGCAGCAGCAGCAGCAGCAGCAGCAGCAGCAGCAGCAGCAGCAGCAGCAGCAGCAGCAGCAGCAGCAG**CAACAGCCG**CCACCGCCGCCGCCGCCGCCGCCGCCTCCTCAGCTTCCTCAGCCGCCGCCGCAGGCACAGCCGCTGCTGCCTCAGC
GCTGATGAAGGCCTTCGAGTCCCT**CAAGTCCTTCC**A-----------------------------------------------AGCAGCAGCAGCAGCAGCAGCAGCAGCAGCAGCAGCAGCAGCAGCAGCAGCAGCAGCAGCAGCAGCAGCAGCAGCAG**CAACAGCCG**CCACCGCCGCCGCCGCCGCCGCCGCCTCCTCAGCTTCCTCAGCCGCCGCCGCAGGCACAGCCGCTGCTGCCTCAGC 242 181 DEL
353 38.0
GCTGATGAAGGCCTTCGAGTCCCT**CAAGTCCTTCC**AGCAGCAGCAGCAGCAGCAGCAGCAGCAGCAGCAGCAGCAGCAGCAGCAGCAGCAGCAGCAGCAGCAGCAGCAGCAGCAGCAGCAGCAGCAGCAGCAGCAGCAGCAGCAGCAGCAGCAGCAGCAGCAG**CAACAGCCG**CCACCGCCGCCGCCGCCGCCGCCGCCTCCTCAGCTTCCTCAGCCGCCGCCGCAGGCACAGCCGCTGCTGCCTCAGC
GCTGATGAAGGCCTTCGAGTCCCT-------------------CAGCAGCAGCAGCAGCAGCAGCAGCAGCAGCAGCAGCAGCAGCAGCAGCAGCAGCAGCAGCAGCAGCAGCAGCAGCAGCAGCAGCAGCAGCAGCAGCAGCAGCAGCAGCAGCAGCAG**CAACAGCCG**CCACCGCCGCCGCCGCCGCCGCCGCCTCCTCAGCTTCCTCAGCCGCCGCCGCAGGCACAGCCGCTGCTGCCTCAGC 242 177 DEL
729 18.0
GCTGATGAAGGCCTTCGAGTCCCT**CAAGTCCTTCC**AGCAGCAGCAGCAGCAGCAGCAGCAGCAGCAGCAGCAGCAGCAGCAGCAGCAGCAGCAGCAGCAGCAGCAGCAGCAGCAGCAGCAGCAGCAGCAGCAGCAGCAGCAGCAGCAGCAGCAGCAGCAGCAG**CAACAGCCG**CCACCGCCGCCGCCGCCGCCGCCGCCTCCTCAGCTTCCTCAGCCGCCGCCGCAGGCACAGCCGCTGCTGCCTCAGC
GCTGATGAAGGCCTTCGAGTCCCT**CAAGTCCTTCC**A-----------------------------------------------------------------AGCAGCAGCAGCAGCAGCAGCAGCAGCAGCAGCAGCAGCAGCAGCAGCAGCAGCAGCAG**CAACAGCCG**CCACCGCCGCCGCCGCCGCCGCCGCCTCCTCAGCTTCCTCAGCCGCCGCCGCAGGCACAGCCGCTGCTGCCTCAGC 242 175 DEL
457 32.0
GCTGATGAAGGCCTTCGAGTCCCT**CAAGTCCTTCC**AGCAGCAGCAGCAGCAGCAGCAGCAGCAGCAGCAGCAGCAGCAGCAGCAGCAGCAGCAGCAGCAGCAGCAGCAGCAGCAGCAGCAGCAGCAGCAGCAGCAGCAGCAGCAGCAGCAGCAGCAGCAGCAG**CAACAGCCG**CCACCGCCGCCGCCGCCGCCGCCGCCTCCTCAGCTTCCTCAGCCGCCGCCGCAGGCACAGCCGCTGCTGCCTCAGC
GCTGATGAAGGCCTTCGAGTCCCTCAAGTC-------------------------------CAGCAGCAGCAGCAGCAGCAGCAGCAGCAGCAGCAGCAGCAGCAGCAGCAGCAGCAGCAGCAGCAGCAGCAGCAGCAGCAGCAGCAGCAGCAGCAG**CAACAGCCG**CCACCGCCGCCGCCGCCGCCGCCGCCTCCTCAGCTTCCTCAGCCGCCGCCGCAGGCACAGCCGCTGCTGCCTCAGC 242 175 DEL
426 34.0
GCTGATGAAGGCCTTCGAGTCCCT**CAAGTCCTTCC**AGCAGCAGCAGCAGCAGCAGCAGCAGCAGCAGCAGCAGCAGCAGCAGCAGCAGCAGCAGCAGCAGCAGCAGCAGCAGCAGCAGCAGCAGCAGCAGCAGCAGCAGCAGCAGCAGCAGCAGCAGCAGCAG**CAACAGCCG**CCACCGCCGCCGCCGCCGCCGCCGCCTCCTCAGCTTCCTCAGCCGCCGCCGCAGGCACAGCCGCTGCTGCCTCAGC
GCTGATGAAGGCCTTCGAGTCCCTCA---------------------------AGCAGCAGCAGCAGCAGCAGCAGCAGCAGCAGCAGCAGCAGCAGCAGCAGCAGCAGCAGCAGCAGCAGCAGCAGCAGCAGCAGCAGCAGCAGCAGCAGCAGCAGCAG**CAACAGCCG**CCACCGCCGCCGCCGCCGCCGCCGCCTCCTCAGCTTCCTCAGCCGCCGCCGCAGGCACAGCCGCTGCTGCCTCAGC 242 173 DEL
73 43.0
GCTGATGAAGGCCTTCGAGTCCCTCAAGTCCTTC--------CAGCAGCAGCAGCAGCAGCAGCAGCAGCAGCAGCAGCAGCAGCAGCAGCAGCAGCAGCAGCAGCAGCAGCAGCAGCAGCAGCAGCAGCAGCAGCAGCAGCAGCAGCAGCAGCAGCAGCAGCAGCAGCAG**CAACAGCCG**CCACCGCCGCCGCCGCCGCCGCCGCCTCCTCAGCTTCCTCAGCCGCCGCCGCAGGCACAGCCGCTGCTGCCTCAGC
GCTGATGAAGGCCTTCGAGTCCCT**CAAGTCCTTCC**ACAGCAGCAGCAGCAGCAGCAGCAGCAGCAGCAGCAGCAGCAGCAGCAGCAGCAGCAGCAGCAGCAGCAGCAGCAGCAGCAGCAGCAGCAGCAGCAGCAGCAGCAGCAGCAGCAGCAGCAGCAGCAGCAGCAGCAG**CAACAGCCG**CCACCGCCGCCGCCGCCGCCGCCGCCTCCTCAGCTTCCTCAGCCGCCGCCGCAGGCACAGCCGCTGCTGCCTCAGC 242 170 INS
543 29.0
GCTGATGAAGGCCTTCGAGTCCCT**CAAGTCCTTCC**AGCAGCAGCAGCAGCAGCAGCAGCAGCAGCAGCAGCAGCAGCAGCAGCAGCAGCAGCAGCAGCAGCAGCAGCAGCAGCAGCAGCAGCAGCAGCAGCAGCAGCAGCAGCAGCAGCAGCAGCAGCAGCAG**CAACAGCCG**CCACCGCCGCCGCCGCCGCCGCCGCCTCCTCAGCTTCCTCAGCCGCCGCCGCAGGCACAGCCGCTGCTGCCTCAGC
GCTGATGAAGGCCTTCGAGTCCCTCA------------------------------------------AGCAGCAGCAGCAGCAGCAGCAGCAGCAGCAGCAGCAGCAGCAGCAGCAGCAGCAGCAGCAGCAGCAGCAGCAGCAGCAGCAGCAGCAGCAG**CAACAGCCG**CCACCGCCGCCGCCGCCGCCGCCGCCTCCTCAGCTTCCTCAGCCGCCGCCGCAGGCACAGCCGCTGCTGCCTCAGC 242 168 DEL
322 40.0
GCTGATGAAGGCCTTCGAGTCCCT**CAAGTCCTTCC**AGCAGCAGCAGCAGCAGCAGCAGCAGCAGCAGCAGCAGCAGCAGCAGCAGCAGCAGCAGCAGCAGCAGCAGCAGCAGCAGCAGCAGCAGCAGCAGCAGCAGCAGCAGCAGCAGCAGCAGCAGCAGCAG**CAACAGCCG**CCACCGCCGCCGCCGCCGCCGCCGCCTCCTCAGCTTCCTCAGCCGCCGCCGCAGGCACAGCCGCTGCTGCCTCAGC
GCTGATGAAGGCCTTCGAGT---------------AGCAGCAGCAGCAGCAGCAGCAGCAGCAGCAGCAGCAGCAGCAGCAGCAGCAGCAGCAGCAGCAGCAGCAGCAGCAGCAGCAGCAGCAGCAGCAGCAGCAGCAGCAGCAGCAGCAGCAGCAGCAGCAGCAG**CAACAGCCG**CCACCGCCGCCGCCGCCGCCGCCGCCTCCTCAGCTTCCTCAGCCGCCGCCGCAGGCACAGCCGCTGCTGCCTCAGC 242 162 DEL
562 40.0
GCTGATGAAGGCCTTCGAGTCCCT**CAAGTCCTTCC**-AGCAGCAGCAGCAGCAGCAGCAGCAGCAGCAGCAGCAGCAGCAGCAGCAGCAGCAGCAGCAGCAGCAGCAGCAGCAGCAGCAGCAGCAGCAGCAGCAGCAGCAGCAGCAGCAGCAGCAGCAGCAGCAG**CAACAGCCG**CCACCGCCGCCGCCGCCGCCGCCGCCTCCTCAGC
GCTGATGAAGGCCTTCGAGTCCCT**CAAGTCCTTCC**AAGCAGCAGCAGCAGCAGCAGCAGCAGCAGCAGCAGCAGCAGCAGCAGCAGCAGCAGCAGCAGCAGCAGCAGCAGCAGCAGCAGCAGCAGCAGCAGCAGCAGCAGCAGCAGCAGCAGCAGCAGCAGCAG**CAACAGCCG**CCACCGCCGCCGCCGCCGCCGCCG---CCTCAGC 200 157 INS+DEL

```
1532           0.0
GCTGATGAAGGCCTTCGAGTCCCTCAAGTCCTTCCAGCAGCAGCAGCAGCAGCAGCAGCAGCAGCAGCAGCAGCAGCAGCAGCAGCAGCAGCAGCAGCAGCAGCAGCAGCAGCAGCAGCAGCAGCAGCAGCAGCAGCAGCAGCAGCAGCAGCAACAGCCGCCACCGCCGCCGCCGCCGCCGCCGCCTCCTCAGCTTCCTCAGCCGCCGCCGCAGGCACAGCCGCTGCTGCCTCAGC
GCTGATGAAGGCCTTCGAGTCCCTCAAGT-----------------------------------------------------------------------------------------------------------------------------------------------------------------------------------CCGCCGCCGCAGGCACAGCCGCTGCTGCCTCAGC         242          157          DEL
1439           0.0
GCTGATGAAGGCCTTCGAGTCCCTCAAGTCCTTCCAGCAGCAGCAGCAGCAGCAGCAGCAGCAGCAGCAGCAGCAGCAGCAGCAGCAGCAGCAGCAGCAGCAGCAGCAGCAGCAGCAGCAGCAGCAGCAGCAGCAGCAGCAGCAGCAGCAGCAACAGCCGCCACCGCCGCCGCCGCCGCCGCCGCCTCCTCAGCTTCCTCAGCCGCCGCCGCAGGCACAGCCGCTGCTGCCTCAGC
GCTGATGAAGGCCTTCGAGTCCCTCAAGTC----------------------------------------------------------------------------------------------------------------------------------------------------------------CCTCCTCAGCTTCCTCAGCCGCCGCCGCAGGCACAGCCGCTGCTGCCTCAGC         242          156          DEL
637            22.0
GCTGATGAAGGCCTTCGAGTCCCTCAAGTCCTTCCAGCAGCAGCAGCAGCAGCAGCAGCAGCAGCAGCAGCAGCAGCAGCAGCAGCAGCAGCAGCAGCAGCAGCAGCAGCAGCAGCAGCAGCAGCAGCAGCAGCAGCAGCAGCAGCAGCAGCAACAGCCGCCACCGCCGCCGCCGCCGCCGCCGCCTCCTCAGCTTCCTCAGCCGCCGCCGCAGGCACAGCCGCTGCTGCCTCAGC
GCTGATGAAGGCCTTCGAGTCCCTCAAGTCCTTCCA-----------------------------------------------------AGCAGCAGCAGCAGCAGCAGCAGCAGCAGCAGCAGCAGCAGCAGCAGCAGCAGCAGCAGCAGCAGCAGCAGCAACAGCCGCCACCGCCGCCGCCGCCGCCGCCGCCTCCTCAGCTTCCTCAGCCGCCGCCGCAGGCACAGCCGCTGCTGCCTCAGC         242          155          DEL
745            22.0
GCTGATGAAGGCCTTCGAGTCCCTCAAGTCCTTCCAGCAGCAGCAGCAGCAGCAGCAGCAGCAGCAGCAGCAGCAGCAGCAGCAGCAGCAGCAGCAGCAGCAGCAGCAGCAGCAGCAGCAGCAGCAGCAGCAGCAGCAGCAGCAGCAGCAGCAACAGCCGCCACCGCCGCCGCCGCCGCCGCCGCCTCCTCAGCTTCCTCAGCCGCCGCCGCAGGCACAGCCGCTGCTGCCTCAGC
GCTGATGAAGGCCTTCGAGTCCCT-------------------------------------------------------------------CAGCAGCAGCAGCAGCAGCAGCAGCAGCAGCAGCAGCAGCAGCAGCAGCAGCAGCAGCAGCAGCAGCAACAGCCGCCACCGCCGCCGCCGCCGCCGCCGCCTCCTCAGCTTCCTCAGCCGCCGCCGCAGGCACAGCCGCTGCTGCCTCAGC         242          148          DEL
949            1.0
GCTGATGAAGGCCTTCGAGTCCCTCAAGTCCTTCCAGCAGCAGCAGCAGCAGCAGCAGCAGCAGCAGCAGCAGCAGCAGCAGCAGCAGCAGCAGCAGCAGCAGCAGCAGCAGCAGCAGCAGCAGCAGCAGCAGCAGCAGCAGCAGCAGCAGCAACAGCCGCCACCGCCGCCGCCGCCGCCGCCGCCTCCTCAGCTTCCTCAGCCGCCGCCGCAGGCACAGCCGCTGCTGCCTCAGC
GCTGATGAAGGCCTTCGAGTCCCTCAAGTCCTTCCGCCCTGAGCAGCAACAGCTG-------------------------------------------------------------------------------------------------AGCAGCAACAGCCGCCACCGCCGCCGCCGCCGCCGCCGCCTCCTCAGCTTCCTCAGCCGCCGCCGCAGGCACAGCCGCTGCTGCCTCAGC         242          147          DEL
611            23.0
GCTGATGAAGGCCTTCGAGTCCCTCAAGTCCTTCCAGCAGCAGCAGCAGCAGCAGCAGCAGCAGCAGCAGCAGCAGCAGCAGCAGCAGCAGCAGCAGCAGCAGCAGCAGCAGCAGCAGCAGCAGCAGCAGCAGCAGCAGCAGCAGCAGCAGCAACAGCCGCCACCGCCGCCGCCGCCGCCGCCGCCTCCTCAGCTTCCTCAGCCGCCGCCGCAGGCACAGCCGCTGCTGCCTCAGC
GCTGATGAAGGCCTTCGAGTCCCTCAAGTCCTTCCA--------------------------------------------------AGCAGCAGCAGCAGCAGCAGCAGCAGCAGCAGCAGCAGCAGCAGCAGCAGCAGCAGCAGCAGCAGCAGCAGCAGCAACAGCCGCCACCGCCGCCGCCGCCGCCGCCGCCTCCTCAGCTTCCTCAGCCGCCGCCGCAGGCACAGCCGCTGCTGCCTCAGC         242          139          DEL
511            40.0
GCTGATGAAGGCCTTCGAGTCCCTCAAGTCCTTCC-AGCAGCAGCAGCAGCAGCAGCAGCAGCAGCAGCAGCAGCAGCAGCAGCAGCAGCAGCAGCAGCAGCAGCAGCAGCAGCAGCAGCAGCAGCAGCAGCAGCAGCAGCAGCAGCAGCAGCAACAGCCGCCACCGCCGCCGCCGCCGCCGCCGCCTCCTCAGCTTCCTCAGCCGCCGCCGCAGGCACAGCCGCTGCTGCCTCAGC
GCTGATGAAGGCCTTCGAGTCCCTCAAGTCCTTCCAAGCAGCAGCAGCAGCAGCAGCAGCAGCAGCAGCAGCAGCAGCAGCAGCAGCAGCAGCAGCAGCAGCAGCAGCAGCAGCAGCAGCAGCAGCAGCAGCAGCAGCAGCAGCAGCAGCAGCAACAGCCGCCA---------------------------------------CCGCCGCCGCAGGCACAGCCGCTGCTGCCTCAGC         242          133          INS+DEL
408            31.0
GCTGATGAAGGCCTTCGAGTCCCTCAAGTCCTTCCAGCAGCAGCAGCAGCAGCAGCAGCAGCAGCAGCAGCAGCAGCAGCAGCAGCAGCAGCAGCAGCAGCAGCAGCAGCAGCAGCAGCAGCAGCAGCAGCAGCAGCAGCAGCAGCAGCAGCAACAGCCGCCACCGCCGCCGCCGCCGCCGCCGCCTCCTCAGCTTCCTCAGCCGCCGCCGCAGGCACAGCCGCTGCTGCCTCAGC
GCTGATGAAGGCCTTCGAGTCCCTCAAGTCCTTCCAG-------------------------AGCAGCAGCAGCAGCAGCAGCAGCAGCAGCAGCAGCAGCAGCAGCAGCAGCAGCAGCAGCAGCAGCAGCAGCAGCAGCAGCAGCAGCAGCAGCAGCAACAGCCGCCACCGCCGCCGCCGCCGCCGCCGCCTCCTCAGCTTCCTCAGCCGCCGCCGCAGGCACAGCCGCTGCTGCCTCAGC         242          130          DEL
59             44.0
GCTGATGAAGGCCTTCGAGTCCCTCAAGTCCTTC-------------CAGCAGCAGCAGCAGCAGCAGCAGCAGCAGCAGCAGCAGCAGCAGCAGCAGCAGCAGCAGCAGCAGCAGCAGCAGCAGCAGCAGCAGCAGCAGCAGCAGCAGCAGCAGCAGCAGCAGCAGCAGCAGCAACAGCCGCCACCGCCGCCGCCGCCGCCGCCGCCTCCTCAGCTTCCTCAGCCGCCGCCGCAGGCACAGCCGCTGCTGCCTCAGC
GCTGATGAAGGCCTTCGAGTCCCTCAAGTCCTTCCAAGCAGCAGCAGCAGCAGCAGCAGCAGCAGCAGCAGCAGCAGCAGCAGCAGCAGCAGCAGCAGCAGCAGCAGCAGCAGCAGCAGCAGCAGCAGCAGCAGCAGCAGCAGCAGCAGCAGCAGCAGCAGCAGCAACAGCCGCCACCGCCGCCGCCGCCGCCGCCGCCTCCTCAGCTTCCTCAGCCGCCGCCGCAGGCACAGCCGCTGCTGCCTCAGC           242          129          INS
652            24.0
GCTGATGAAGGCCTTCGAGTCCCTCAAGTCCTTCCAGCAGCAGCAGCAGCAGCAGCAGCAGCAGCAGCAGCAGCAGCAGCAGCAGCAGCAGCAGCAGCAGCAGCAGCAGCAGCAGCAGCAGCAGCAGCAGCAGCAGCAGCAGCAGCAGCAGCAACAGCCGCCACCGCCGCCGCCGCCGCCGCCGCCTCCTCAGCTTCCTCAGCCGCCGCCGCAGGCACAGCCGCTGCTGCCTCAGC
GCTGATGAAGGCCTTCGAGTCCCTCAAGTC-------------------------------------------------------CAGCAGCAGCAGCAGCAGCAGCAGCAGCAGCAGCAGCAGCAGCAGCAGCAGCAGCAGCAGCAGCAGCAGCAGCAACAGCCGCCACCGCCGCCGCCGCCGCCGCCGCCTCCTCAGCTTCCTCAGCCGCCGCCGCAGGCACAGCCGCTGCTGCCTCAGC         242          126          DEL
1216           0.0
GCTGATGAAGGCCTTCGAGTCCCTCAAGTCCTTCCAGCAGCAGCAGCAGCAGCAGCAGCAGCAGCAGCAGCAGCAGCAGCAGCAGCAGCAGCAGCAGCAGCAGCAGCAGCAGCAGCAGCAGCAGCAGCAGCAGCAGCAGCAGCAGCAGCAGCAACAGCCGCCACCGCCGCCGCCGCCGCCGCCGCCTCCTCAGCTTCCTCAGCCGCCGCCGCAGGCACAGCCGCTGCTGCCTCAGC
GCTGATGAAGGCCTTCGAGTCCCT-------------------------------------------------------------------------------------------------------------------------------------CAACAGCCGCCACCACCGCCGCCGCCGCCGCCGCCGCCTCCTCAGCTTCCTCAGCCGCCGCCGCAGGCACAGCCGCTGCTGCCTCAGC         242          126          DEL
628            25.0
GCTGATGAAGGCCTTCGAGTCCCTCAAGTCCTTCCAGCAGCAGCAGCAGCAGCAGCAGCAGCAGCAGCAGCAGCAGCAGCAGCAGCAGCAGCAGCAGCAGCAGCAGCAGCAGCAGCAGCAGCAGCAGCAGCAGCAGCAGCAGCAGCAGCAGCAACAGCCGCCACCGCCGCCGCCGCCGCCGCCGCCTCCTCAGCTTCCTCAGCCGCCGCCGCAGGCACAGCCGCTGCTGCCTCAGC
GCTGATGAAGGCCTTCGAGTCCCTCAAGTC----------------------------------------------------CAGCAGCAGCAGCAGCAGCAGCAGCAGCAGCAGCAGCAGCAGCAGCAGCAGCAGCAGCAGCAGCAGCAGCAGCAGCAACAGCCGCCACCGCCGCCGCCGCCGCCGCCGCCTCCTCAGCTTCCTCAGCCGCCGCCGCAGGCACAGCCGCTGCTGCCTCAGC         242          125          DEL
399            33.0
GCTGATGAAGGCCTTCGAGTCCCTCAAGTCCTTCCAGCAGCAGCAGCAGCAGCAGCAGCAGCAGCAGCAGCAGCAGCAGCAGCAGCAGCAGCAGCAGCAGCAGCAGCAGCAGCAGCAGCAGCAGCAGCAGCAGCAGCAGCAGCAGCAGCAGCAACAGCCGCCACCGCCGCCGCCGCCGCCGCCGCCTCCTCAGCTTCCTCAGCCGCCGCCGCAGGCACAGCCGCTGCTGCCTCAGC
GCTGATGAAGGCCTTCGAGTCCCTCAAGTCCTTC------------------------CAGCAGCAGCAGCAGCAGCAGCAGCAGCAGCAGCAGCAGCAGCAGCAGCAGCAGCAGCAGCAGCAGCAGCAGCAGCAGCAGCAGCAGCAGCAGCAGCAGCAACAGCCGCCACCGCCGCCGCCGCCGCCGCCGCCTCCTCAGCTTCCTCAGCCGCCGCCGCAGGCACAGCCGCTGCTGCCTCAGC         242          123          DEL
585            39.0
GCTGATGAAGGCCTTCGAGTCCCTCAAGTCCTTCCAGCAGCAGCAGCAGCAGCAGCAGCAGCAGCAGCAGCAGCAGCAGCAGCAGCAGCAGCAGCAGCAGCAGCAGCAGCAGCAGCAGCAGCAGCAGCAGCAGCAGCAGCAGCAGCAGCAGCAACAGCCGCCACCGCCGCCGCCGCCGCCGCCGCCTCCTCAGCTTCCTCAGCCGCCGCCGCAGGCACAGCCGCTGCTGCCTCAGC
GCTGATGAAGGCCTTCGAGTCCCTCAAGTCCTTCCA--AGCAGCAGCAGCAGCAGCAGCAGCAGCAGCAGCAGCAGCAGCAGCAGCAGCAGCAGCAGCAGCAGCAGCAGCAGCAGCAGCAGCAGCAGCAGCAGCAGCAGCAGCAGCAGCAGCAACAGCCGCCACCGCCGCCGCCGCCGCCGCCGCCG---CCTCAGCTTCCTCAGCCGCCGCCGCAGNCACAGCCGCTGCTGCCTCAGC         242          121          DEL
883            11.0
GCTGATGAAGGCCTTCGAGTCCCTCAAGTCCTTCCAGCAGCAGCAGCAGCAGCAGCAGCAGCAGCAGCAGCAGCAGCAGCAGCAGCAGCAGCAGCAGCAGCAGCAGCAGCAGCAGCAGCAGCAGCAGCAGCAGCAGCAGCAGCAGCAGCAGCAACAGCCGCCACCGCCGCCGCCGCCGCCGCCGCCTCCTCAGCTTCCTCAGCCGCCGCCGCAGGCACAGCCGCTGCTGCCTCAGC
GCTGATGAAGGCCTTCGAGTCCCTCAAGTCCTTCCA--------------------------------------------------------------------------------------AGCAGCAGCAGCAGCAGCAGCAGCAGCAGCAGCAGCAACAGCCGCCACCGCCGCCGCCGCCGCCGCCGCCTCCTCAGCTTCCTCAGCCGCCGCCGCAGGCACAGCCGCTGCTGCCTCAGC         242          115          DEL
738            21.0
GCTGATGAAGGCCTTCGAGTCCCTCAAGTCCTTCCAGCAGCAGCAGCAGCAGCAGCAGCAGCAGCAGCAGCAGCAGCAGCAGCAGCAGCAGCAGCAGCAGCAGCAGCAGCAGCAGCAGCAGCAGCAGCAGCAGCAGCAGCAGCAGCAGCAGCAACAGCCGCCACCGCCGCCGCCGCCGCCGCCGCCTCCTCAGCTTCCTCAGCCGCCGCCGCAGGCACAGCCGCTGCTGCCTCAGC
GCTGATGAAGGCCTTCGAGTCCCTCA------------------------------------------------------------------AGCAGCAGCAGCAGCAGCAGCAGCAGCAGCAGCAGCAGCAGCAGCAGCAGCAGCAGCAGCAGCAGCAGCAACAGCCGCCACCGCCGCCGCCGCCGCCGCCGCCTCCTCAGCTTCCTCAGCCGCCGCCGCAGGCACAGCCGCTGCTGCCTCAGC         242          114          DEL
1373           0.0
GCTGATGAAGGCCTTCGAGTCCCTCAAGTCCTTCCAGCAGCAGCAGCAGCAGCAGCAGCAGCAGCAGCAGCAGCAGCAGCAGCAGCAGCAGCAGCAGCAGCAGCAGCAGCAGCAGCAGCAGCAGCAGCAGCAGCAGCAGCAGCAGCAGCAGCAACAGCCGCCACCGCCGCCGCCGCCGCCGCCGCCTCCTCAGCTTCCTCAGCCGCCGCCGCAGGCACAGCCGCTGCTGCCTCAGC
GCTGATGAAGGCCTTCGAGTCCCTCAAGTCCTTCC-------------------------------------------------------------------------------------------------------------------------------------------------------TCCGCCTCCTCAGCTTCCTCAGCCGCCGCCGCAGGCACAGCCGCTGCTGCCTCAGC         242          111          DEL
452            31.0
GCTGATGAAGGCCTTCGAGTCCCTCAAGTCCTTCCAGCAGCAGCAGCAGCAGCAGCAGCAGCAGCAGCAGCAGCAGCAGCAGCAGCAGCAGCAGCAGCAGCAGCAGCAGCAGCAGCAGCAGCAGCAGCAGCAGCAGCAGCAGCAGCAGCAGCAACAGCCGCCACCGCCGCCGCCGCCGCCGCCGCCTCCTCAGCTTCCTCAGCCGCCGCCGCAGGCACAGCCGCTGCTGCCTCAGC
GCTGATGAAGGCCTTCGAGTCCCTCAAGTCCTTC------------------------------CAGCAGCAGCAGCAGCAGCAGCAGCAGCAGCAGCAGCAGCAGCAGCAGCAGCAGCAGCAGCAGCAGCAGCAGCAGCAGCAGCAGCAGCAGCAACAGCCGCCACCGCCGCCGCCGCCGCCGCCGCCTCCTCAGCTTCCTCAGCCGCCGCCGCAGGCACAGCCGCTGCTGCCTCAGC         242          104          DEL
1326           0.0
GCTGATGAAGGCCTTCGAGTCCCTCAAGTCCTTCCAGCAGCAGCAGCAGCAGCAGCAGCAGCAGCAGCAGCAGCAGCAGCAGCAGCAGCAGCAGCAGCAGCAGCAGCAGCAGCAGCAGCAGCAGCAGCAGCAGCAGCAGCAGCAGCAGCAGCAACAGCCGCCACCGCCGCCGCCGCCGCCGCCGCCTCCTCAGCTTCCTCAGCCGCCGCCGCAGGCACAGCCGCTGCTGCCTCAGC
GCTGATGAAGGCCTTCGAGTC----------------------------------------------------------------------------------------------------------------------------------------------------CCACCGCCGCCGCCGCCGCCGCCGCCTCCTCAGCTTCCTCAGCCGCCGCCGCAGGCACAGCCGCTGCTGCCTCAGC         242          104          DEL
40             47.0
GCTGATGAAGGCCTTCGAGTCCCTCAAGTCCTTC----------------------CAGCAGCAGCAGCAGCAGCAGCAGCAGCAGCAGCAGCAGCAGCAGCAGCAGCAGCAGCAGCAGCAGCAGCAGCAGCAGCAGCAGCAGCAGCAGCAGCAGCAGCAGCAGCAGCAGCAGCAGCAGCAGCAACAGCCGCCACCGCCGCCGCCGCCGCCGCCGCCTCCTCAGCTTCCTCAGCCGCCGCCGCAGGCACAGCCGCTGCTGCCTCAGC
GCTGATGAAGGCCTTCGAGTCCCTCAAGTCCTTCCAAGCAGCAGCAGCAGCAGCAGCAGCAGCAGCAGCAGCAGCAGCAGCAGCAGCAGCAGCAGCAGCAGCAGCAGCAGCAGCAGCAGCAGCAGCAGCAGCAGCAGCAGCAGCAGCAGCAGCAGCAGCAGCAGCAGCAGCAGCAGCAACAGCCGCCACCGCCGCCGCCGCCGCCGCCGCCTCCTCAGCTTCCTCAGCCGCCGCCGCAGGCACAGCCGCTGCTGCCTCAGC  242          102          INS
1539           0.0
GCTGATGAAGGCCTTCGAGTCCCTCAAGTCCTTCCAGCAGCAGCAGCAGCAGCAGCAGCAGCAGCAGCAGCAGCAGCAGCAGCAGCAGCAGCAGCAGCAGCAGCAGCAGCAGCAGCAGCAGCAGCAGCAGCAGCAGCAGCAGCAGCAGCAGCAACAGCCGCCACCGCCGCCGCCGCCGCCGCCGCCTCCTCAGCTTCCTCAGCCGCCGCCGCAGGCACAGCCGCTGCTGCCTCAGC
GCTGATGAAGGCCTTCGAGTCCCTCA------------------------------------------------------------------------------------------------------------------------------------------------------------AGCCGCCGCCGCAGGCACAGCCGCTGCTGCCTCAGC         242          102          DEL
```
